# A funnel-type pipeline for accelerating discovery of tailored designer xylanosomes for efficient arabinoxylan valorization

**DOI:** 10.1101/2025.11.12.687223

**Authors:** Maria João Maurício da Fonseca, Julie Vanderstraeten, Tibo De Coninck, Silke Vlyminck, Yves Briers

**Author notes:** **Corresponding author**: Yves Briers.

## Abstract

Designer cellulosomes (DCs) are synthetic multi-enzyme complexes composed of a scaffold and docked enzymes, engineered for efficient lignocellulose degradation. However, designing these systems is challenged by an expansive design space, low protein yields, instability, and aggregation. To overcome these significant hurdles, we developed a novel, funnel-type methodological pipeline for the parallel construction, production, and analysis of designer xylanosomes (DXs), a subset of DCs targeting hemicellulose’s xylan fraction. This streamlined approach leverages VersaTile, a combinatorial DNA assembly technique, to generate an unprecedented number of modular proteins. We demonstrate the efficient construction of 96 bicatalytic dockerin-enzyme (DE) fusions. Parallel high-throughput assays, including Enzyme-Linked ImmunoSorbent Assay (ELISA) and DNA Sequencer-Assisted Fluorophore-Assisted Carbohydrate Electrophoresis (DSA-FACE), enabled rapid analysis of DE binding efficiencies and substrate preferences, facilitating selection of optimal components for downstream processing. Subsequently, we assembled 23 unique DXs and systematically evaluated their assembly capacity and hydrolytic efficiency. Our findings reveal that DX hydrolytic efficiencies are highly dependent on composition and architecture, yielding either specific arabinoxylan-oligosaccharide (AXOS) mixtures or more complete degradation to monomers. This diversity allows for tailored outcomes, leading to the identification of three superior DX variants. This innovative pipeline significantly accelerates the discovery and optimization of highly efficient multi-enzyme complexes, paving the way for more effective biomass valorization processes.

**Highlights:**

1. Funnel-type pipeline enables high-throughput testing of designer xylanosomes
2. VersaTile allows rapid assembly of an infinite number of bicatalytic dockerin-enzymes
3. High-throughput assessment of substrate preferences resulting in product profiles
4. Streamlined selection yields 23 unique DXs with diverse hydrolytic profiles
5. Proposed pipeline accelerates discovery of efficient enzyme complexes

## 1. Introduction

Cellulosomes are multi-enzyme complexes produced by specialist micro-organisms that feed on plant cell wall carbohydrates. They consist of a large backbone molecule, named the scaffoldin, that is able to incorporate a selection of different lignocellulose-degrading enzymes. The main advantage of cellulosomes compared to free enzyme systems is that multiple enzymes are brought in close proximity to each other and the substrate, ensuring a higher synergy and catalytic activity, especially for the degradation of complex substrates [1]. About 40 years ago, the first cellulosome was discovered [2–4] and since then, a vast variety has been found in a range of environments [5–7]. There has been an interest in using cellulosomes to aid in the efficient degradation of lignocellulosic biomass. However, expressing natural cellulosomes is associated with insufficient yields to be competitive for industrial purposes [8]. In addition, the majority of natural cellulosomes will not have the kinetics and intrinsic properties needed for the harsh reaction conditions to hydrolyze industrially relevant biomass streams [9]. Therefore, Bayer and colleagues proposed the construction of so-called ‘designer cellulosomes’ (DCs), which are synthetic multi-enzyme complexes, engineered to have a controlled composition, size and architecture in terms of specificity, module positioning and number of catalytical subunits [10–12]. Tailored DCs aim to provide enzyme complexes with specified substrate preferences and optimum reaction conditions for biomass degradation. Engineering these enzyme complexes permits the controlled incorporation of the needed catalytic activities depending on the target substrate and pushes the enzyme complexes to higher expression levels and optimal reaction conditions. Originally, DCs were created by selectively incorporating dockerin-containing cellulolytic enzymes, also called dockerin-containing enzymes (DEs), into functional DCs [12]. Over the years, DCs have become increasingly more complex in terms of type and number of DEs and scaffoldins [13–15]. In 2016, Davidi et al. started paving the way to combined cellulose- and lignin degradation by adding a lignin degrading enzyme to their DC [16].

Engineering high performance DCs demands the possibility and flexibility to integrate many catalytic domains with different substrate preferences. Increasing the number of integrated catalytic domains can be achieved by: (1) enlarging the scaffoldin protein with more cohesins [17] and/or (2) by creating multifunctional DEs [18–20]. Such multifunctional DEs may also bring along synergies based on enzyme-enzyme proximity, improved (thermo)stability, operating at a different pH value than the individual enzymes and/or novel substrate specificities [19,21–23]. In addition, multifunctional DEs reduce the number of proteins required for advanced cell wall degradation, offering more cost-effective protein expression. Creating inter- and intramolecular synergies with multifunctional DEs on top of the intermolecular synergies within a DC complex, offers an interesting avenue of investigation to build more elaborate and specialized multi-enzyme complexes.

The construction of multi-enzyme DCs has become a complex, often empirical, multivariate process comprising five major steps: (1) selection and delineation of protein modules to be incorporated into the DC; (2) design of the DC components (*i.e.* DEs and scaffoldin); (3) cloning of the DNA encoding the DC components; (4) expression and assembly of the DC components; and (5) activity assessment [24]. Each specific application requires a multi-directional study to develop the most capable DCs. However, technical constraints in each of the five major steps have made this a Herculean task. One of the main concerns is that a practically infinite number of multifunctional DEs can be envisioned and current methods to produce and characterize these enzymes are tedious and cumbersome. Therefore, we are in need of methods and assays to further facilitate *in vitro* DC engineering to overcome the immense design space.

In most DC studies, activity assays include the addition of the constructed DC to a defined substrate (*e.g.* cellulose, xylan, wheat straw) with the overall goal of reaching complete degradation. However, in some cases, it could be interesting to have a more detailed understanding of the intermediates formed during the reaction, allowing protein engineers to gain important insights about the cooperative mechanisms of the selected enzymes. For example, identifying the rate-limiting enzyme could help in optimizing the ratio of enzymes to be incorporated in the complex to achieve the desired product [25]. Moreover, incomplete degradation to specific oligosaccharides could also be the end goal. For instance, in the case of arabinoxylan (AX), degradation to arabinoxylan oligosaccharides (AXOS) has an economic potential because they can be commercialized as prebiotics [26].

In our previous works, we have introduced our in-house developed molecular DNA assembly method, VersaTile [27], which can be used to construct the coding sequences for a wide range of multi-modular proteins. During the past years, we have applied the technique to optimize the high-throughput DC DNA assembly process [25,28], thereby alleviating the issues encountered in the third major step of the DC construction process. However, the expression, purification and activity analysis of such a large number of complexes remained a huge effort. Indeed, these fourth and fifth major steps of the construction process are consistently realized using low-throughput methods. Typical techniques to analyze reaction products include High-Performance Anion Exchange Chromatography (HPAEC-PAD) or DNA Sequencer-Aided Fluorophore Assisted Carbohydrate Electrophoresis (DSA-FACE). HPAEC-PAD typically yields very complex data, which is not appropriate for high-throughput screening, while DSA-FACE allows for analysis of a higher number of samples in parallel in a relatively short time [29,30].

The goal of this study is twofold: (1) to generate performant designer xylanosomes (DXs) capable of complete or partial AX degradation, and (2) to showcase the possibilities, flexibility and usability of the VersaTile platform for creating multi-enzyme complexes. Given the immense diversity of enzyme, dockerin and cohesin domains in nature, it is virtually impossible to rationally predetermine the most optimal dockerin-enzyme (DE) combinations for incorporation into a final DC. Successful DC design requires the selection of DEs that exhibit: (1) adequate expression and purification yields, (2) structural stability without degradation, (3) correct binding to the matching cohesin domain, and (4) relevant catalytic properties for the intended application. To streamline this multivariate selection process, we introduce a (semi-)high-throughput funnel-type pipeline for identifying promising DEs for DC/DX assembly. This parallel approach enables the construction of hundreds of DC/DX variants per day, followed by expression, purification and evaluation of stability, substrate preference (via DSA-FACE) and dockerin-cohesin specificity (via ELISA). Applying this pipeline to 96 AX-active DEs allowed rapid assessment of the most promising candidates, ultimately yielding a manageable number of performant DXs for complete or partial AX degradation, with reduced time, effort and resources.

## 2. Material and methods

### 1.1. VersaTile cloning of AX-active enzyme sequences

Genomic DNA sequences of AX-active enzymes from *Thermobifida fusca* CIP 105594 (Institut Pasteur, Paris, France), *Clostridium thermocellum* LMG 10435 (BCCM/LMG Gent Bacteria Collection, Ghent, Belgium) and *Bifidobacterium adolescentis* LMG 10502 (ATCC15703) (BCCM/LMG Gent Bacteria Collection, Ghent, Belgium) were used for tile construction by PCR using Pfu or Phusion DNA polymerase (Thermo Fisher Scientific, Waltham (MA), USA), following the protocol as reported earlier [25]. Relevant properties of the enzymes are shown in **Supplementary File S1**. The used primers are shown in **Supplementary File S2** and were designed as explained earlier [25]. Amplicon/tile length was assessed by 1% agarose DNA gel electrophoresis and purified using the GeneJET PCR purification kit (Thermo Fisher Scientific).

Purified tiles encoding AX-active enzymes were ligated in a modified pUC19 vector (named pVTE1), in which the BsaI site within the ampicillin resistance gene was mutated. Tiles and vector digestion was done by incubating 1 µg DNA, 1x Tango buffer, 10 U XbaI and 20 U HindIII/PstI (and 1 U FastAP Thermosensitive alkaline phosphatase in case of the vector) for 2 h at 37 °C followed by enzyme inactivation at 80 °C for 20 min. Afterwards, tiles and vector were purified using the GeneJET PCR purification kit, ligated using T4 DNA ligase according to the manufacturer’s guidelines and transformed into *E. coli* TOP10 chemically competent cells. All compounds were purchased from Thermo Fisher Scientific. Putatively transformed cells were selected on lysogeny broth (LB)-agar Amp^100^ X-Gal^20^ and analyzed by colony PCR using M13 forward (5’-GTAAAACGACGGCCAGT-3’) and M13 reverse (5’-GGAAACAGCTATGACCAT-3’) primers. Colonies containing a fragment with the correct size were confirmed by Sanger sequencing (LGC genomics, Berlin, Germany).

BsaI recognition sites present in the cloned tiles were removed by inverse PCR using Phusion DNA polymerase and the primers listed in **Supplementary File S3**. After 1% agarose DNA gel electrophoresis, 40 µL amplified DNA was incubated with 10 U DpnI (Thermo Fisher Scientific) for 1 h at 37 °C followed by enzyme inactivation at 80 °C for 20 min and purification using the GeneJET PCR purification kit (Thermo Fisher Scientific). Then, 50 ng of amplicon was digested by 10 U SapI and recirculated by 3 U T4 DNA ligase, transformed into chemically competent *E. coli* TOP10 cells and selected on LB agar Amp^100^. Colonies containing a fragment with the correct size were confirmed by Sanger sequencing (LGC genomics) and served as building blocks for VersaTile assembly [27].

### 2.2 VersaTile assembly: construction of a library of bicatalytic dockerin-containing AX-active enzymes (DEs)

Entry vectors containing the AX-active tiles and dockerin tiles were combined with the pVTD3 destination vector containing a *sacB* selection marker, T7 *lac* promoter and C-terminal His_6_-tag. Dockerin tiles were already created [31]. The used dockerin and AX-active tiles are listed in **Supplementary File S4**. A number of 96 VersaTile assembly reactions were executed in parallel and contained each 100 ng pVTD3, 50 ng entry vector with AX-active or dockerin tile sequence, 10 U BsaI (Thermo Fisher Scientific), 3 U T4 DNA ligase and 2 µL 10X ligation buffer, in a final volume of 20 µL and making use of the thermal cycling conditions as described earlier [27]. Chemically competent *E. coli* BL21(DE3) CodonPlus RIL and *E. coli* Rosetta^TM^(DE3) pLysS cells were transformed with 10 µL of the VersaTile assembly mixture and selected on LB agar Kan^50^ and 5 % (w/w) sucrose and incubated overnight at 37 °C. All variants were sequenced by Sanger sequencing (LGC genomics).

### 1.2. Expression and purification of scaffoldins and bicatalytic AX-active DEs

Monovalent scaffoldins containing the sequences coding for the N-terminal cohesin n°2 of the *Clostridium thermocellum* ScaA scaffoldin, a CAZy family CBM3 (*Ct*-CBM3a) module, one cohesin (cohesin tiles are listed in **Supplementary File S5**) and a C-terminal His_6_-tag were constructed using VersaTile assembly and expressed in *E. coli* on large scale (250 mL LB with 0.5 mM IPTG + 2 mM CaCl_2_) at 16 °C for 16 h. Single colonies producing AX-active DEs were used to inoculate 500 mL Terrific Broth (TB) (24 g/L yeast extract, 20 g/L tryptone, 0.4% (v/v) glycerol, 0.017 M KH_2_PO_4_ and 0.072 M K_2_HPO_4_) in 2 L baffled flasks and expression was induced with 0.5 mM IPTG for 20 h at 16 °C.

Cells were lysed with 1x TBS buffer pH 7.4, 2 mM CaCl_2_, 10 mM imidazole, 1 mg/mL lysozyme, 1 mM PMSF, 0.1 mg/mL DNase I (Thermo Fisher Scientific) and cOmplete™, EDTA-free Protease Inhibitor cocktail (F. Hoffmann-La Roche AG, Basel, Switzerland), followed by sonication and three freeze thaw-steps. AX-active DEs and scaffoldins were purified using His GraviTrap columns (GE Healthcare, Chicago (IL), USA) according to the manufacturer’s guidelines, using 1x TBS buffer pH 7.4, 2 mM CaCl_2_ and 10 mM, 50 mM and 500 mM imidazole as equilibration, wash and elution buffers, respectively. Protein concentration was determined with Bradford assay (Bio-Rad). Elution fractions were analyzed by 12% acrylamide SDS-PAGE and Western Blot (WB). The PageRuler unstained broad range protein ladder (Thermo Fisher Scientific) and the Precision Plus Protein™ Dual Color (Bio-Rad, Hercules (CA), USA) standards were used for SDS-PAGE and WB, respectively.

### 1.3. Analysis of arabinoxylanolytic activity of DEs

Activity assays were executed in 96-well microtiter plates on 100 µL scale, containing 10 µM of (A)XOS (X_5_, XA^3^XX or XA^2+3^XX) (Megazyme, Wicklow, Ireland), purified DE (between 0.02 and 0.4 µM) and 1x TBS buffer pH 7.4, 2 mM CaCl_2_. 30 µL of mineral oil was added on top of the reaction to avoid evaporation. All reactions and substrate blanks were incubated for 22 h at 50 °C and shaking at 750 rpm and stopped by adding 1% SDS in combination with heating at 95 °C for 5 min. Reaction hydrolysates were diluted 10-fold with ultrapure water and 10 µL amounts were lyophilized. Carbohydrates present in the lyophilized fraction were derivatized with APTS by reductive amination. Afterwards, samples were quenched by diluting the reaction mixtures 200x with ultrapure water and 10 µL amounts were analyzed by DSA-FACE (3500 Genetic Analyzer, Thermo Fisher Scientific). Identification of carbohydrates after enzymatic reactions and product profiles and degradation maps were done as described earlier [29].

### 1.4. Analysis of cohesin-dockerin interactions by customized ELISA

An ELISA assay to analyze the cohesin-dockerin interactions was customized [32]. In this method, ELISA plates are coated with purified DEs. Afterwards, a CBM-containing scaffoldin is added. A primary antibody (against the CBM), is added followed by a secondary antibody able to interact with the primary antibody. A positive ELISA signal thus depends on a successful cohesin-dockerin interaction. To perform this customized ELISA assay, *Ct*-CBM3a-specific antiserum needed to be raised and characterized (see peptide sequence of *Ct*-CBM3a in **Supplementary File S6**). The production of the polyclonal rabbit anti-*Ct*-CBM3a primary antibody was ordered at Eurogentec (Seraing, Belgium). A standard 3-month, 2-animal (rabbit) immunization program was performed and the resulting serums were tested for our application.

The ELISA assay was executed with 5 nM of DE and 0.025 nM scaffoldin. ELISA plates were coated for 1 h at 37 °C or for 18 h at 4 °C with 100 µL of purified DE diluted in coating solution (0.1 M Na_2_CO_3_, pH 9). Afterwards, the coating solution was discarded, 100 µL blocking buffer (1x TBS buffer pH 7.4, 10 mM CaCl_2_, 0.05% Tween20 and 2% bovine serum albumin) was added and the plate was incubated at 37 °C for 1 h. The blocking buffer was discarded, 100 µL of the desired scaffoldin (diluted in blocking buffer) was added and the plate was incubated at 37 °C for 1 h. The plate was washed 3x with wash buffer (1x TBS buffer pH 7.4, 10 mM CaCl_2_ and 0.05% Tween20). Next, 100 µL of the polyclonal rabbit anti-*Ct*-CBM3a primary antibody was added, the plate was incubated at 37 °C for 1 h after which the plate was washed 3x with wash buffer. Then, 100 µL of the secondary antibody (horseradish peroxidase (HRP) labelled goat anti-rabbit IgG) was added and the plate was incubated at 37 °C for 1 h. The plate was washed again 4x. Finally, the interaction was detected by adding 100 µL 3,3’,5,5’-tetramethylbenzidine (TMB) substrate-chromogen. The color formation was terminated by the addition of 50 µL 1 M H_2_SO_4_. The absorbance was measured at 450 nm.

### 1.5. DX assembly and assessment of protein production

For DX assembly, the scaffoldins contained an N-terminal glutathione-S-transferase (GST) tag, *Ct*-CBM3a module, cohesin n°2 of the *Clostridium thermocellum* ScaA scaffoldin, cohesin n°1 of the *Clostridium cellulolyticum* (reclassified to *Ruminiclostridium cellulolyticum*) CipC scaffoldin and a *C*-terminal His_6_-tag. DEs and scaffoldins were expressed separately in *E. coli* BL21(DE3) CodonPlus RIL or *E. coli* Rosetta™(DE3) pLysS in duplicate 96-well plates, containing 150 µL LB Kan^50^ Cm^25^ and 5 w/v % sucrose, for 16 h at 37 °C and 700 rpm. Afterwards, deep 24-well plates containing 5 mL TB Kan^50^ Cm^25^ were inoculated with 25 µL of the pre-culture and grown at 37 °C and 650 rpm for 8.5 h. Protein production was induced with 0.5 mM IPTG and inoculated at 16 °C and 650 rpm for 16 h. Cells were harvested by centrifugation (3100 g, 30 min) and pellets were lysed using 600 µL BugBuster Master Mix (Novagen) with 1 mM PMSF and cOmplete™, EDTA-free Protease Inhibitor cocktail. Lysates were incubated at room temperature for 20 min at a slow shaking. Upon centrifugation (3100 g for 40 min), the clear supernatants were collected for further analysis through dot blot, SDS-PAGE and WB.

Non-diluted, 2x, 5x, 10x and 20x diluted (in 1x TBS buffer pH 7.4 + 2 mM CaCl_2_) clarified lysates of DEs and scaffoldins were analyzed in duplicate by dot blot. All steps were performed at room temperature. 2 µL of each sample was loaded on a nitrocellulose membrane and was left to dry for 1.5 h and was thereafter incubated with blocking buffer (30 g/L milk powder in 0.2% Tween20 in 1x PBS buffer with 0.02% NaN_3_) for 1 h. His_6_-tagged proteins were detected using consecutively Penta-His antibody (Qiagen, Hilden, Germany) and goat anti-mouse IgG human ads-HRP polyclonal (ImTec Diagnostics, Antwerp, Belgium), for 30 min each. In between antibody incubations, the membrane was washed 3×15 minutes with 1x PBS with 0.2% Tween20. Finally, the membrane was washed with demineralized water and incubated with 1-Step Ultra TMB-Blotting solution (Thermo Fisher Scientific). Color development was stopped by rinsing membrane with demineralized water and the blot was analyzed by ImageJ. In parallel, 10 and 8 µL of 2x, 4x and 200x diluted (in 1x TBS buffer pH 7.4 + 2 mM CaCl_2_) clarified DE/scaffoldin lysates were analyzed on 8% acrylamide SDS-PAGE and WB, respectively. The Roti®Mark (Carl Roth, Karlsruhe, Germany) and the Precision Plus Protein™ Dual Color (Bio-Rad) standards were used for SDS-PAGE and WB, respectively.

DXs were assembled in 24-well plates by combining DE lysate, scaffoldin lysate and 1x TBS buffer pH 7.4, 2 mM CaCl_2_, to a final volume of 500 µL, depending on the dot blot and WB analyses. The mixtures were incubated at 37°C for 1.5 h at 800 rpm. Controls containing only DE or scaffoldin were also performed. The DX complexes and controls were then incubated with 200 µL washed (1 mL 1x TBS buffer pH 7.4, 2 mM CaCl_2_) glutathione Sepharose® 4B beads (Cytiva, Washington D.C., USA) in a deep 96-well plate at room temperature and shaking at 1000 rpm for 30 min. The GST beads were collected through centrifugation (500 g, 5 min), the flow-through was removed and the beads were washed 3x with 1 mL 1x TBS buffer pH 7.4, 2 mM CaCl_2_. The DX complexes were eluted by adding 3x 100 µL 50 mM Tris-HCl pH 8, 20 mM reduced glutathione and 2 mM CaCl_2_. Upon each addition of elution buffer, samples were incubated at room temperature for 10 min at 700 rpm followed by centrifugation (500 g, 5 min) and removal of supernatant. Finally, 8 µL of flow-through (FT), washing samples and elution samples were analyzed by 8% polyacrylamide SDS-PAGE.

### 1.6. Parallel activity measurement of DXs

The activity of the newly constructed DXs was assessed by the DNS (3,5-dinitrosalicylic acid) assay [33], the D-xylose and L-arabinose/D-galactose assay kits (Megazyme) and DSA-FACE. Enzymatic reactions were performed in parallel in 96-well plates by adding 50 µL of purified DXs to 900 µL 1% (w/v) high-viscosity arabinoxylan (HV-AX) (Megazyme) dissolved in 1x TBS buffer pH 7.4, 2 mM CaCl_2_. Samples were incubated at 50 °C and 850 rpm. Samples of 50 µL were collected after 2, 5, 15, 25, 40, 50, 55, 60, 65, 70 and 75 min and incubated with 50 µL DNS reagent (0.25% DNS, 10% potassium sodium tartrate, 0.05% phenol and 0.1% potassium disulfide, in 0.4% NaOH). Collected samples were incubated for 10 min at 100 °C and the absorbance at 540 nm was measured. A xylose calibration curve between 0.5-8 mM was used to determine the amounts of reducing sugars released. Samples taken after 15, 40, 55, 70, 840 and 1260 min were also analyzed with the Megazyme kits, using half area UV microplates. A xylose and arabinose standard curve between 4-449 µM and 18-359 µM, respectively, was used. For DSA-FACE analysis, 2 µL of each hydrolysate was freeze-dried and labelled with APTS as mentioned earlier [30]. Afterwards, hydrolysates upon 70 min reaction were diluted 2000x with ultrapure water and hydrolysates upon 14 and 21 hours of reaction were diluted 10000x with ultrapure water. 15 µL hydrolysates together with 1 nM (A)XOS standards and a substrate blank were analyzed by DSA-FACE as described earlier [30].

## 3. Results and discussion

### 3.1 Methodological pipeline for the rational choice of bicatalytic DEs to be used in a DX

The choice of DEs has far-reaching consequences for the stability and catalytical properties of the final DX, and will determine if complete or partial AX degradation will be achieved. It is widely known that DE design must be carried out carefully, since fusion of the dockerin and the enzyme module is not always straightforward [28,34–38], for instance due to formation of SDS-resistant aggregates by exposure of hydrophobic regions that otherwise mediate non-covalent dockerin-cohesin interactions. Furthermore, the positioning of the protein modules exerts a considerable effect on the activity of the final DC [28].

The employed methodological pipeline, making use of the VersaTile platform [27], yielded a library of 96 sequences encoding bicatalytic, AX-active DEs. Afterwards, expression, purification and desalting were performed in 24- and 96-well plates. Protein analysis using SDS-PAGE and WB allowed assessment of (un)successful expression and purity. Afterwards, the ability to bind cognate cohesins was investigated for each purified DE. Finally, the activity of the 96 bicatalytic DEs was tested on AXOS to screen for activity and substrate preference, using DSA-FACE with a total lead time of 16-18 hours [29].

### 1.7. Construction of a library of 96 bicatalytic AX-active DEs

#### 1.7.1. VersaTile cloning

The coding sequences encoding AX-active enzymes were made compatible with the VersaTile platform [27] for different positions in the modular assembly, as defined by their position tags (PT_x_-PT_x+1_ with x referring to the position in the modular assembly) (**Figure 1**). Tiles for endo-1,4-β-xylanases from *Thermobifida fusca* (*i.e. Tf*-Xyl10A, *Tf*-Xyl10B, *Tf*-Xyl11A), a β-xylosidase (*i.e. Ba*-XylC), an exo-oligoxylanase (*i.e. Ba*-RexA) and α-L-arabinofuranosidases from *Bifidobacterium adolescentis* (*i.e. Ba*-AbfA and *Ba*-Axhd3) were constructed. The properties and characteristics of the AX-active enzymes converted to tiles are shown in **Supplementary File S1**. The xylanases differed in terms of substrate recognition and preference [39–41] but work synergistically for complete xylan degradation. The exo-xylosidases act on either reducing or non-reducing xylose units and prefer different substrates [41]. The arabinofuranosidases act on either mono-substituted or bi-substituted xylose residues. To our knowledge, this is the first time that β-xylosidases and α-L-arabinofuranosidases from *B. adolescentis* were applied in a DC context.

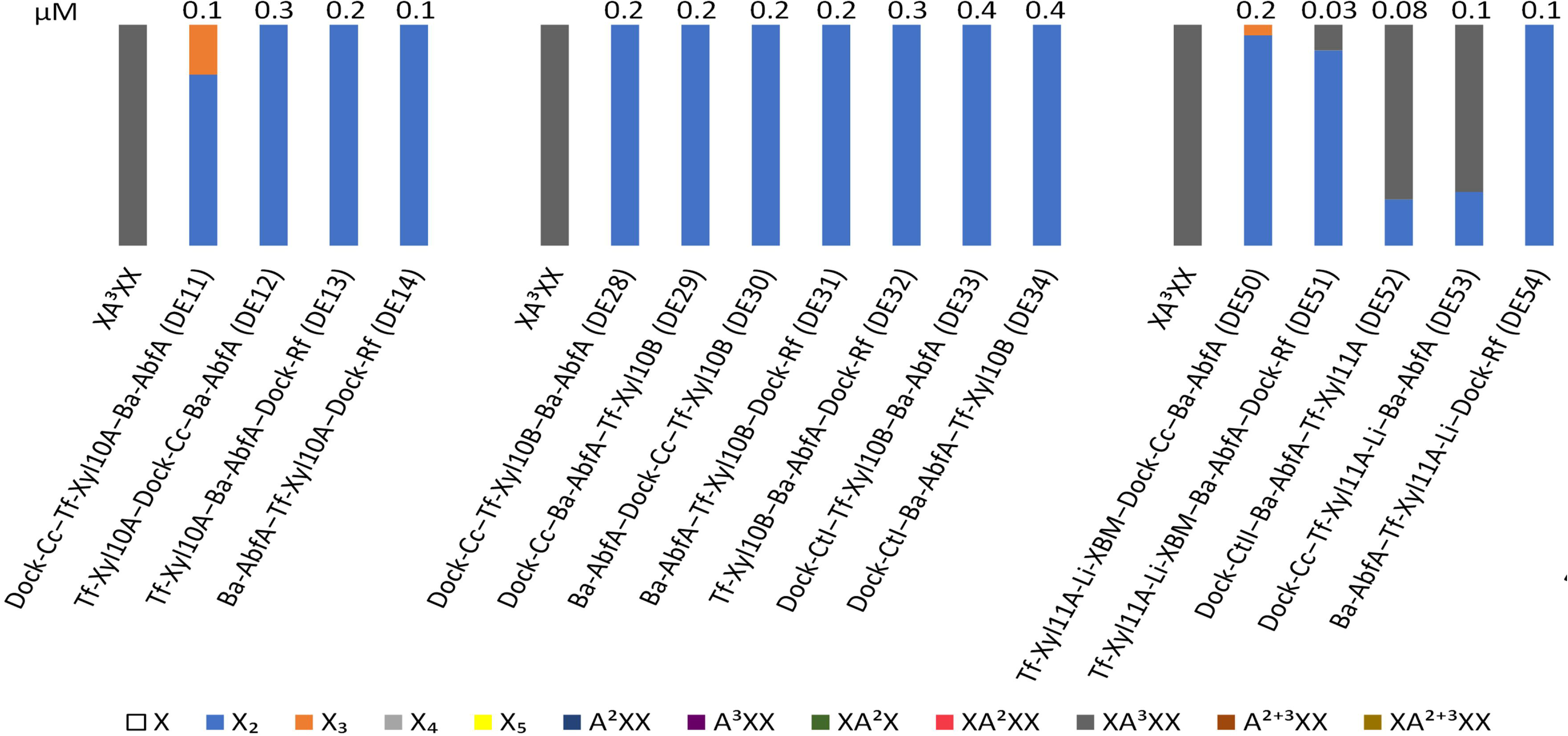

The entire sequences, except signal peptide and stop codon, encoding *Tf*-Xyl10B, *Ba*-XylC, *Ba-*RexA, *Ba-*AbfA and *Ba*-Axhd3 were amplified (**Supplementary File S2**). Tiles encoding *Tf*-Xyl10A were designed to contain solely the N-terminal catalytic domain and not the C-terminal linker and CBM. Three variants of *Tf*-Xyl11A were constructed covering either the catalytic domain (*Tf*-Xyl11A), or the catalytic domain and the natural linker present between the catalytic domain and the xylan-binding domain (XBM) (*Tf*-Xyl11A-Li), or the catalytic domain, linker and the XBM (*Tf*-Xyl11A-Li-XBM). The XBM of *Tf*-Xyl11A belongs to the CBM2 CAZy family and enhances the enzymatic activity when integrated in the DE [13]. Domain delineation was based on Pfam annotation and tertiary structure modelling using Phyre² [42], taking into account the typical linker composition of *T. fusca* xylanase [39]. ColabFold analysis confirmed that the predicted tertiary structure was not disturbed by the deletion of linker regions/CBMs [43], and that *Tf*-Xyl10A and *Tf*-Xyl11A retained a small linker of eight and seven amino acids out of the predicted 36 and 25 long linkers, respectively.

Linker length and N- or C-terminal orientation of the DE building blocks are important parameters to take into account when designing DEs. Earlier it was demonstrated that varying the dockerin-enzyme linker from 9-22 amino acid residues had little to no effect on enzymatic activity of a *T. fusca* Cel5A domain, while the N- or C-terminal position of the Cel5A domain demonstrated measurable effects on the enzymatic activity [44]. However, in contrast, a hyper-thermostable DC containing enzymes from *Caldicellulosiruptor bescii* revealed that a longer dockerin-enzyme linker (64-73 residues instead of 7 or less residues) resulted in enhanced GH9 activity [45]. Concordantly, Vazana and colleagues reported that DCs built with long linkers exhibited higher hydrolytic activity compared to those with short or no linkers [46]. The differential effect of linker length on enzymatic activity reinforces the need to screen for linker length, as observed by Vanderstraeten and colleagues [25,28]. Recently, the iMARS framework for rational multienzyme architecture design was introduced, which could be adapted for DEs and DCs, especially for optimizing linker length and relative domain orientations [47]. In addition, it might be worthwhile to inspire DC design in terms of linker length and module orientation by naturally occurring cellulosomes [24].

The dockerin tiles were selected from *Clostridium thermocellum*, *Ruminococcus flavefaciens* and *Clostridium cellulolyticum* (Dock-*Ct*I, Dock-*Ct*II, Dock-*Rf* and Dock-*Cc*). Dockerin-cohesin pairs were chosen and designed based on earlier work [31]. The available tile repository for AX-active enzymes and dockerin tiles is given in **Supplementary File S4**.

#### 1.7.2. VersaTile assembly

VersaTile (VT) assembly encompasses the assembly of tiles in a specified order into a destination vector [27] (**Figure 1**). Every assembly reaction contained one dockerin tile and two AX-active tiles to achieve the coding sequence for a bicatalytic DE. Using eight AX-active tiles and four dockerin tiles, no less than 464 combinatorial DEs can theoretically be constructed (**Supplementary File S7**). It is practically impossible to estimate in advance the better dockerin-enzyme combinations or conformations of the DEs to be incorporated in the DXs. Instead of testing all 464 possible DEs, it was chosen to test a representative, non-redundant and functionally diverse subset of DEs. We selected 96 variants to be rationally assembled (**Table 1**) in a single step with short hands-on time (∼1.3 min per construct). Depending on the available tiles per position, different configurations were tested with the dockerin at the C-terminus, N-terminus or internal position. From all possible combinations, many were removed that would give overlapping or very similar substrate specificities, thereby preventing DEs that would result in practically identical product profiles. Each bicatalytic DE was designed to have two different catalytic domains with non-overlapping target linkages in AX (**Table 1**), so that each DE, when assembled in the final DX, has the potential for synergistic action rather than duplication of enzymatic activity. Catalytic domains from different organisms and GH families (**Supplementary File S1**) were combined to increase diversity in substrate preferences, stability and pH/temperature optima. Furthermore, a selection of 96 DEs allow for screening in 96-well plates. Upon sequencing, all variants showed the correct tile combination.

**Table 1.**
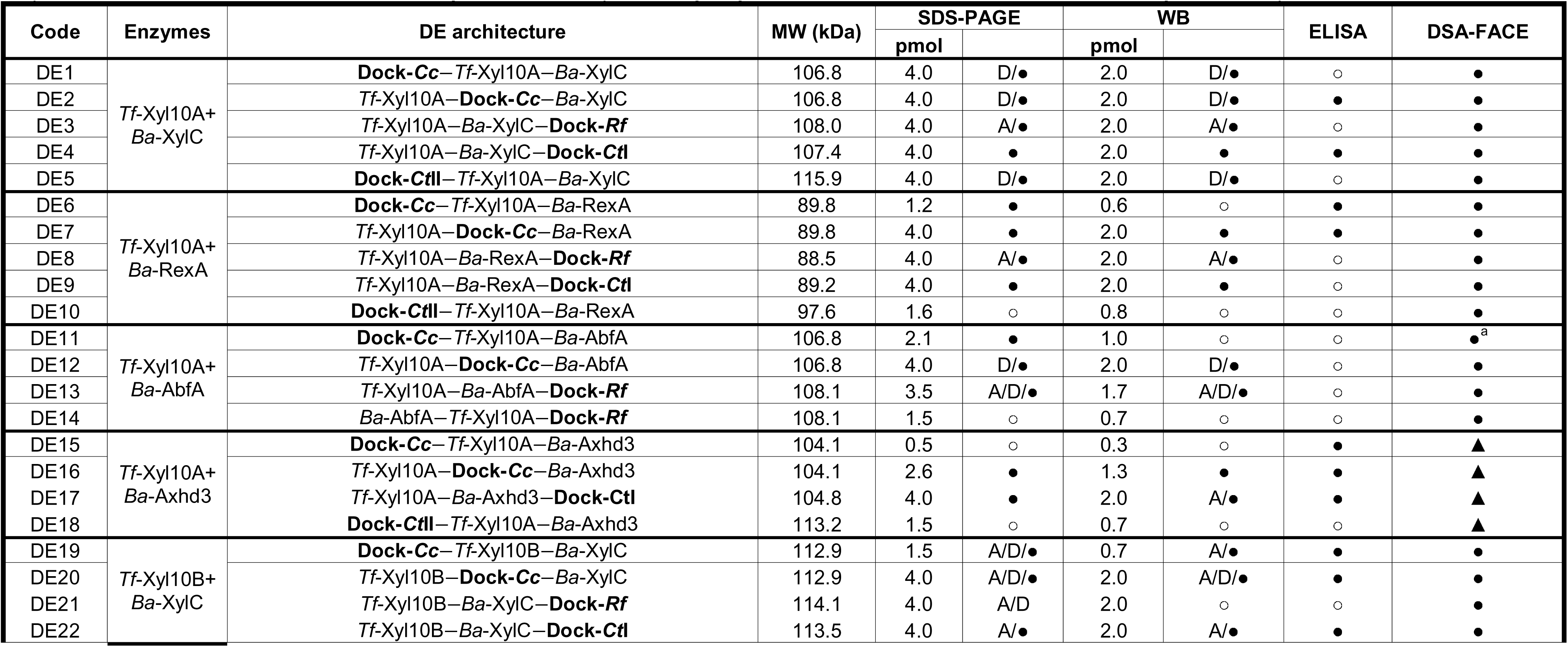
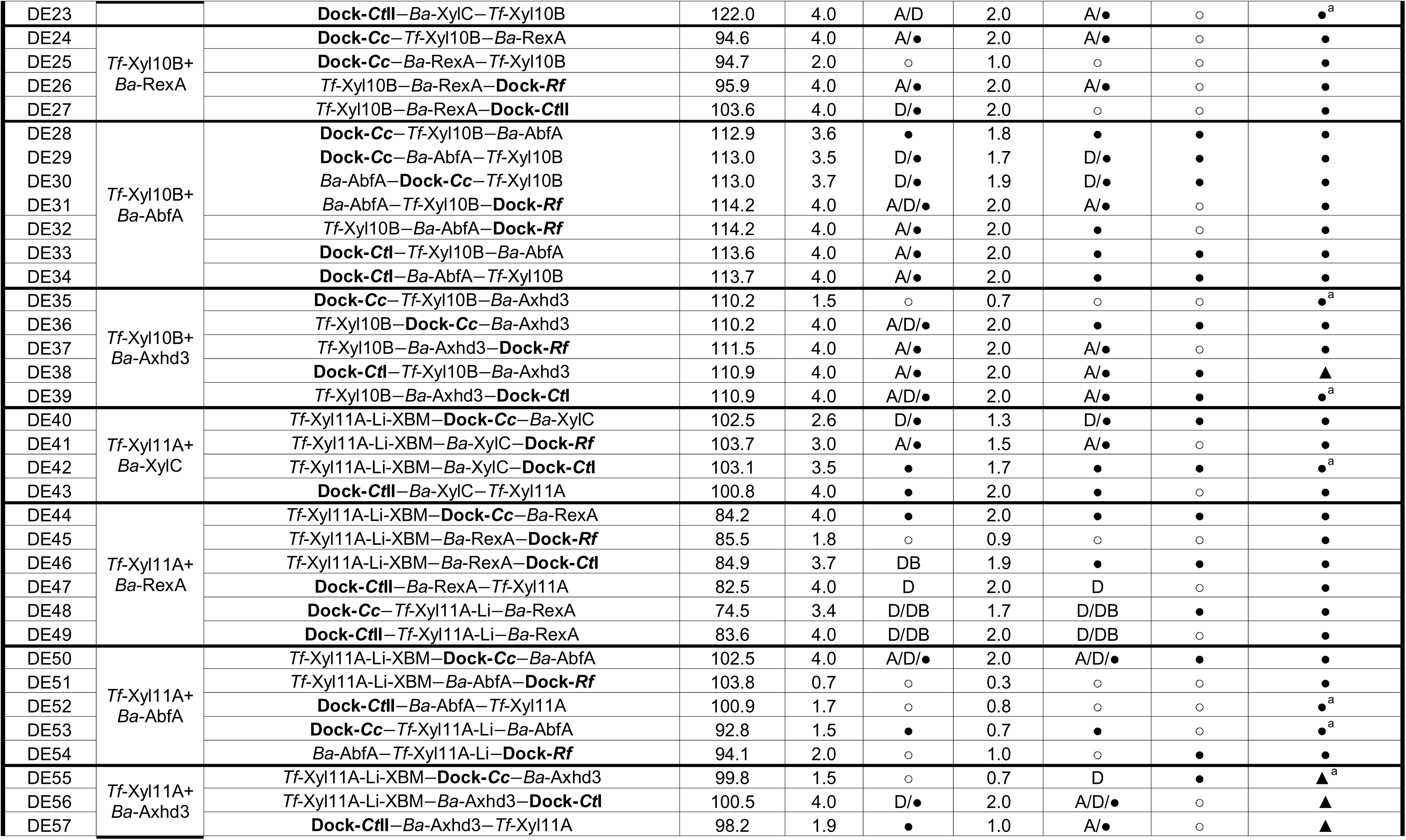
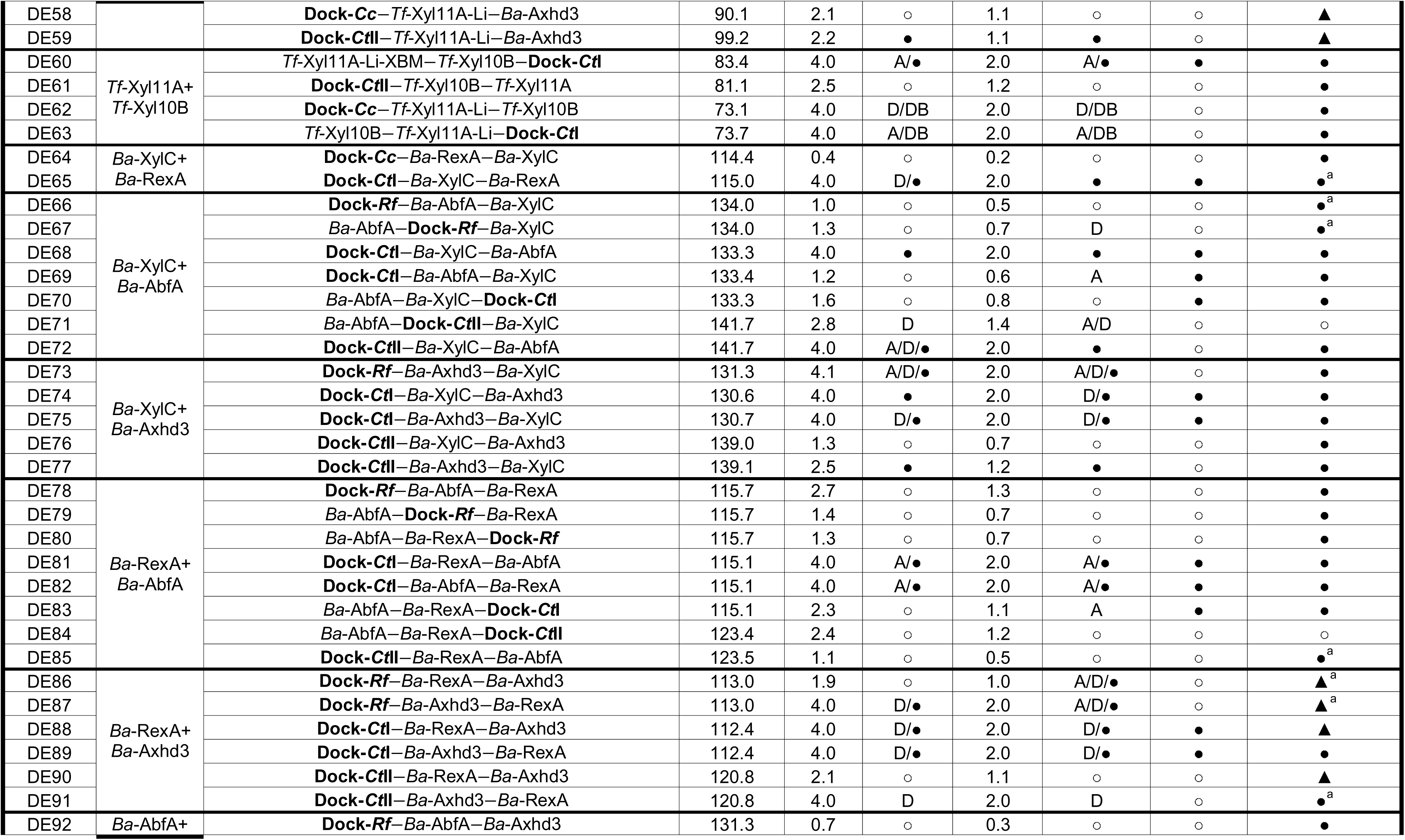
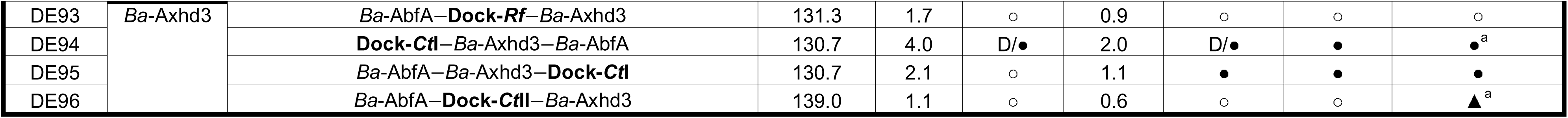
Analysis of 96 bicatalytic AX-active DEs, constructed using the VersaTile platform. . The ‘Code’ column indicates the short name for each DE constructed. In the column ‘Enzymes’ the original enzymes of the catalytic domains incorporated in each DE are indicated. Origin and details for each enzyme can be found in **Supplementary File S1**. In the ‘DE architecture’ column the full name of the newly constructed DE is written. The DE name contains the abbreviations of each domain written from *N*- to *C*-terminus. The origin and details for each dockerin used can be found in **Supplementary File S5**. Molecular weight (MW) of the constructs was determined by the Expasy ProtParam tool. For each DE, the amount of protein applied on SDS-PAGE and WB to assess expression/purification is given in pmol (**Table 2**). Overview of the results of the expression/purification checked by SDS-PAGE and WB is summarized as ‘•’ and ‘○’ if a protein band or no protein band is observed at the expected MW. In addition, ‘D’ for when one or more bands at a lower MW than the expected, likely corresponding to degradation products, are observed, ‘A’ for when one or more bands at a higher MW than the expected, likely corresponding to SDS-resistant aggregation products, are observed and ‘DB’ for when double bands at very similar MW were observed at the expected MW (**Table 2**). ELISA results are also summarized with a ‘•’ when the respective DE binds solely to the cohesin from the same organism as the dockerin or ‘○’ when no or non-specific binding is observed. Original data of the ELISA assays is included in **Supplementary File S9**. The last column corresponds to the DSA-FACE results showing whether or not there was substrate degradation, ‘•’ and ‘○’ stand for when substrate was degraded or not, respectively, and ‘▲’ when a different substrate preference than the one expected for the modules of the DEs is detected by DSA-FACE. ^a^ Spikes for hydrolysates of these DEs were done to confirm identity of reaction products.

### 1.8. Expression and purification of AX-active DEs

Upon parallel expression and purification, all 96 purified fractions were analyzed by SDS-PAGE and WB (**Supplementary File S8**). More than 67% of the bicatalytic DEs was expressed and purified and visible on SDS-PAGE (65/96) and WB (67/96). A number of the 17/96 (SDS-PAGE) and 20/96 (WB) proteins appeared as a single band at the correct molecular weight. However, many proteins suffered from degradation or aggregation (**Table 2**). There was no specific trend observed in terms of type of dockerin or position of the dockerin influencing the SDS-PAGE/WB band pattern. It is once more confirmed that production and purification of DEs *in vitro* proves to be challenging and highly unpredictable, due to the recurrent degradation and/or aggregation phenomena [48,49].

**Table 2.**
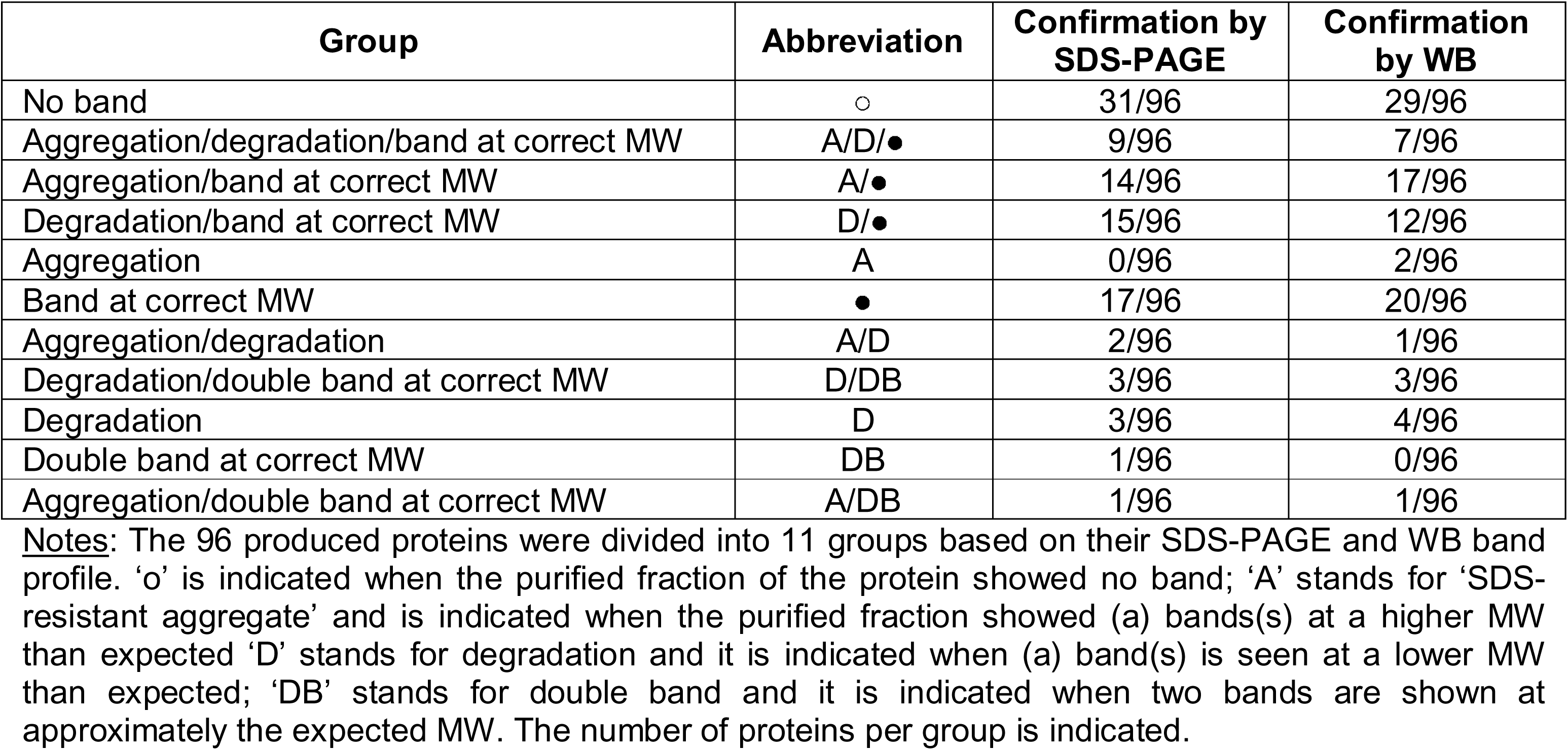
Summary of the SDS-PAGE and Western Blot results when 96 AX-active Des were produced in *E. coli* and purified, in parallel.

| Group | Abbreviation | Confirmation by SDS-PAGE | Confirmation by WB |
| --- | --- | --- | --- |
| No band | ○ | 31/96 | 29/96 |
| Aggregation/degradation/band at correct MW | A/D/● | 9/96 | 7/96 |
| Aggregation/band at correct MW | A/● | 14/96 | 17/96 |
| Degradation/band at correct MW | D/● | 15/96 | 12/96 |
| Aggregation | A | 0/96 | 2/96 |
| Band at correct MW | ● | 17/96 | 20/96 |
| Aggregation/degradation | A/D | 2/96 | 1/96 |
| Degradation/double band at correct MW | D/DB | 3/96 | 3/96 |
| Degradation | D | 3/96 | 4/96 |
| Double band at correct MW | DB | 1/96 | 0/96 |
| Aggregation/double band at correct MW | A/DB | 1/96 | 1/96 |

**Notes:** The 96 produced proteins were divided into 11 groups based on their SDS-PAGE and WB band profile. '○' is indicated when the purified fraction of the protein showed no band; 'A' stands for 'SDS-resistant aggregate' and is indicated when the purified fraction showed (a) band(s) at a higher MW than expected; 'D' stands for degradation and it is indicated when (a) band(s) is seen at a lower MW than expected; 'DB' stands for double band and it is indicated when two bands are shown at approximately the expected MW. The number of proteins per group is indicated.

### 1.9. Cohesin-dockerin binding

The 96 DEs were tested for their capacity to bind specifically to their cognate cohesin, using a modified ELISA assay [32]. An overview of the results of the ELISA assays is shown in **Table 1** and in **Supplementary File S9**. A large diversity in binding specificity was observed amongst the DEs, with 37% of the 96 DEs specifically binding to its cognate cohesin (*i.e.* DEs containing Dock-*Ct*I and Dock-*Cc*), whereas others showed no binding or bound to multiple cohesins (*i.e.* DEs containing Dock-*Ct*II and Dock-*Rf*). Most of the DEs containing Dock-*Ct*I (27% of the 96 DEs) and Dock-*Cc* (27% of the 96 DEs) specifically bound to the corresponding scaffoldin. However, some DEs containing Dock-*Ct*I and Dock-*Cc* could bind to both Coh-*Ct*I and Coh-*Cc* (*f.i.* DE1, DE56, DE62 and DE63). Indeed, Barak et al. reported cross-specificity of *C. thermocellum* dockerins and *C. cellulolyticum* cohesins [32]. These observations show the need to evaluate the potential of each variant in a multi-enzyme complex.

### 1.10. Product profiles and degradation maps of the library

The application of DSA-FACE offers substantial advantages for the characterization of degradation products generated by arabinoxylanolytic enzymes acting on (A)XOS. DSA-FACE enables detection of (oligo)saccharide products at the femtomole-picomole level, including minor or transient products that often escape detection by alternative methods such as TLC, HPAEC-PAD or MALDI-TOF [50–52]. DSA-FACE distinguishes (oligo)saccharides differing by a single sugar or branching pattern, providing structural insights into side chain arabinosylation and backbone cleavage and allows differentiation between isomeric products. Unlike HPAEC-PAD, which yields chromatograms requiring careful interpretation of elution profiles, often complicated by co-eluting isomers, DSA-FACE produces electrophoretic banding patterns that are easier to interpret visually, with each band corresponding to a distinct (oligo)saccharide. Therefore, the production profiles obtained by using DSA-FACE serve as molecular fingerprints of enzyme activity, revealing substrate preferences, cleavage modes, and synergistic interactions between enzymes or DEs in the context of designer cellulosomics. DSA-FACE is especially suited for monitoring enzymatic activity on (A)XOS substrates and for time-course studies, and it enables semi-quantitative comparison of product abundances across experimental conditions [29,30].

DSA-FACE was performed with all 96 purified DE fractions using AXOS and XOS as substrates. The obtained electropherograms were converted to product profiles based on standards and spiking experiments [29,30]. Below we discuss the different subgroups of bicatalytic DEs, disclosing a remarkable variety of enzymatic activities and product profiles.

#### 1.10.1. AX backbone-active DEs show the expected specificity

Endo-xylanases *Tf*-Xyl10A, *Tf*-Xyl10B and *Tf*-Xyl11A are expected to hydrolyze the backbone of AX and/or XOS into products with a degree of polymerization (DP) ≥ 2 (*i.e.* xylobiose, X_2_). The β-xylosidase (*Ba*-XylC) should then be able to hydrolyze XOS in an exo-xylanolytic way to xylose (X). Several DE variants combining endo-xylanase and β-xylosidase activities, fully converted xylopentaose (X_5_) to X, particularly DE20, DE22, DE42 and DE43 (**Table 1**), which all contain either *Tf*-Xyl10B or *Tf*-Xyl11A as endo-xylanase. However, half of the bicatalytic DEs from this subgroup had X_2_ as end product, with a few other variants (DE5, DE40 and DE41) yielding a mixture of X_2_ and X_3_ or X_5_. In contrast, combinations with *Tf*-Xyl10A often stalled at xylobiose (X_2_), indicating it may hinder complete hydrolysis. Furthermore, variants containing *Tf*-Xyl10B and *Tf*-Xyl11A hydrolyzed X_5_ into X_2_ (**Figure 2A**).

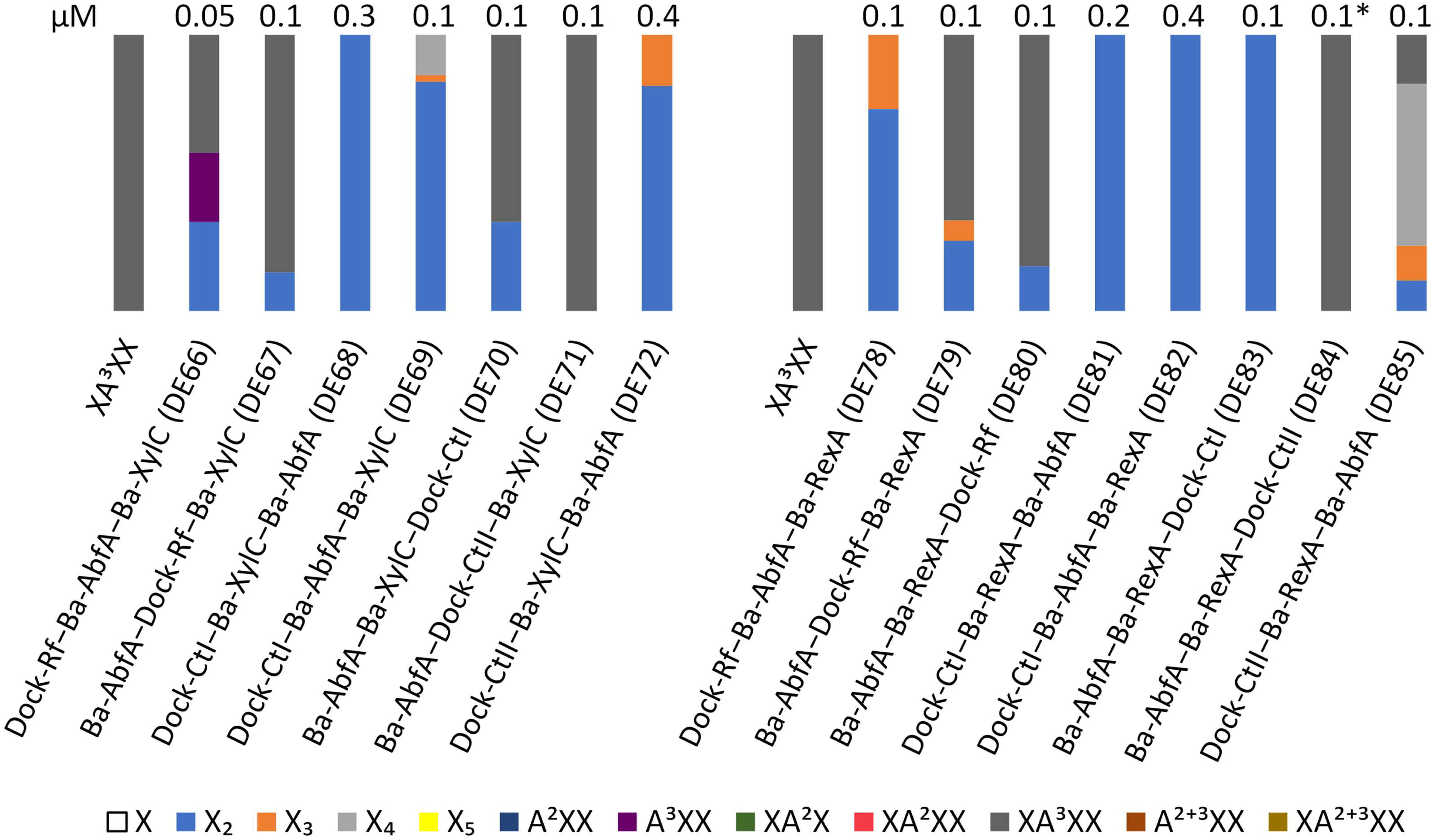

*Tf*-Xyl10A, *Tf*-Xyl10B or *Tf*-Xyl11A and the reducing-end xylose-releasing exo-oligoxylanase *Ba*-RexA combinations completely hydrolyzed X_5_ into X_2_, in most cases. DE6-10, DE24-27 and DE44-49 are active and show the expected reaction products. However, the data cannot uncover whether one or both catalytic modules are active, because of overlapping substrate preferences. When combining *Ba*-XylC and *Ba*-RexA, XOS should be completely converted to X. However, DE64 and DE65 generated mainly X_2_ (**Figure 2B**).

These observations underline the importance of evaluating the performance of each DE individually, as it is clear that the relationship between DE structure/architecture and enzymatic function is less straightforward than anticipated.

#### 1.10.2. DEs combining AX backbone-active activities with *Ba*-AbfA show the expected specificity

In line with their expected activity, DEs with an endo-xylanase and *Ba*-AbfA (*i.e.* DE12-14, DE28-34, DE54) (**Table 1**) completely hydrolyzed XA^3^XX into X_2_. *Ba*-AbfA is able to hydrolyze the *O*-3 arabinose (A) from mono-substituted X of XA^3^XX, while Xyl10A, Xyl10B or Xyl11A hydrolyze the resulting X_4_ into X_3_ and/or X_2_. The endo-xylanases can hydrolyze X_3_ further into X_2_. Exceptions were observed for DE11 and DE50, which showed X_3_ as an intermediate product, and DE51, DE52 and DE53, which showed unreacted XA^3^XX in the reaction hydrolysate (**Figure 3**).

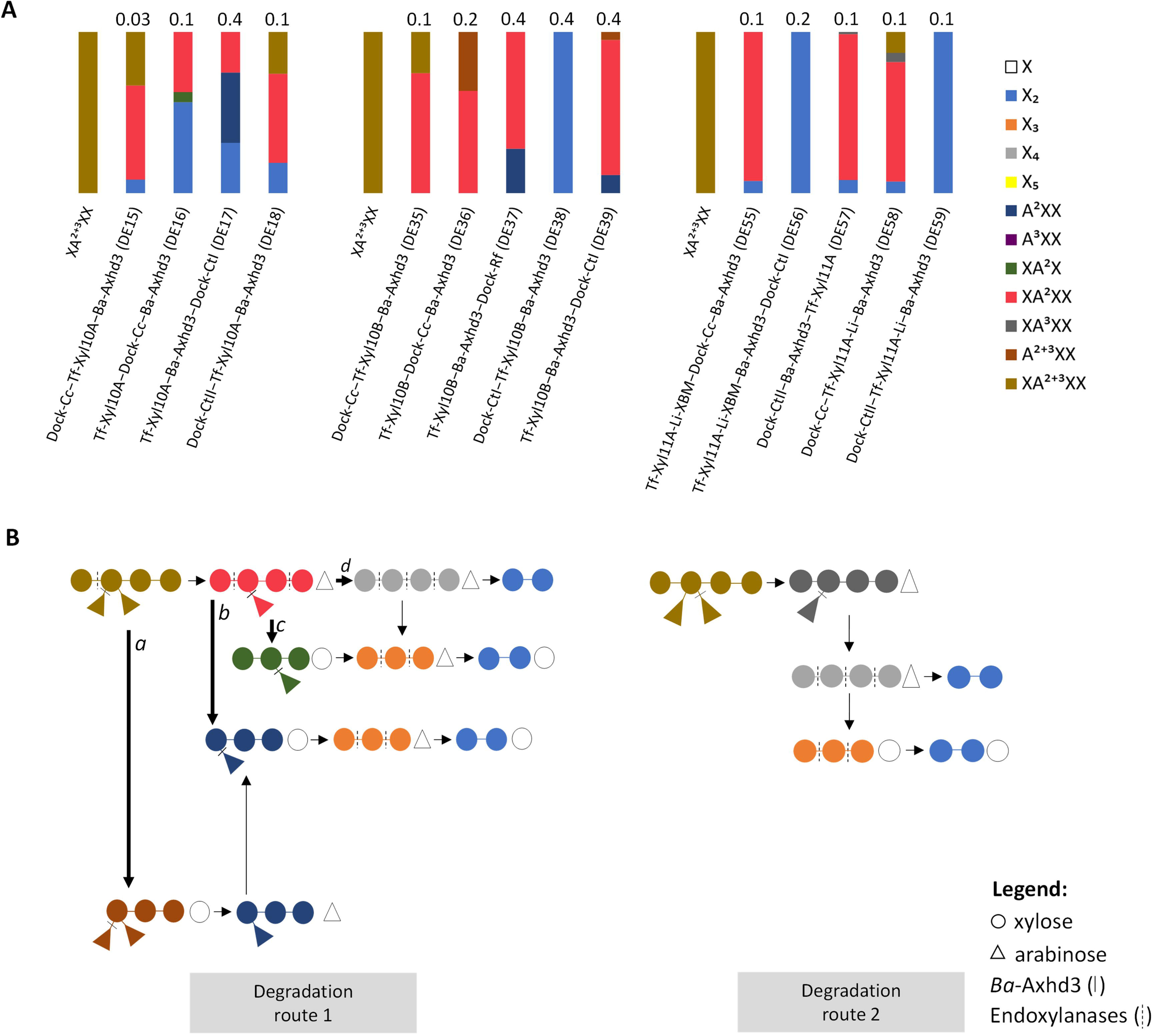

In contrast, none of the DEs with β-xylosidase (*Ba-*XylC) and α-L-arabinofuranosidase (*Ba-*AbfA) modules (DE66-72) could completely hydrolyze XA^3^XX into X. Here, it was expected that upon removal of *O*-3 A by *Ba-*AbfA, *Ba-*XylC would hydrolyze X_4_ into X. In all DEs except DE71, *Ba*-AbfA hydrolyzed the *O*-3 A residue from XA^3^XX into X_4_ and X_4_ was then hydrolyzed into the smaller XOS X_2_ and X_3_. In DE66, A^3^XX was detected in the hydrolysate showing the non-reducing activity of *Ba*-XylC on XA^3^XX. The same is seen with DEs containing *Ba-*RexA and *Ba-*AbfA catalytic modules (DE78-85), where XA^3^XX was converted into X_2_ (DE81-83) (**Figure 4**).

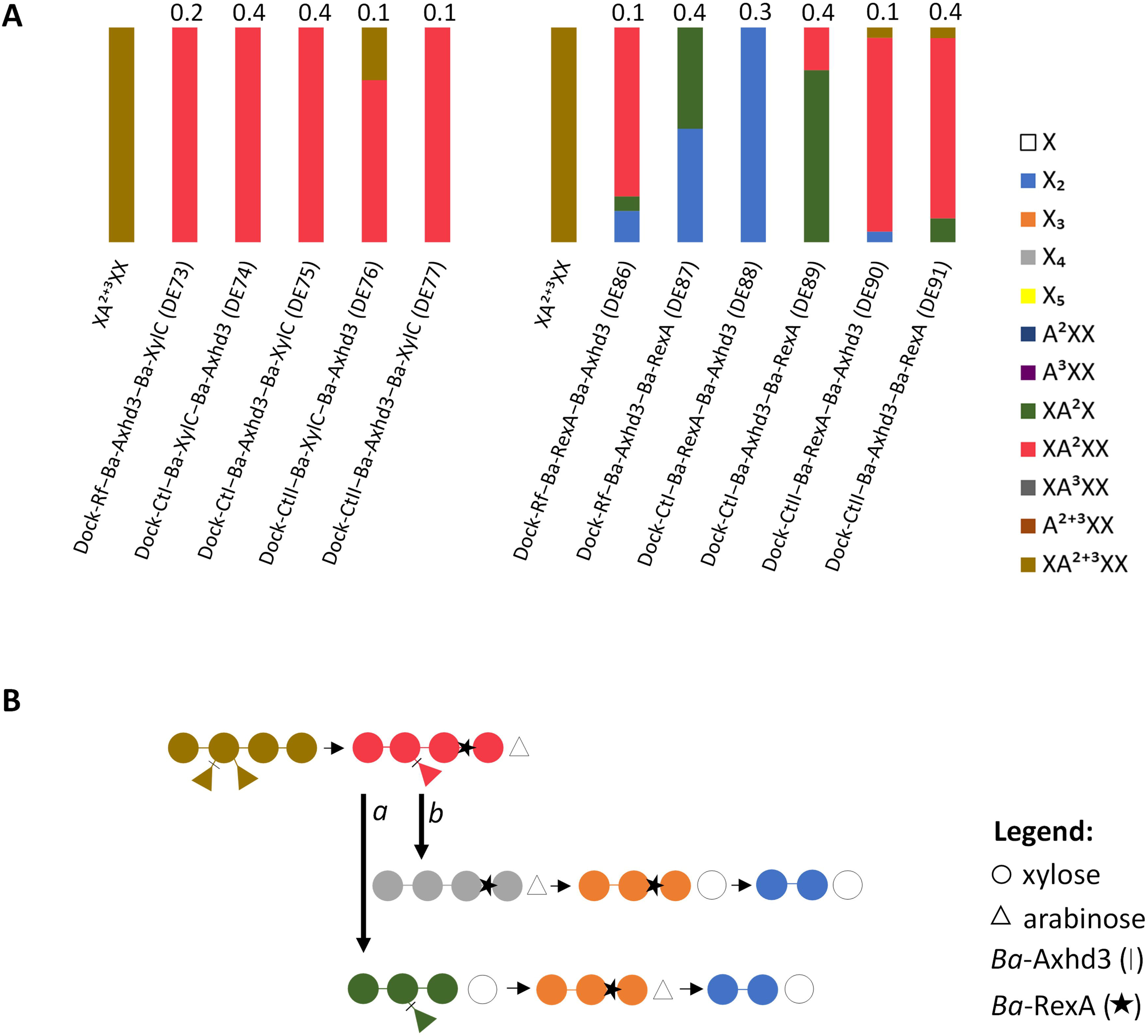

#### 1.10.3. DEs with *Ba*-Axhd3 and endo-xylanases or *Ba*-RexA act on singly and doubly **α**-L-arabinose-substituted AXOS

The α-L-arabinofuranosidase *Ba*-Axhd3 is reported to hydrolyze only *O*-3 A units from bi-substituted (*O*-2/*O*-3) X [53,54]. Therefore, enzymatic reactions with XA^2+3^XX and DEs containing *Ba*-Axhd3 as the only α-L-arabinofuranosidase are expected to yield final reaction products substituted with *O*-2 A. However, the product profiles of DEs containing *Ba*-Axhd3 and *Tf*-Xyl10A (DE15-18), *Tf*-Xyl10B (DE38) or *Tf*-Xyl11A (DE55-59), respectively, reveal besides the *O*-2 substituted oligosaccharides A^2^XX (dark blue), XA^2^X (dark green) and XA^2^XX (fuchsia), also unsubstituted X_2_, suggesting that both *O*-2 and *O*-3 A substitutions are cleaved from the bi-substituted X (**Figure 5A**). Alpha-L-arabinofuranosidases have been reported to show multiple substrate preferences depending on the linkage type and position of substituents [55–57]. This observation suggests that the *Ba*-Axhd3 catalytic module is responsible for the divergent activity seen in the bicatalytic DEs. However, it cannot be excluded that the bicatalytic architecture and corresponding intermodular interactions affect the conformation and thus specificity of each module.

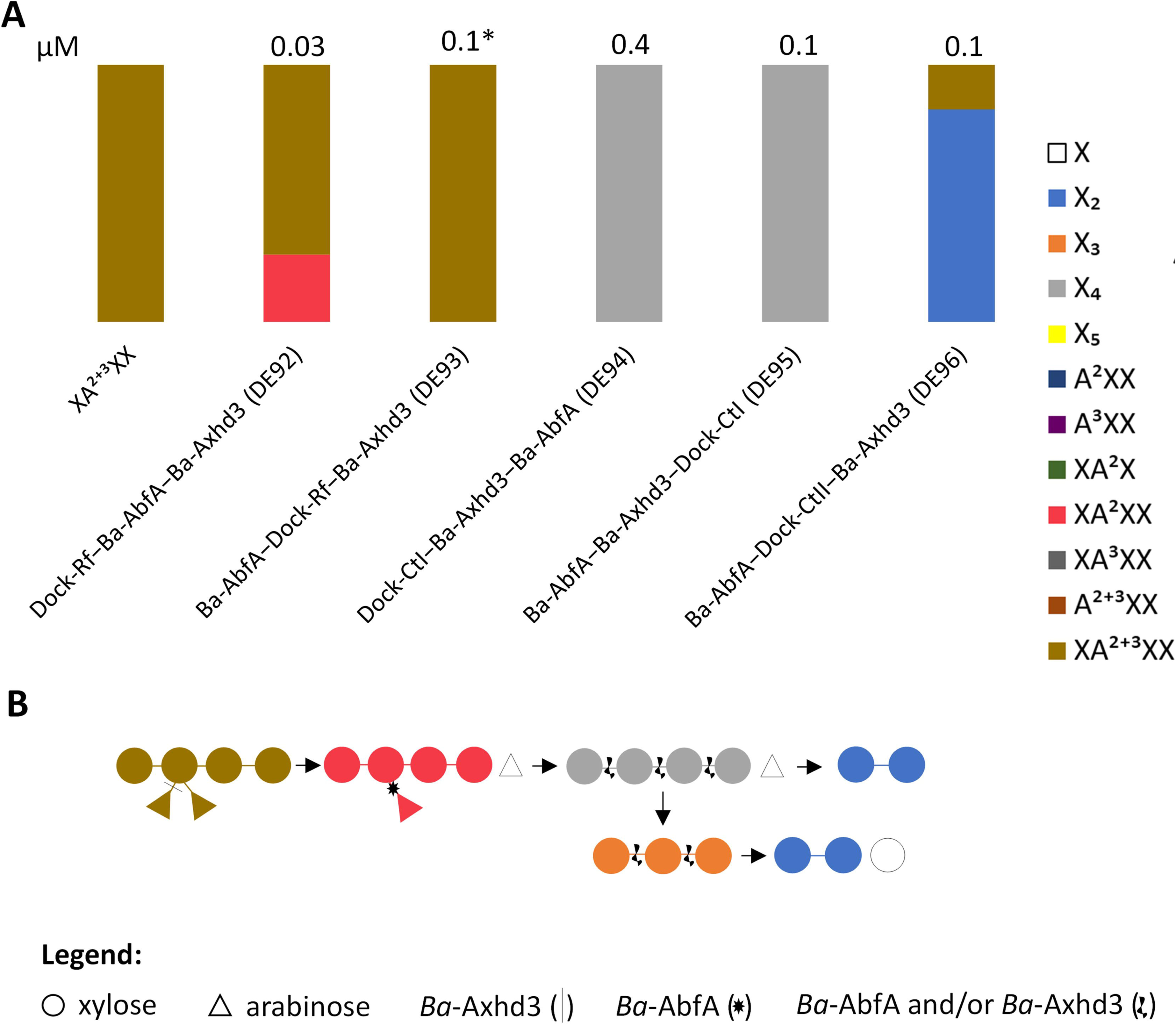

The appearance of the intermediate products A^2^XX (dark blue) and XA^2^X (dark green) in the presence of *Tf*-Xyl10A (DE16 and DE17) or *Tf*-Xyl10B (DE37 and DE39) is in accordance with their substrate preferences where GH10 endo-xylanases can tolerate A substitutions at the +1 and -2 subsites [58]. A^2+3^XX (brown) also appears in the hydrolysate of DE36, which indicates *Tf*-Xyl10B tolerates double A substitutions at the +1 subsite. Accordingly, DEs composed of *Tf*-Xyl10A/*Tf*-Xyl10B and *Ba-*Axhd3 are expected to follow degradation route 1 via 4 possible branches (*a, b*, *c*, *d*) (**Figure 5B**). DEs containing *Tf*-Xyl11A and *Ba*-Axhd3 (DE55-59) seem to follow any of the branches of degradation route 1. Noticeably, minor amounts of XA^3^XX are generated in DE57 and DE58 indicating that *Ba*-Axhd3 can also attack the *O*-2 A of a bi-substituted X as shown in degradation route 2 (**Figure 5B**).

Variants combining *Ba*-XylC and *Ba*-Axhd3 (DE73-77) generated XA^2^XX as the only end product and the XylC module did not further hydrolyze XA^2+3^XX and/or XA^2^XX (**Figure 6A**). This is in accordance with the work of Lagaert et al. who showed that XylC is hindered by A substitutions [59]. However, the XylC module of DE66 (Dock-−*Ba*-AbfA− *Ba*-XylC) seemed to hydrolyze XA XX into A XX. Whether XylC is inhibited by *O*-2 A substitutions (XA^2^XX) and/or whether it was hindered in the subset of variants tested in combination with Axhd3 (DE73-DE77) should be further evaluated. In contrast, most of *Ba*-RexA/*Ba*-Axhd3 variants (DE86-91) yield X_2_ as the final product via branch *a* and/or *b* (**Figure 6B**). The product profiles demonstrate that the outcome is not mere the theoretical reaction products of both incorporated enzymatic domains, but that modular composition and order are a major determinant as well. This underscores the usefulness of the presented parallel VersaTile- and DSA-FACE-driven pipeline, as only an empirical, combinatorial high-throughput approach is currently able to identify the best variant.

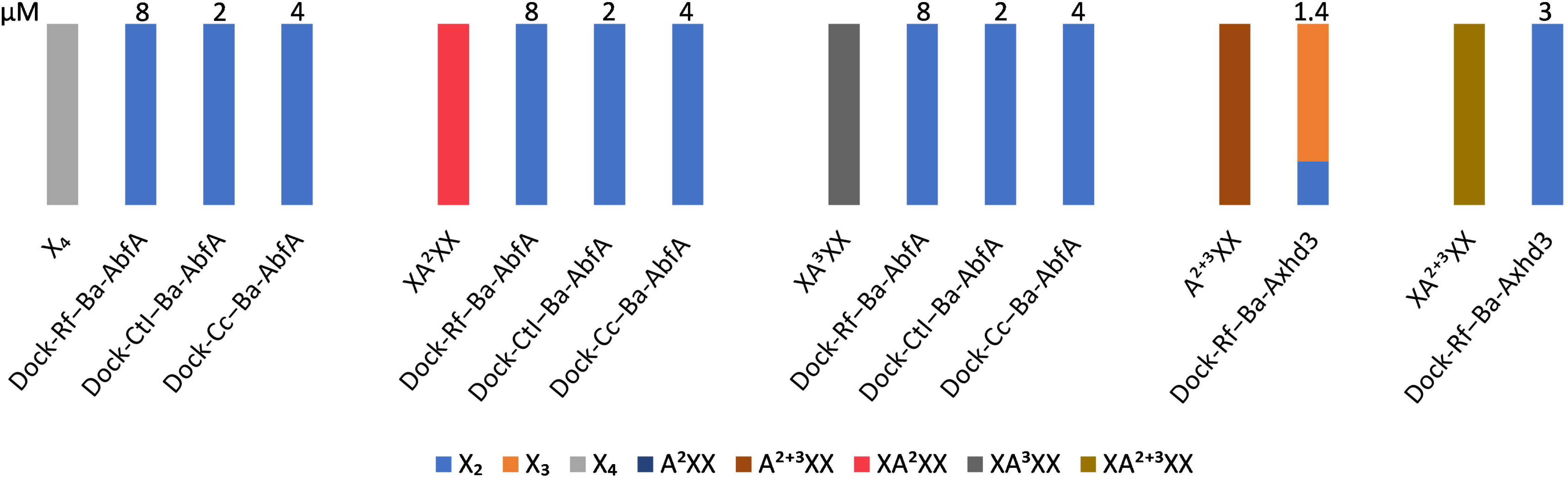

#### 1.10.4. DEs with *Ba*-AbfA and *Ba*-Axhd3 act on the XA^3^XX and XA^2+3^XX backbone

When performing enzymatic reactions with DEs with *Ba*-AbfA and *Ba*-Axhd3 and XA^2+3^XX (DE92-96), one would expect to obtain X_4_ as the reaction product, as *Ba*-Axhd3 is expected to hydrolyze the *O*-3 A monomer from XA^2+3^XX, generating XA^2^XX, and *Ba*-AbfA is able to hydrolyze *O*-2 or *O*-3 A monomers from xyloses. As such, XA^2^XX would be hydrolyzed into X_4._ This is the case for DE94 and DE95 (**Figure 7A**). DE93 is inactive (*i.e.* most likely not expressed since no band on SDS-PAGE and WB was seen) and DE92 only partially converts XA^2+3^XX into XA^2^XX, which may be explained by the different configuration/tile composition or the lower purification yields, limiting the concentration used in the DSA-FACE analysis. Surprisingly, DE96 generated X_2_, meaning that *Ba*-AbfA, *Ba*-Axhd3 or both are able to hydrolyze the xylan backbone as well, likely in an exo-xylanolytic way (Error! Reference source not found.).

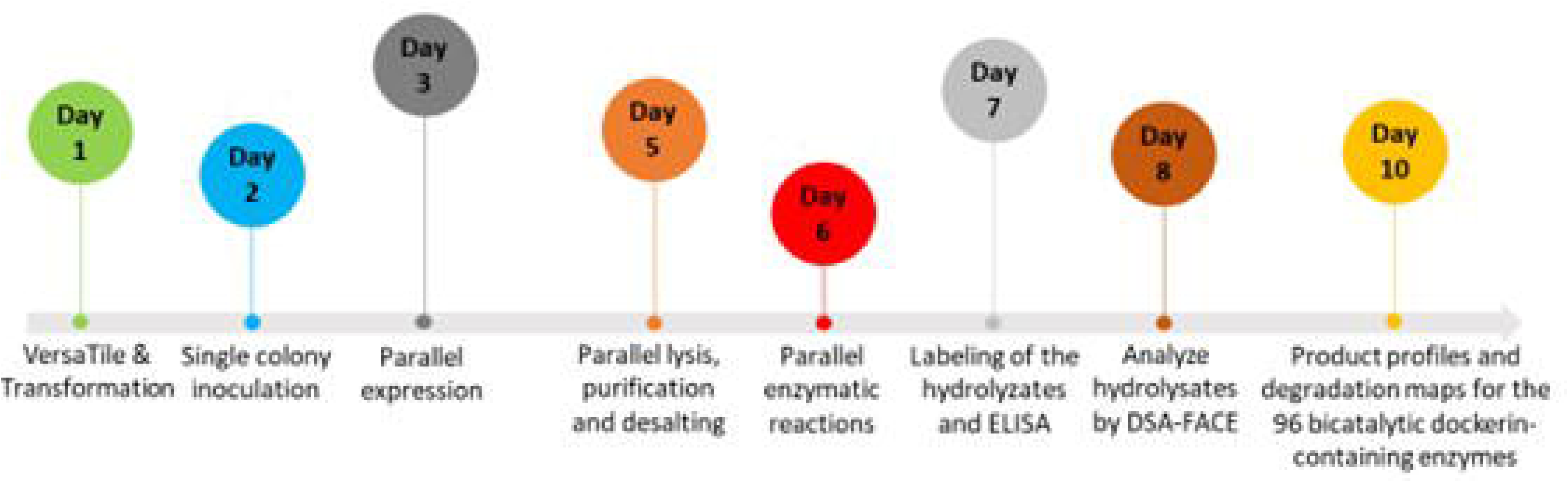

Many efforts have been made to understand the mode of action of members of the GH43 family (*i.e. Ba*-AbfA and *Ba*-Axhd3 are GH43 members) and to understand the variety of activities and substrate specificities [60]. Interestingly, many GH43 enzymes are bifunctional or even trifunctional [55]. Furthermore, the activity and specificity of GH43 enzymes is significantly affected by the used enzyme concentration as well as the presence of bivalent ions such as Ca^2+^ [60–63].

Enzymatic reactions with monocatalytic DEs containing a dockerin module at the *N*-terminus and a *Ba*-AbfA or *Ba*-Axhd3 catalytic domain at the *C*-terminus were also performed with X_4_, XA^2^XX, XA^3^XX, A^2+3^XX and XA^2+3^XX (**Figure 8**). Unexpectedly, like the bicatalytic DE96, all monocatalytic DEs tested hydrolyzed the analyzed (A)XOS into X_2_. This means that DE96 and monocatalytic DEs present AXH-m2,3 and AXH-d2,3 activities and endo- and/or exo-xylanase activity [64]. Enzymatic reactions with increasing concentrations of *Ba*-AbfA and *Ba*-Axhd3 with X_4_, A^2^XX, XA^3^XX, A^2+3^XX and XA^2+3^XX, reveal that both *Ba*-AbfA and *Ba*-Axhd3 are able to hydrolyze both *O*-2- and *O*-3-A substitutions from mono- and bi-substituted xyloses [30].

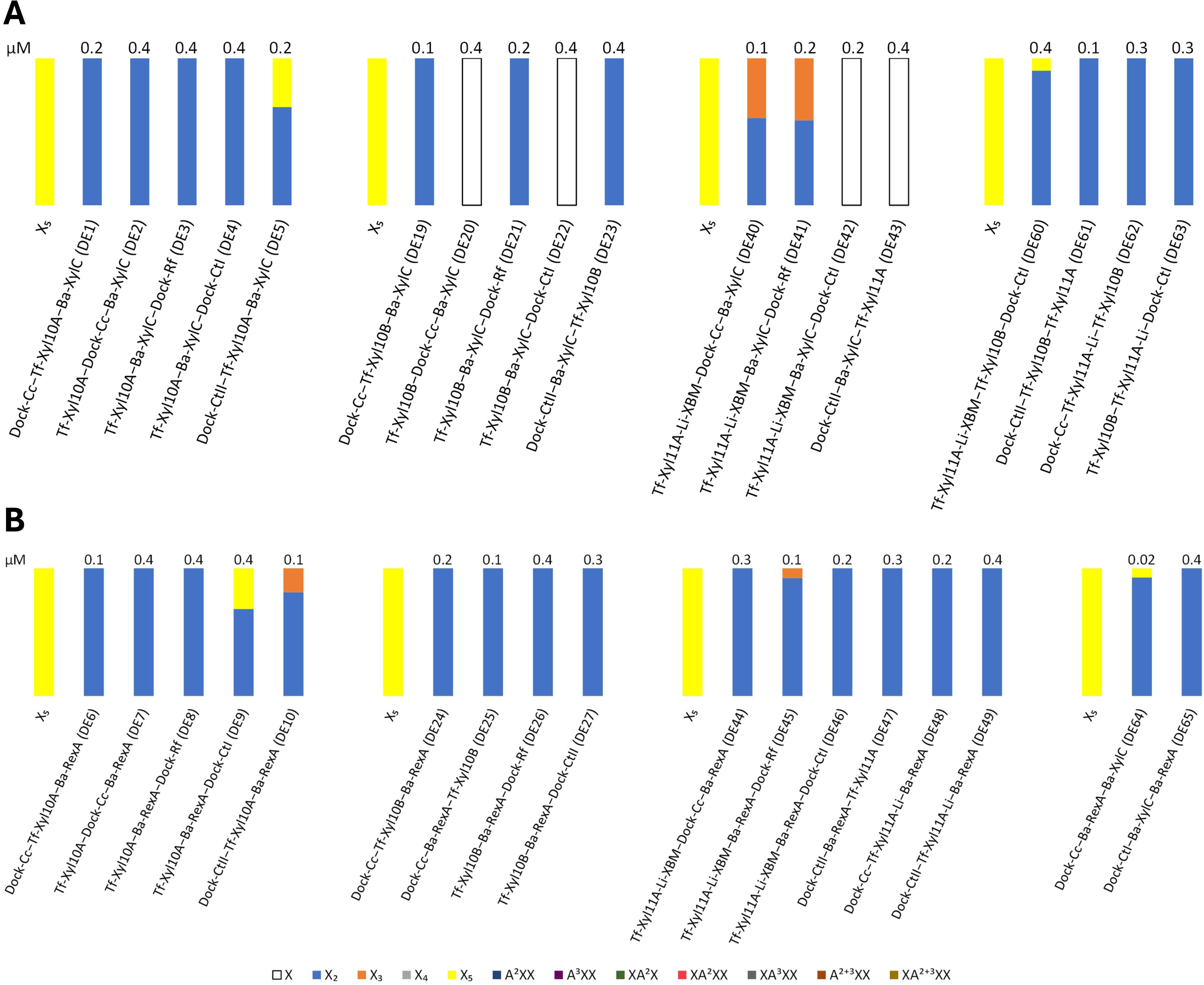

### 1.11. Parallel one-step DX assembly and purification

Our approach aims to evaluate the production and activity of different DXs in parallel in a time-efficient manner and with low resource efforts. In this way, we provide a rough assessment of the most promising DEs for further investigation. This approach is particularly interesting when the number of DEs to be incorporated into a DX is large. To reduce purification time and costs, our strategy consists of combining *E. coli* lysates containing DEs and scaffoldin. Due to the high dockerin-cohesin affinity, the complex can be formed and subsequently purified by affinity chromatography using the GST-tag present at the N-terminus of the scaffoldin.

DEs with potential catalytic activities were chosen to construct DCs with four different catalytic domains (*i.e.* resulting in a bivalent scaffoldin with two bicatalytic DEs) that would completely or partially degrade AX into (A)XOS. DE selection was performed based on a critical analysis of the SDS-PAGE, WB, ELISA and DSA-FACE results. This narrowed our selection to docking enzymes DE7, DE20, DE28, DE30, DE48, DE74, DE88 and DE94 (**Table 1**). To estimate the volumes of lysates to be combined for DX assembly, WB and dot blot assays were used (**Supplementary File S10**). By adding an excess of DEs in comparison to the scaffoldin, we aimed to maximize the probability that the purified complexes contain both DEs and that unbound DEs will be washed away before elution. The success of DX assembly was assessed by analyzing the FT, wash and elution fractions of the GST pull-down assay by means of SDS-PAGE (**Supplementary File S11**). The GST pull-down assay was done in 96-well plates to enable parallel purification of multiple DXs. Here, we performed the purification of 23 DXs and nine controls. From the 23 DXs, eight comprised a single bicatalytic DE (DC1-8) and fifteen contained two bicatalytic DEs (DX9-23). The controls were either the bicatalytic DE alone (C1-8) or the scaffoldin alone (C9). In general, SDS-PAGE analysis of DX1-8 clearly showed that all DEs bind to the bivalent scaffoldin. In most cases, DEs also appeared in the FT and wash fractions, indicating an excess DE was present, as intended. In case of DXs containing two bicatalytic DEs, SDS-PAGE analysis for DX9, DX12-13, DX15 and DX17-19 showed clear binding for both DEs. Successful binding of the remaining DXs (DX10-11, DX14, DX16, DX20-23) could not be verified. In these cases, either the three expected bands for the DX components were not present at all, or could not be identified due to the lysate background and/or having similar MWs.

### 1.12. Comparison of hydrolytic activities of the assembled DXs

To analyze the hydrolytic capabilities of the assembled multi-enzyme complexes, all DXs were incubated with high viscosity wheat flour arabinoxylan (MW: 370 kDa, A/X ratio of 38/62). The X monomers are bi- and mono-substituted with *O*-2/*O*-3 and *O*-3 A residues, respectively. The rate of released sugars was measured using the DNS method and quantified using dedicated biochemical kits. The reducing end signal is mainly a measure for the (A)XOS released from AX, as the amount of released X and A as measured by the biochemical kits was below the range of the DNS calibration curve (0.5 and 8 mM). The maximum released amount of X and A was 225 and 128 µM, respectively. The amount of X or A found in the hydrolysates was divided by the maximum amount of X or A that could be theoretically released from the substrate (39 mM for X and 23 mM for A). The expected products are shown in **Table 3**, while the hydrolytic efficiencies of the DXs are shown in **Figure 9**.

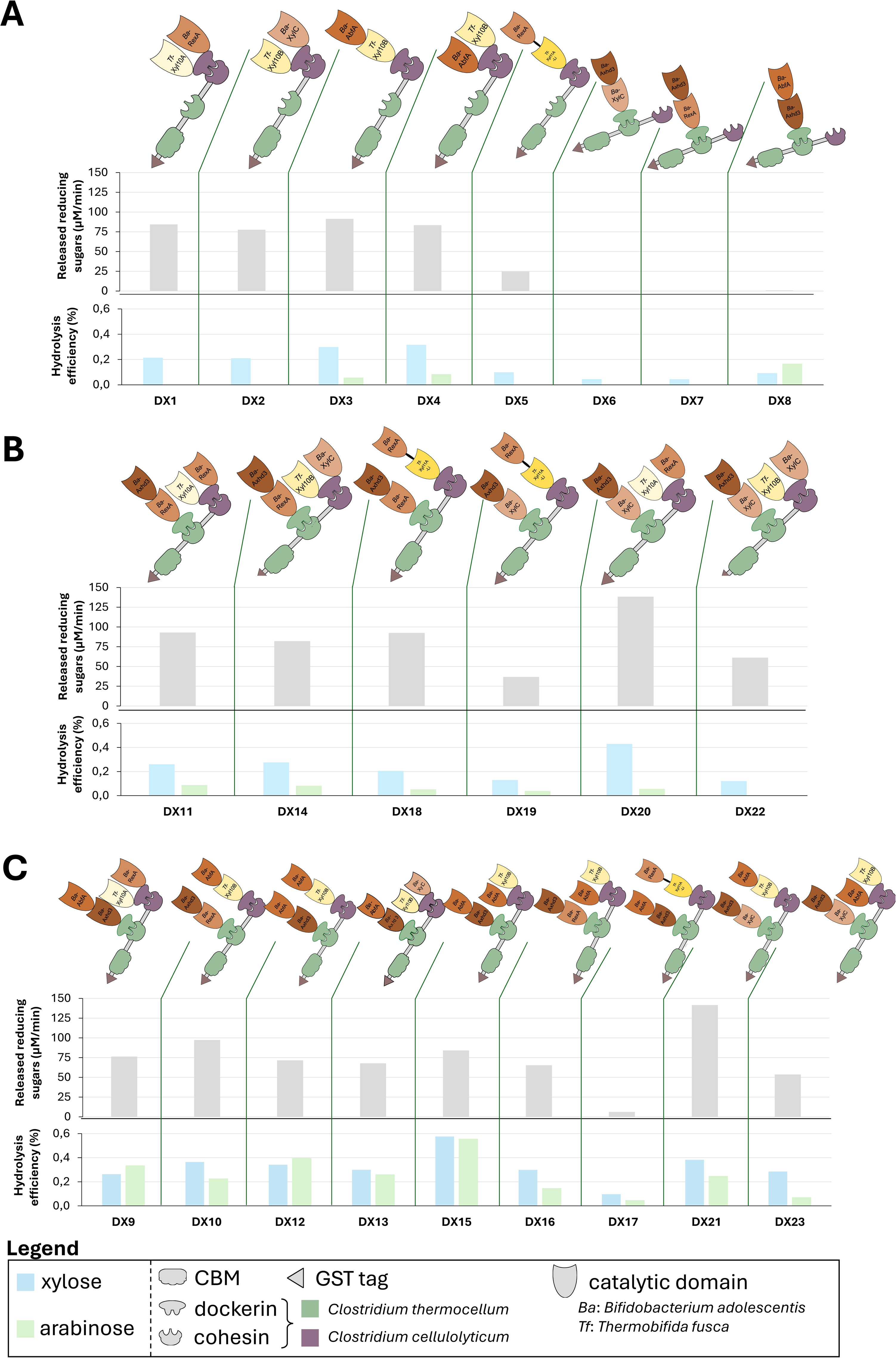

**Table 3.**
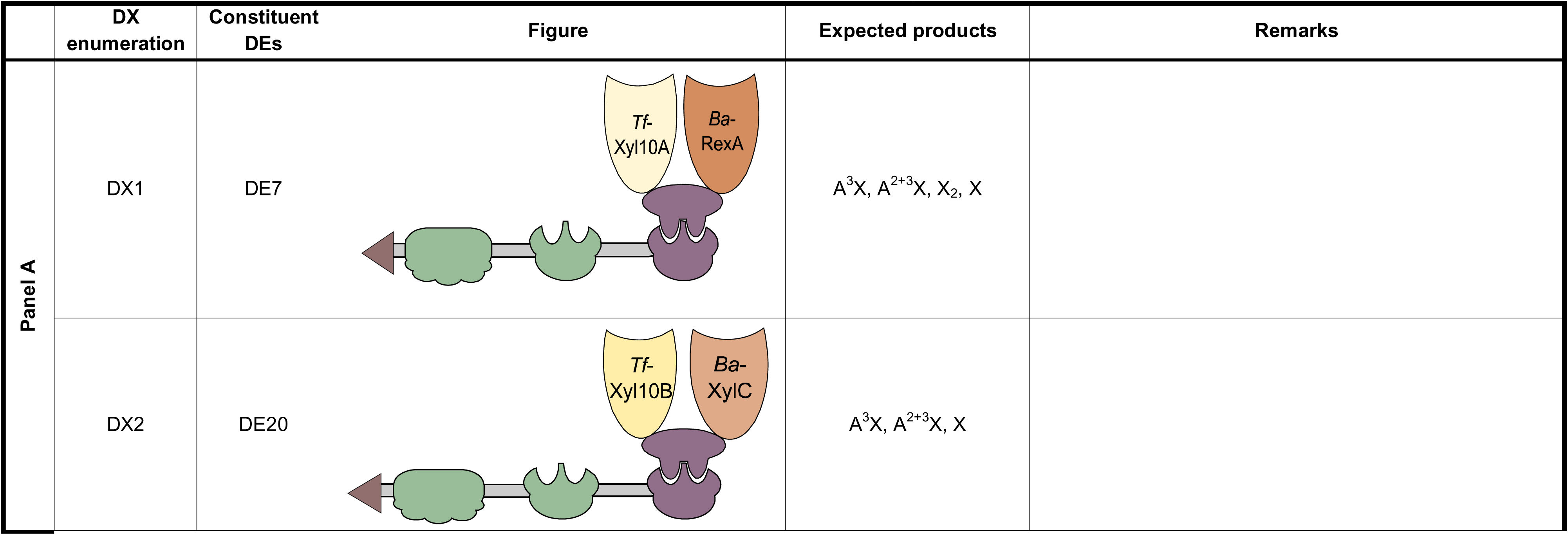
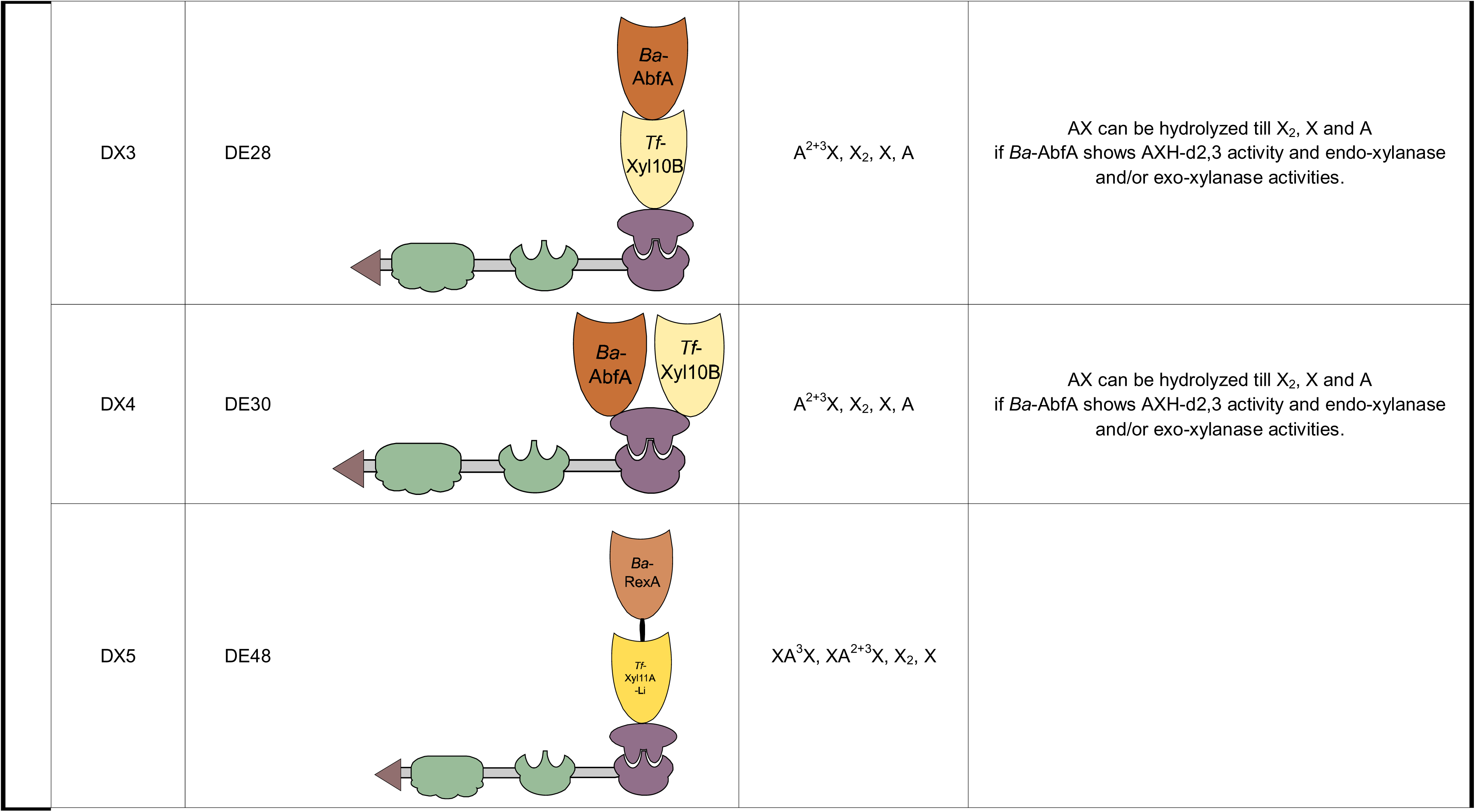
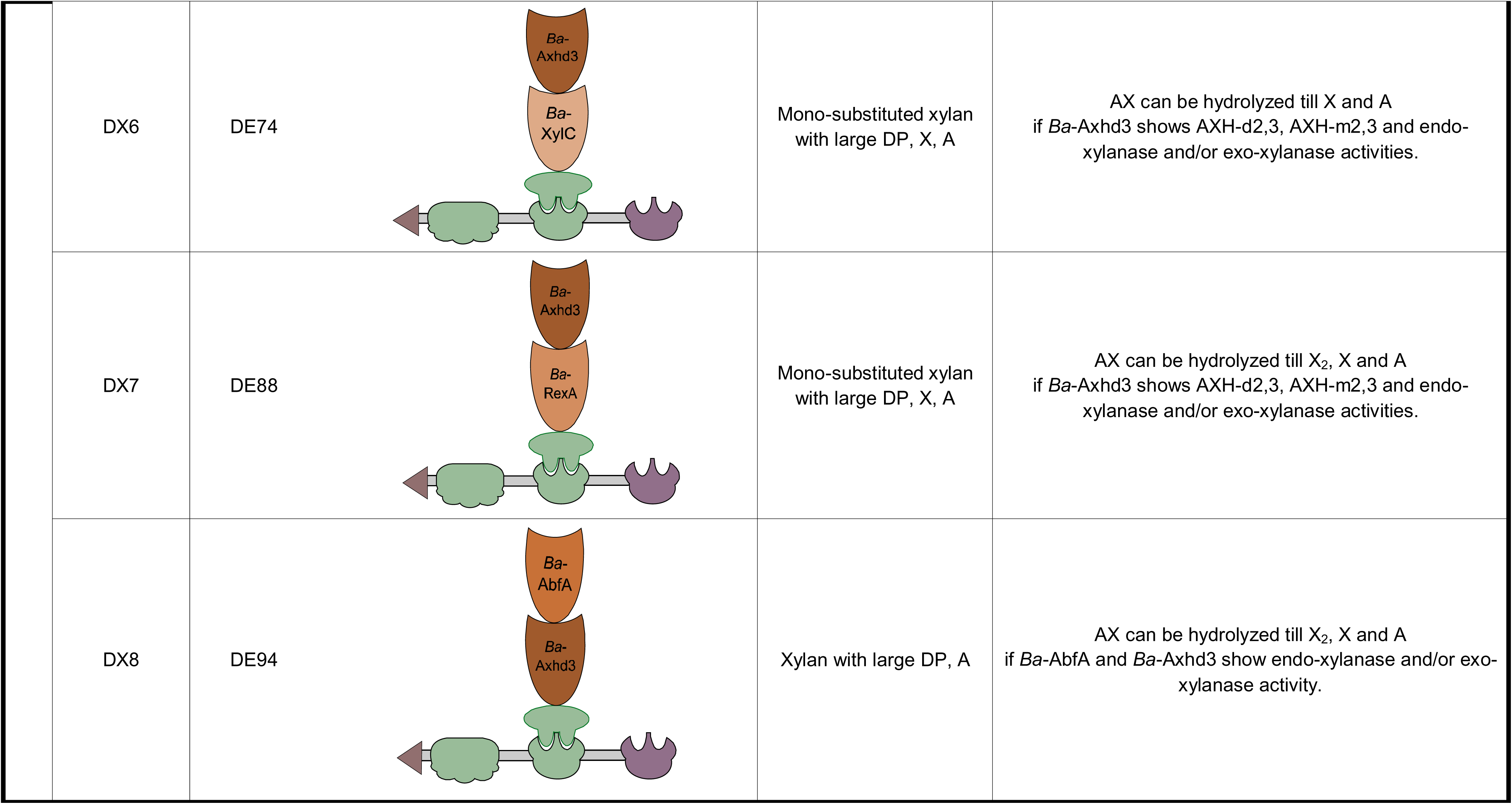
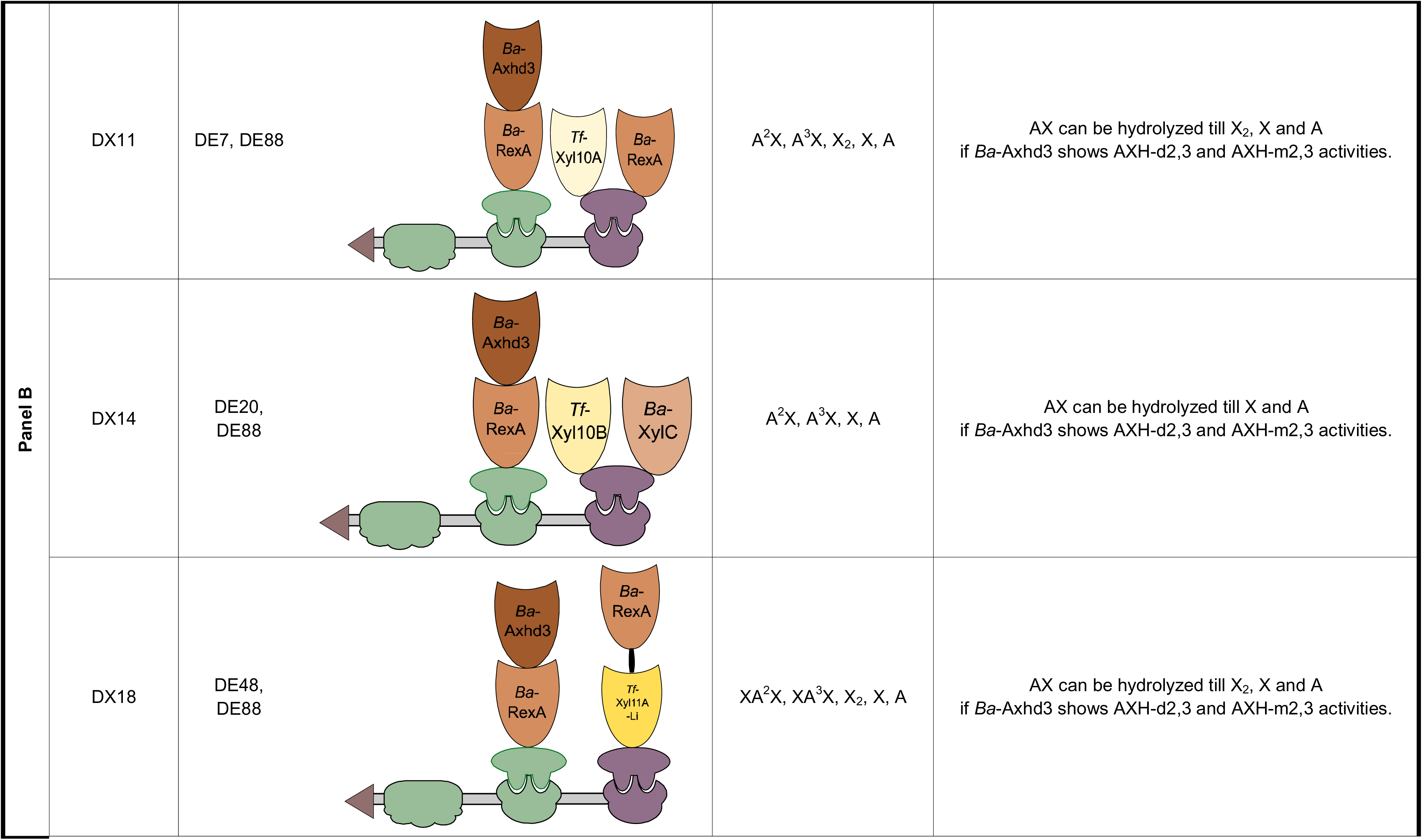
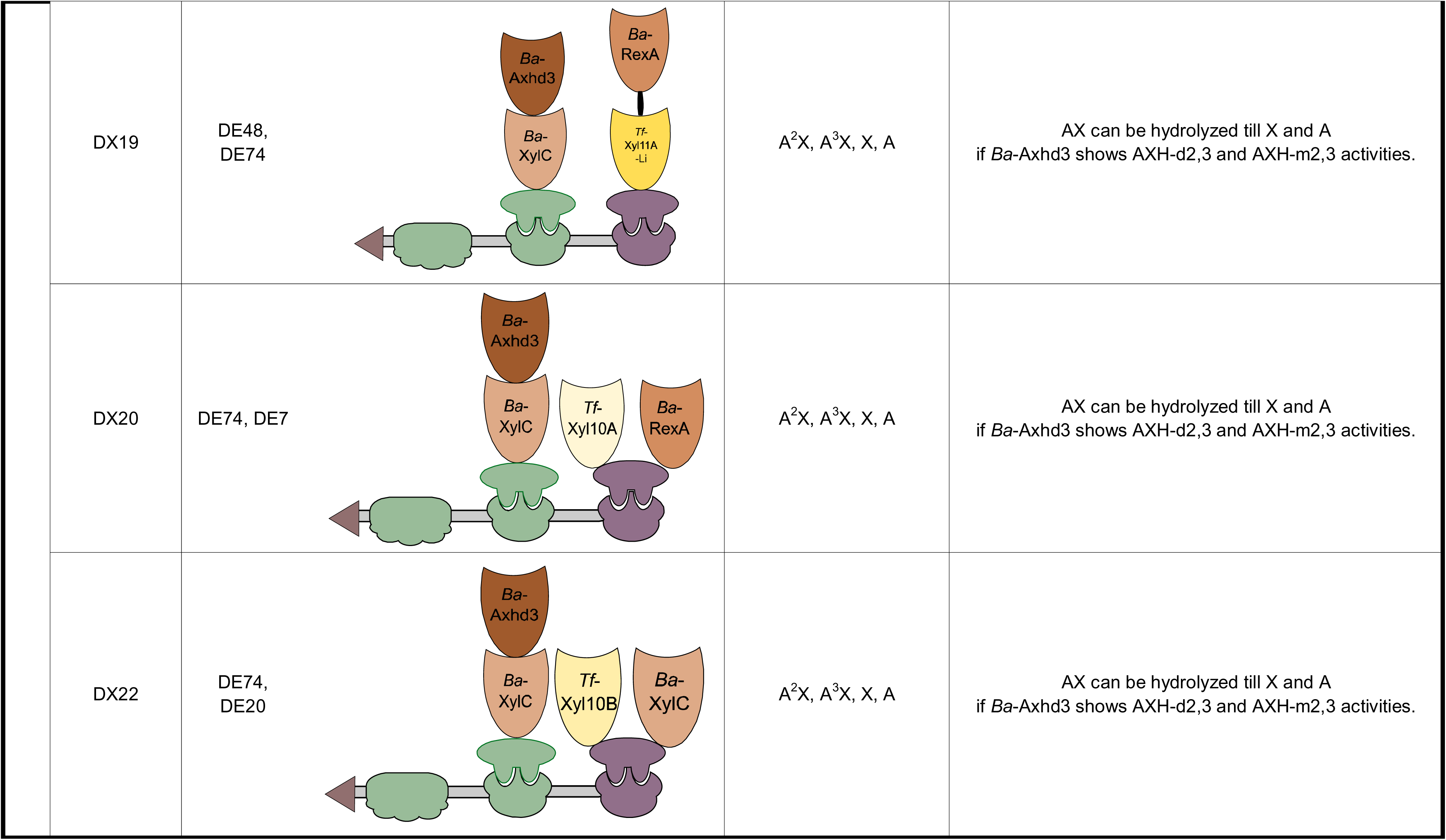
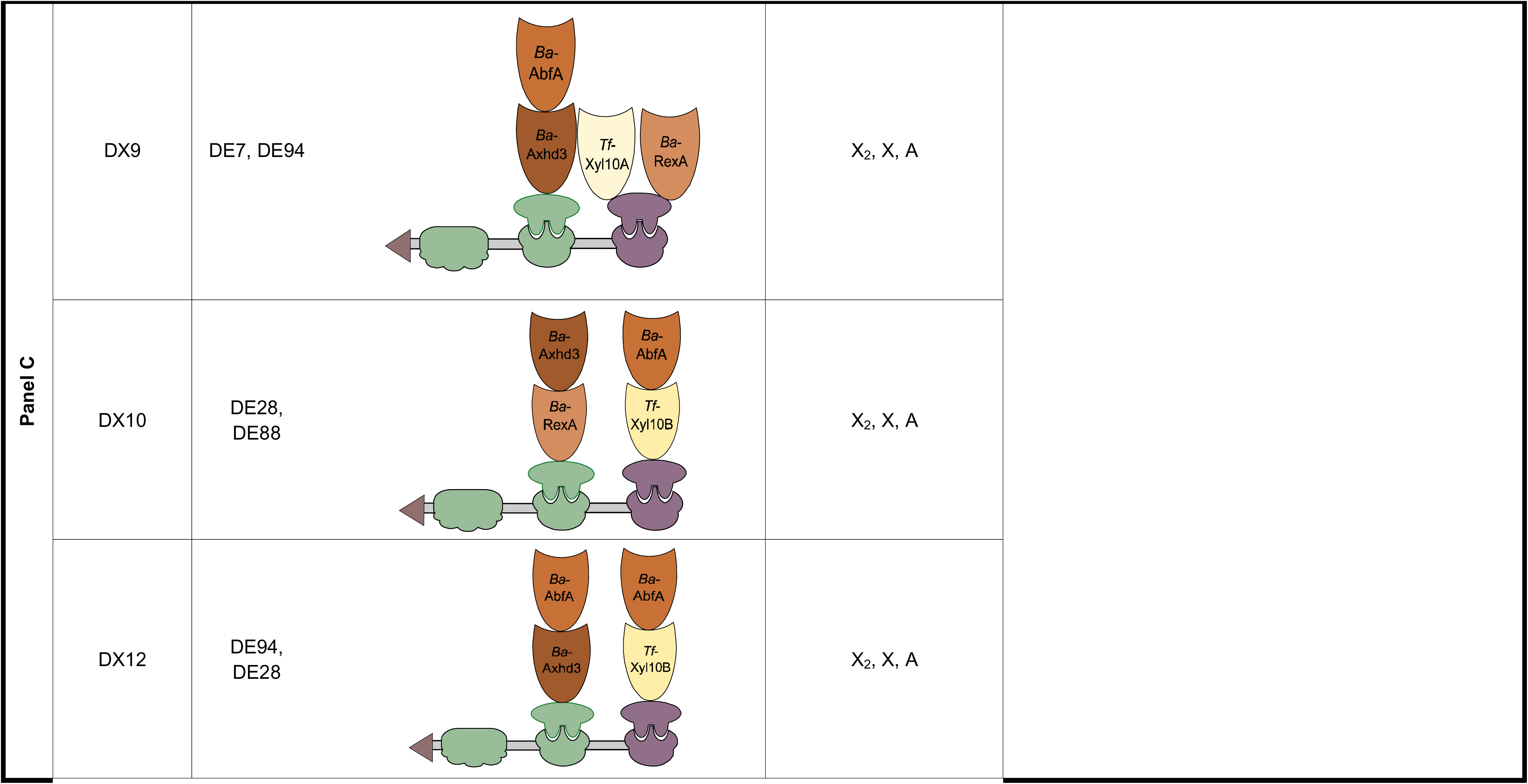
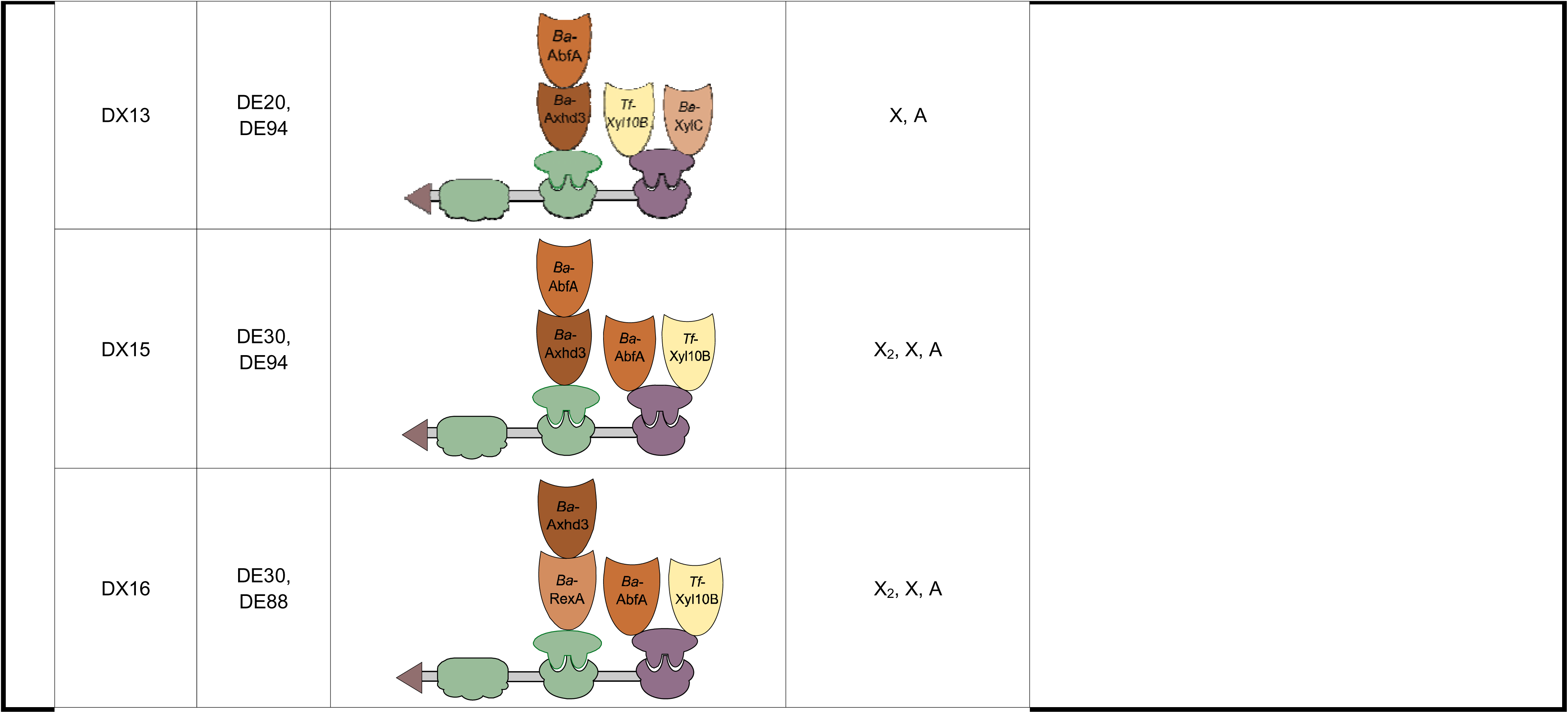
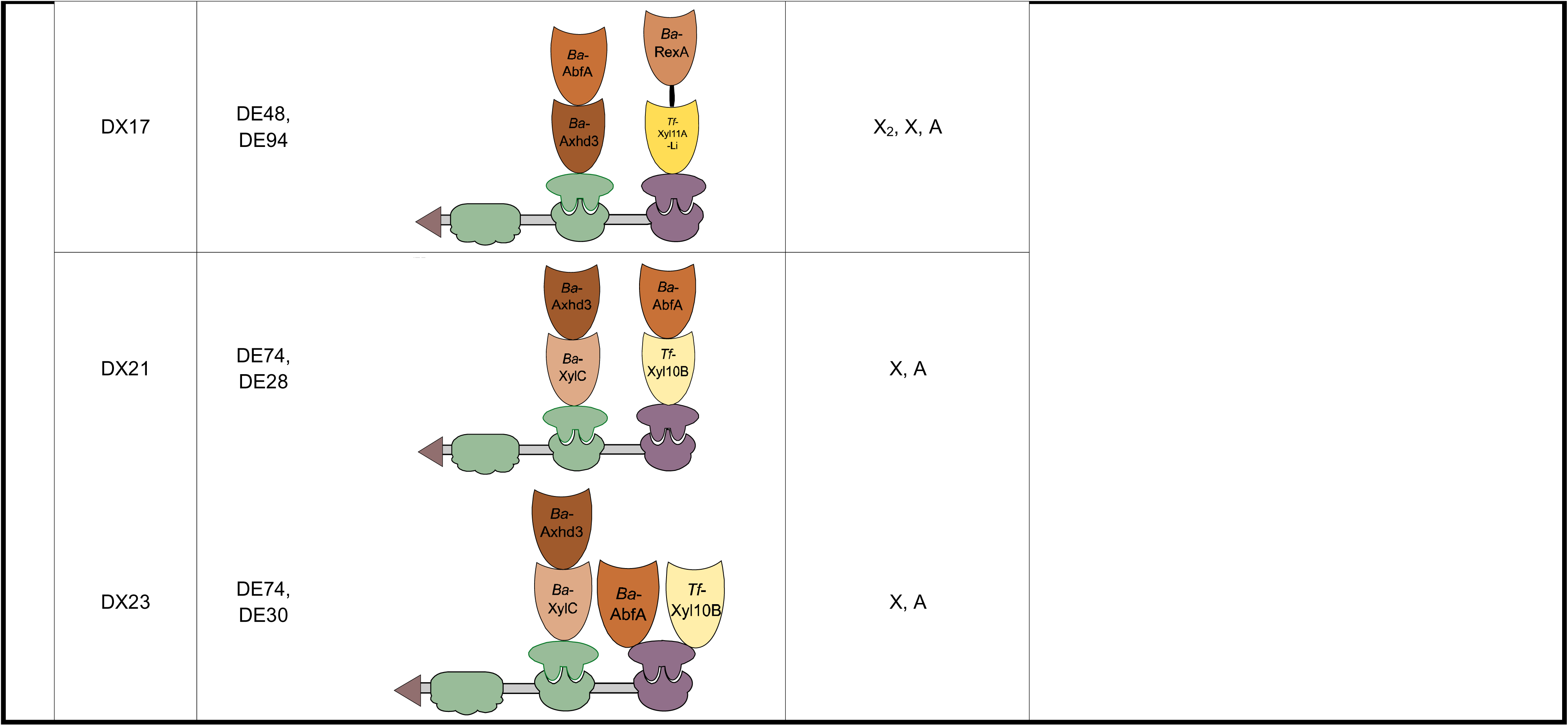
Products expected to be obtained upon incubation of the newly constructed DXs, assuming complete digestion. The expected products are determined according to the known substrate preferences from literature and the observed activity upon testing the separate DEs. DXs are organized in this table corresponding to panels A-C from **Figure 9**. (**A**): DXs comprising a single bicatalytic DE; (**B**): DXs with two bicatalytic DEs resulting in partial AX degradation; (**C**): DXs with two bicatalytic DEs resulting in total AX degradation to monosaccharides. Note that the analysis of *Ba*-AbfA- and *Ba*-Axhd3- containing DEs revealed some unexpected activities. The difference between the DXs of panel B and panel C, is that the latter contains both arabinofuranosidases able to completely remove arabinose substitutions from AX.

In **Figure 9A**, DX1-5 containing 1 of the endo-xylanases tested, have similar rates of AX hydrolysis, except for DX5. Indeed, *Tf*-Xyl11A is known to be more hindered by A substitutions and hydrolyzes glycosidic bonds more distant from Xs substituted with A. Interestingly, DX3-4 that contain *Ba*-AbfA, release higher amounts of X than DX1-2. This suggests that either removal of A enhances endo-xylanase activity and/or that *Ba*-AbfA is able to release X from AX. The latter could be in agreement with the earlier observation that *Ba*-AbfA may have endo-/exo-xylanase activity (**Figure 8**). DX6-8 did not release any detectable reducing sugars. This is unexpected as these DXs contain A- and/or X-releasing enzymes (*Ba*-AbfA and *Ba*-Axhd3, and *Ba*-XylC and *Ba*-RexA, respectively), but might be explained by the lower sensitivity of the DNS method compared to the measurement of X and A using the dedicated kit. DX6-7 release minor amounts of X but no detectable A. Note that DX6-7 contain DE74 and DE88, respectively. For both DEs, the ability to remove A from XA^2+3^XX was shown in their product profiles (**Figure 6**). It is possible that the *Ba*-Axhd3 domain is not active under the conditions tested, or that the incorporation of *Ba*-Axh3 in the DX renders the *Ba*-Axh3 domain inactive (*f.i.* through sterical shielding). Furthermore, it could be that *Ba*-Axhd3 releases undetectable amounts of A from AX. DX8, comprising DE94, releases X and A, which was also seen for DE96 (**Figure 8**).

**Figure 9B** shows hydrolytic activities of DXs that would convert AX into mono-substituted AXOS and XOS/X/A (**Table 3**). DX20 has the highest DNS signal, indicating the highest AXOS yield, likely (A)XOS with a lower DP and degree of substitution (DS). The *Ba*-Axhd3 module of DX20 will remove AX double substitutions and boost the access of the endo-xylanase *Tf*-Xyl10A and X-removal *Ba*-XylC and *Ba*-RexA. This is particularly interesting when the conversion of AX into prebiotic (A)XOS is desired [65]. To understand the extent of AX degradation and to know the type of (A)XOS that were formed, the DSA-FACE electropherograms of the hydrolysates were compared to the electrophoretic mobilities of XOS standards (X_2_ to X_6_) (**Supplementary File S11**). After 70 min reaction time, DX2 (containing *Tf*-Xyl10B) is able to hydrolyze AX to a larger extent. With increasing reaction times, DX11, DX14 and DX20 (containing *Tf*-Xyl10A and *Tf*-Xyl10B) are able to generate a mix of (A)XOS and XOS with different DPs. DX18-19 seem to preferentially generate XOS. When analyzing the electropherogram of DX22 hydrolysate, it was observed that it generates less reaction products, likely due to a lower or absent *Ba*-Axhd3 activity. Indeed, DX22 is the only variant that does not release arabinose, indicating that *Ba*-Axhd3 functionality is truly context-dependent.

**Figure 9C** shows hydrolytic activities of DXs converting AX into X_2_, X and A (DX9-10, 12, DX15-17) or completely into X and A when also *Ba-*XylC is present (DX13, DX21 and DX23) (**Table 3**). Whereas there are only single replicates, the observed differences appear substantial with a high diversity in both the reducing signal and released amounts of A and X. The average reducing end signal (74 µM/min) is not higher than in **Figure 9B** (84 µM/min), but generally an elevated release of X and particularly A is seen in **Figure 9C**. The increased levels of A can be explained by the inclusion of *Ba*-AbfA. DX15 shows the highest A and X release efficiencies and DX21 cleaves AX into higher amounts of reducing sugars. This indicates a better synergy between DEs of these complexes in comparison to the other DXs. DX21 shows a comparable reducing end and X signal to DX20 (**Figure 9B**), but higher A release.

Overall, the hydrolysis efficiencies are low, ranging between 0.04-0.58% for xylose release and 0-0.56% for arabinose release. The observed low yields may partly reflect inhibitory effects caused by the production of AXOS, which are known to exert strong feedback inhibition on glycoside [66]. Providing a detailed kinetic analysis of product inhibition was beyond the scope of this work. However, the possibility of AXOS accumulation constraining further hydrolysis should be considered when translating these findings towards (industrial) application scales. In such contexts, mitigation strategies such as continuous product removal, enzyme engineering to reduce inhibitor affinity, or coupling downstream bioprocesses might be required to achieve higher hydrolysis efficiencies. Furthermore, it must be emphasized that DEs with different temperature optima might be combined into a final DX, which could explain why several DXs are enzymatically inactive.

## 4. Conclusions

In nature, enzymes often work in synergy to degrade lignocellulosic polymers. Due to the complexity of the lignocellulosic substrates, anaerobic microbes have developed efficient enzyme systems with varied substrate preferences able to tackle the crystalline intertwined lignocellulosic polymers. Specifically, many enzymes show a modular structure containing active domains with different substrate preferences and non-catalytic domains like CBMs to stabilize the enzyme and/or to direct the enzymes to the target substrate. Since 2001, researchers have been attempting to mimic the natural cellulosomes by engineering their modular composition and adding diverse dockerin-cohesin pairs, enzyme activities, accessory modules, linkers, etc [12]. When looking at the natural repertoire of catalytic domains, for instance, one will encounter diversity at different levels as microbial origin, substrate preference, kinetic parameters, optimum reaction parameters (pH, temperature and cofactors needed) and environmental influence. All these factors will sum up into bicatalytic DEs with specific expression, solubility, purity and stability levels, substrate preferences, dockerin-cohesin affinities and activity performance properties. However, reaching similar hydrolytic levels is still challenging partially due to the large dimension of natural cellulosomes: *Clostridium clariflavum* can express enzyme complexes containing 160 enzymes [1]. Although larger scaffoldins could accommodate more DEs in the context of DCs, and in this way resemble the large natural cellulosomes, increased scaffoldin sizes for DC construction have been correlated with lower expression levels [67].

Therefore, inspired by the modular architecture of natural cellulosomes, we developed a parallel (semi-) high-throughput pipeline for the design, expression and functional screening of bicatalytic DEs active on AX. Central to this approach is the VersaTile platform, a combinatorial DNA assembly technology that enables the rapid generation of large libraries of modular enzyme constructs. Combined with parallel expression, purification and activity assays, this platform offers a scalable strategy to explore the vast sequence and architectural diversity found in nature’s enzymatic toolbox.

In this study, we demonstrated the power of this funnel approach by constructing and screening 96 bicatalytic DEs, comprising dockerins and various AX-active catalytic domains. Bicatalytic DEs also offer the possibility to optimize the ratio of enzymes for an optimal biomass polymer degradation as it has been observed that in some cases a higher presence of certain activities is necessary in comparison to others [14]. DEs were chosen in order to obtain a representative set of enzyme activities and modular orientations. The DEs were systematically analyzed for expression levels, stability, substrate preference and dockerin-cohesin binding affinity using streamlined workflows based on microplate formats (24/96-well). The DEs were produced and purified in less than one week, followed by enzymatic assays and binding tests, allowing the characterization of nearly 100 DEs in approximately 10 working days (**Figure 10**). Substrate preferences of the new 96 bicatalytic dockerin-containing AX-active enzymes were tested by performing enzymatic reactions with the purified fractions of the enzymes and representative (A)XOS. Afterwards hydrolysates were labelled with the fluorescent tag APTS and analyzed by DSA-FACE. Product profiles of the reaction hydrolysates of the different variants can then be compared to assess substrate preferences of the DEs. ELISA was done to test the affinity between the dockerin present in the variants and the scaffoldin containing corresponding cohesins. Differences in temperature optima are possible and might explain why some DEs did not make it to the stage of DX assembly.

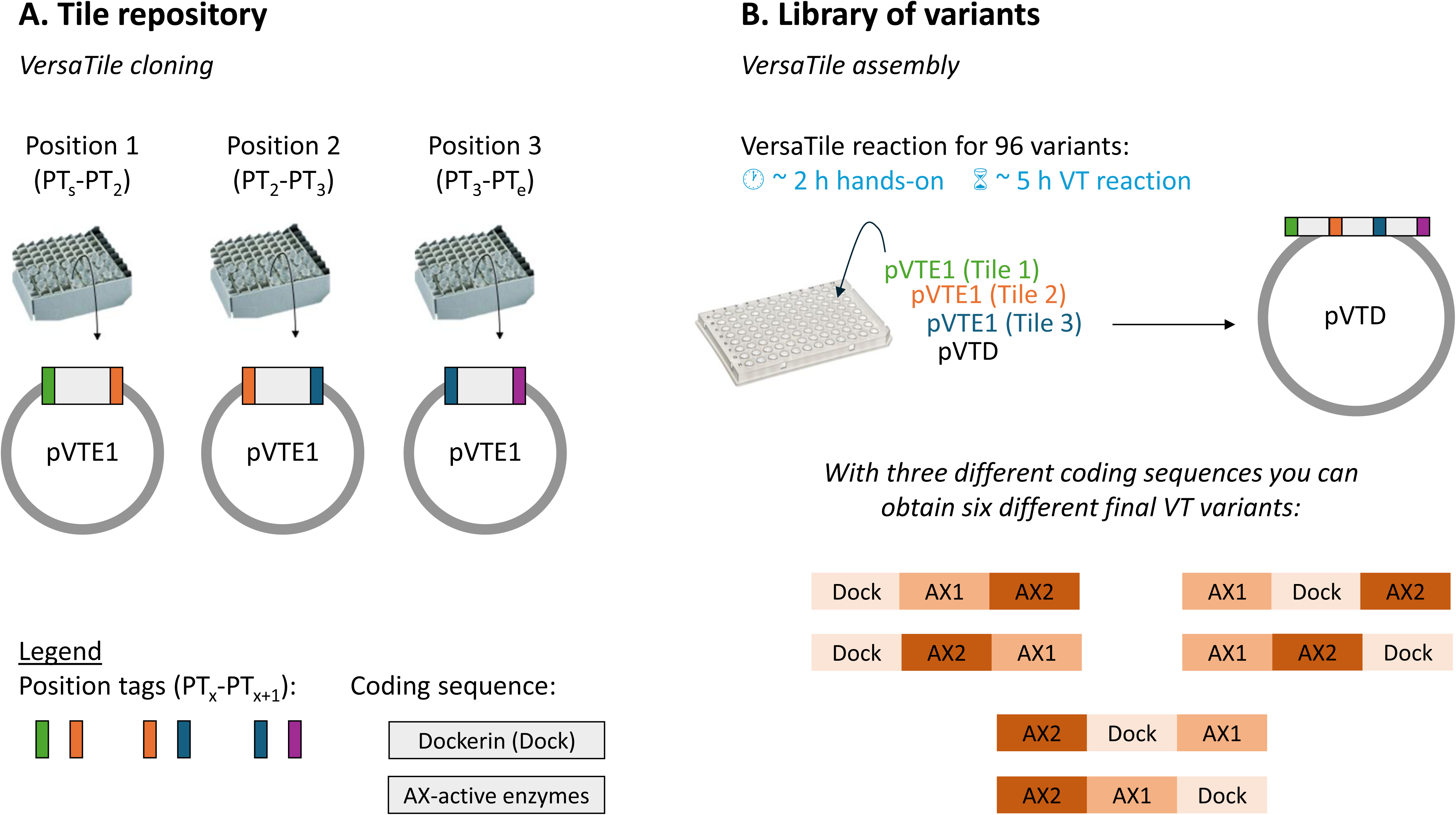

A key strength of the proposed pipeline is its potential for large-scale, omics-style exploration of modular enzyme diversity. The VersaTile platform enables the combinatorial generation of hundreds to thousands of DEs in a matter of days (*i.e.* with theoretical capacities of 464 DEs, 48 scaffoldins, resulting in >645000 DCs) (**Figure 11**). However, while the construction of DE libraries has been greatly accelerated, the functional characterization remains an important bottleneck. Screening large numbers of enzyme variants for activity, specificity and stability still demands substantial time, materials and infrastructure. In this context, the use of DSA-FACE represents a significant step forward. Compared to traditional methods like HPAEC-PAD, DSA-FACE is faster, requires smaller sample volumes, allows parallel processing and yields detailed product profiles for a wide variety of substrates. These features make DSA-FACE particularly suitable for (semi-) high-throughput activity screening, enabling researchers to screen large enzyme libraries and prioritize the most promising variants for further study.

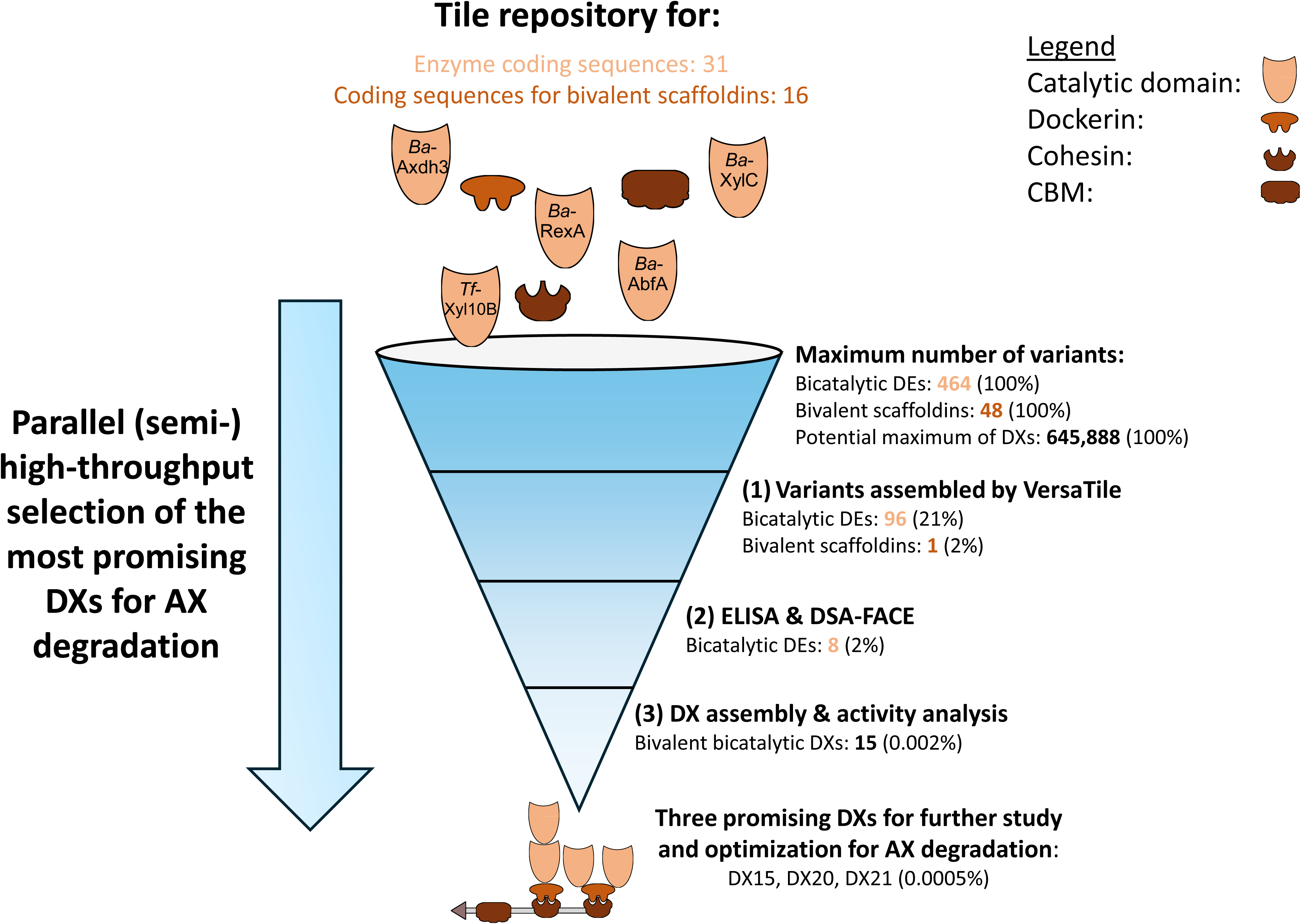

Despite a high frequency of degradation and aggregation phenomena among the DEs (62%), up to 89% of the retained DEs showed detectable hydrolytic activity on (A)XOS. Interestingly, the order and combination of catalytic modules influenced activity and degradation patterns, illustrating the need for empirical exploration in modular enzyme engineering. For example, DE59 (Dock-*Ct*II – *Tf*-Xyl11A – *Ba*-Axhd3) showed enhanced activity compared to the reversed arrangement in DE57, thereby stressing the importance of domain architecture on DE performance. In some cases, remarkable activities were observed [30]. Upon removal of arabinose from doubly substituted xyloses, DEs containing α-L-arabinofuranosidase *Ba*-Axhd3 (DE15− 18, DE38, DE55−59, DE86−88, DE90) hydrolyzed the remaining *O*-2 arabinose from mono-substituted xyloses. Whether this activity is a result of the new arrangement of the newly constructed enzymes, in the case of DE57 (Dock-*Ct*II− *Ba*-Axhd3−*Tf*-Xyl11A) and DE58 (Dock-*Cc*−*Tf*-Xyl11A-Li−*Ba*-Axhd3), must be further investigated. It could be investigated if this activity is enhanced by the presence of cofactors like Ca^2+^, due to enzyme dosage and/or due to a specific *Ba*-AbfA/*Ba*-Axhd3 structural arrangement. Fusing enzymes can result in architectures characterized by protein-protein interactions that will allow the accommodation of unexpected linkages in the active site of the enzymes. From the DE library, eight promising DEs were selected for incorporation into 23 AX-active designer cellulosomes, which we call here designer xylanosomes (DXs). The DXs comprised either one or two bicatalytic DEs. Screening of these DXs against high viscosity wheat flour AX, which is a more complex and industrially relevant substrate, revealed substantial variability in hydrolytic efficiency. Notably, DX15, DX20 and DX21 demonstrated superior or synergistic degradation performance and are currently considered strong candidates for further optimization and application.

Our findings highlight DSA-FACE by enabling faster, more sensitive and scalable activity screening. Nevertheless, further advancements in analytical throughput are essential to unlock the full potential of DE/DC libraries. Obviously, there is room for adaptation/optimization of the methodological pipelines developed to match industrial reaction conditions and bring DC hydrolytic efficiencies to increased levels. The options are endless and high-throughput approaches that offer time and cost-saving alternatives remain a major need. Furthermore, our future efforts will focus on integrating yeast surface display systems and automated screening platforms to further streamline the identification of optimal enzyme assemblies. As modular enzyme engineering moves into the era of synthetic biology and big data, pipelines as described in this study will be essential to screen and manage the immense combinatorial diversity required for next-generation biomass conversion technologies.

The presented funnel-type methodological pipeline may be extrapolated to other cases where complete or partial hydrolysis of complex plant polysaccharides (*f.i.* pectic rhamnogalacturonan I and II, fucogalacto-xyloglucan, *O*/*N*-glycans, …) may be desired. Moreover, the principles of our funnel-type methodology based on combinatorial design may be of use in any process or industrial context in which complex enzyme cascades are required for the modification of specific biomolecules and fine chemicals [68].

In conclusion, we have presented a funnel-type approach for the rapid construction and screening of enzymes for the assembly of arabinoxylan-active designer xylanosomes. Our results indicate that the utilized approach is fit for selecting a manageable number of promising candidates for further characterization, optimization, up-scaling and utilization on industrially relevant side streams.

## 5. Declarations

### Author contributions

MF and YB conceptualized and designed the experiments. MF and JV prepared the tiles. JV established the ELISA protocol with the polyclonal rabbit anti-CBM primary antibody. SV compiled all the VersaTile variants possible to be made with the tiles available and contributed to the choice of which VersaTile variants would be used in the final experiments for this chapter. JV designed and constructed the monovalent scaffoldins used in this work. MF, JV and SV performed preliminary small scale enzymes expression, purification and activity analysis experiments to develop the (semi-)high-throughput parallel expression, purification and activity analysis protocol. JV established the GST-pull down protocol and has designed and constructed tiles for the GST-tag. MF performed all the experiments, the data analysis and interpretation of results. MF wrote the first draft. TDC reorganized and redrafted the final version of the manuscript. MF, JV, TDC and YB revised the work critically.

### CRediT

Conceptualization: MF, YB

Methodology: MF

Validation: MF

Formal analysis: MF

Investigation: MF, JV, SV

Writing - Original Draft: MF, JV, TDC

Writing - Review & Editing: JV, MF, TDC, YB

Visualization: MF

Supervision: YB

Project administration: MF

Funding acquisition: YB

MF: Conceptualization, Methodology, Validation, Formal Analysis, Investigation, Writing – Original draft, Writing – Review & Editing, Visualization, Project administration

JV: Investigation, Writing – Original Draft, Writing - Review & Editing

TDC: Writing – Original Draft, Writing – Review & Editing

SV: Investigation

YB: Conceptualization, Writing – Review & Editing, Supervision, Funding Acquisition

## Acknowledgments

The authors thank Hans Gerstmans and Boris Bekaert for designing and creating the tiles for the dockerins used in this study, respectively. We thank Dennis Grimon for the design and generation of the vectors used in this study and for all the support in tile design. The authors wish to acknowledge prof. Ed Bayer for his contributions to the research track focusing on designer cellulosomes.

## Data availability statement

All data generated in this study is available as supplementary files and/or available upon request by the authors.

