## Supplementary File S2 for "A funnel-type pipeline for accelerating discovery of tailored designer xylanosomes for efficient arabinoxylan valorization"

**Supplementary File S2: Primers used to amplify AX-active enzymes for the tile repository**

Endo-xylanases Xyl10A, Xyl10B and Xyl11A from *Thermobifida fusca*, β-xylosidase XylC, reducing-end xylose-releasing exo-oligoxylanase RexA and α-L-arabinofuranosidases AbfA and Axhd3 from *Bifidobacterium adolescentis* were converted to tiles to be compatible with the VersaTile assembly technique. The first column shows the name of the tile which comprises the name of the enzyme whose coding sequence will be present in the tile flanked by the position tag codes (PTx-PTx+1), which indicate the position of the tile in the final construct (with S and E referring to ‘start’ and ‘end’, respectively). Next to it the number of BsaI recognition sites present in the enzyme coding sequences is displayed. This is important since BsaI coding sequences must be deleted prior to VersaTile assembly. In the column ‘linker’ it is stated whether or not the enzyme coding sequences were amplified with their natural linker between the N-terminal catalytic domain and C-terminal CBM. This is only the case for *Tf*-Xyl10A and *Tf*-Xyl11A since the other enzymes did not contain a CBM. In all other cases, the complete coding sequence was amplified. The next columns show the different sequences that the VersaTile compatible forward and reverse primers comprise. Starting from the 5’ end, each primer contains three nucleotides for clamping of the restriction enzyme, the recognition/restriction site of a specific restriction enzyme and the BsaI recognition site. Afterwards there is the sequence of the position tags (PTx-PTx+1) (restriction site of BsaI) and the sequence that is complementary with the fragment of interest. *In red, G nucleotide was replaced by A in the forward primer and T in the reverse primer (silent mutations) to delete the recognition site for the BsaI restriction enzyme and to obtain a better design of the primers, respectively.

| **Tile** | **N° BsaI** | **Linker** | **Forward primer (5’🡪3’)** | | | | | **Reverse primer (5’🡪3’)** | | | | |
| --- | --- | --- | --- | --- | --- | --- | --- | --- | --- | --- | --- | --- |
|  |  |  | **3nt** | **XbaI** | **BsaI** | **Position tag (PT)** | **Complementary region** | **3nt** | **HindIII/PstI** | **BsaI** | **Position tag (PT)** | **Complementary region** |
| PT_s__*Tf*-Xyl10A_PT_2_ | 0 | No | ATA | TCTAGA | GGTCTC | ACCATG | GAGTCGACCCTGCGGGAACTGG | ATA | AAGCTT | GGTCTC | TGAACC | ACCGCCGCCGGAGGAGTC |
|  |  | Yes |  |  |  |  |  |  |  |  |  | GGGGCCGCCCGGTTC |
| PT_2__*Tf*-Xyl10A_PT_3_ |  | No |  |  |  | GGTTCA |  |  |  |  | ACCAGA | ACCGCCGCCGGAGGAGTC |
|  |  | Yes |  |  |  |  |  |  |  |  |  | GGGGCCGCCCGGTTC |
| PT_3__*Tf*-Xyl10A_PT_e_ |  | No |  |  |  | TCTGGT |  |  |  |  | ATACTT | ACCGCCGCCGGAGGAGTC |
| PT_s__*Tf*-Xyl10B_PT_2_ | 0 | No |  |  |  | ACCATG | GGACCGGTCCACGACC |  |  |  | TGAACC | GCAGTGATCGTGCTTGG |
| PT_2__*Tf*-Xyl10B_PT_3_ |  |  |  |  |  | GGTTCA |  |  |  |  | ACCAGA |  |
| PT_3__*Tf*-Xyl10B_PT_e_ |  |  |  |  |  | TCTGGT |  |  |  |  | ATACTT |  |
| PT_s__*Tf-*Xyl11A_PT_2_ | 2 | No | ATA | TCTAGA | GGTCTC | ACCATG | GCCGTGACCTCCAACGAAACCGGGTACCAC | ATA | AAGCTT | GGTCTC | TGAACC | GTTGCCACCGCCGCTGG |
|  |  | Yes |  |  |  |  |  |  |  |  |  | TGGTGGGTTGCCGCCACCG^*^ |
| PT_2__*Tf-*Xyl11A_PT_3_ |  | No |  |  |  | GGTTCA |  |  |  |  | ACCAGA | GTTGCCACCGCCGCTGG |
|  |  | Yes |  |  |  |  |  |  |  |  |  | TGGTGGGTTGCCGCCACCG^*^ |
| PT_3__*Tf-*Xyl11A_PT_e_ |  | No |  |  |  | TCTGGT |  |  |  |  | ATACTT | GTTGCCACCGCCGCTGG |
| PT_s__*Tf-*Xyl11A-XBM_PT_2_ | 2 | No |  |  |  | ACCATG |  |  |  |  | GAACC | GTTGGCGCTGCAGGACACCGTGG |
| PT_3__*Tf-*Xyl11A-XBM_PT_e_ |  |  |  |  |  | TCTGGT |  |  |  |  | ATACTT |  |
| PT_s__*Ba*-XylC_PT_2_ | 1 | No |  |  |  | ACCATG | ATGAAGATTTCCAACCCGGTGC |  |  |  | TGAACC | CTGGTTATCGGAAAGCTCC |
| PT_2__*Ba*-XylC_PT_3_ |  |  |  |  |  | GGTTCA |  |  |  |  | ACCAGA |  |
| PT_3__*Ba*-XylC_PT_e_ |  |  |  |  |  | TCTGGT |  |  |  |  | ATACTT |  |
| PT_s__*Ba*-RexA_PT_2_ | 0 | No |  |  |  | ACCATG | ATGACAAATGCAACCGATACC |  |  |  | TGAACC | TTCATACCGATACTTTCCGC |
| PT_2__*Ba*-RexA_PT_3_ |  |  |  |  |  | GGTTCA |  |  |  |  | ACCAGA |  |
| PT_3__*Ba*-RexA_PT_e_ |  |  |  |  |  | TCTGGT |  |  |  |  | ATACTT |  |
| PT_s__*Ba*-AbfA_PT_2_ | 0 | No |  |  |  | ACCATG | ATGACCACAACGATTACCATCG |  | CTGCAG |  | TGAACC | TGCCATGAAGCCGGC |
| PT_2__*Ba*-AbfA_PT_3_ |  |  |  |  |  | GGTTCA |  |  |  |  | ACCAGA |  |
| PT_3__*Ba*-AbfA_PT_e_ |  |  |  |  |  | TCTGGT |  |  |  |  | ATACTT |  |
| PT_s__*Ba*-Axhd3_PT_2_ | 1 | No |  |  |  | ACCATG | ATGATGATTACCTCAACTAATCCTATGG |  | AAGCTT |  | TGAACC | TTGCTCTCTTTCCTTCGTATTCG |
| PT_2__*Ba*-Axhd3_PT_3_ |  |  |  |  |  | GGTTCA |  |  |  |  | ACCAGA |  |
| PT_3__*Ba*-Axhd3_PT_e_ |  |  |  |  |  | TCTGGT |  |  |  |  | ATACTT |  |
