## Supplementary File S3 for "A funnel-type pipeline for accelerating discovery of tailored designer xylanosomes for efficient arabinoxylan valorization"

**Supplementary File S3: Primers used to remove BsaI recognition sites by inverse PCR**

The used primers contain a single silent nucleotide mutation in the BsaI recognition site and a SapI recognition site at both primers’ 5’ end. The inverse PCR is performed on the entry vector where the genes coding for *Tf-*Xyl11A, *Ba-*XylC and *Ba-*Axhd3 flanked with BsaI recognition sites and positions markers is cloned. Upon amplification and digestion with SapI, the two amplified fragments will have compatible overhangs that will be ligated upon a ligation reaction.

| **Enzyme coding sequence present in tile** | **Primer**  **(5'🡪3')** | **3nt** | **SapI+nt** | **Complementary region with point mutation** |
| --- | --- | --- | --- | --- |
| ***Tf-*Xyl11A** | Forward primer | TTT | GCTCTTCA | GT**T**TCCATGGAGCTGGGC |
|  | Reverse primer |  |  | **A**ACCGTTCCGGGCGCGT |
| ***Ba-*XylC** | Forward primer |  |  | GA**A**ACCTCCATCCAGAAAGTCG |
|  | Reverse primer |  |  | **T**TCGCGTCCGAGTGTGC |
| ***Ba-*Axhd3** | Forward primer |  |  | GA**A**ACCGGTGGAATCGCC |
|  | Reverse primer |  |  | **T**TCGCGTCCGAGTGTGC |
