## Supplementary File S8 for "A funnel-type pipeline for accelerating discovery of tailored designer xylanosomes for efficient arabinoxylan valorization"

**Supplementary File S8: SDS-PAGE and Western Blot results of the 96 AX-active DEs analyzed in this study**

Panels A-D show the SDS-PAGE (top) and Western Blot (bottom) gels of the fractions of the purified and desalted 96 bicatalytic DEs. The orange arrow points to the presumed DE. Picomole amounts loaded on the gel are written in **Table 1**. The Roti®Mark (Carl Roth) and the Precision Plus Protein™ Dual Color (Bio-Rad) standards were used for SDS-PAGE and Western Blot, respectively.

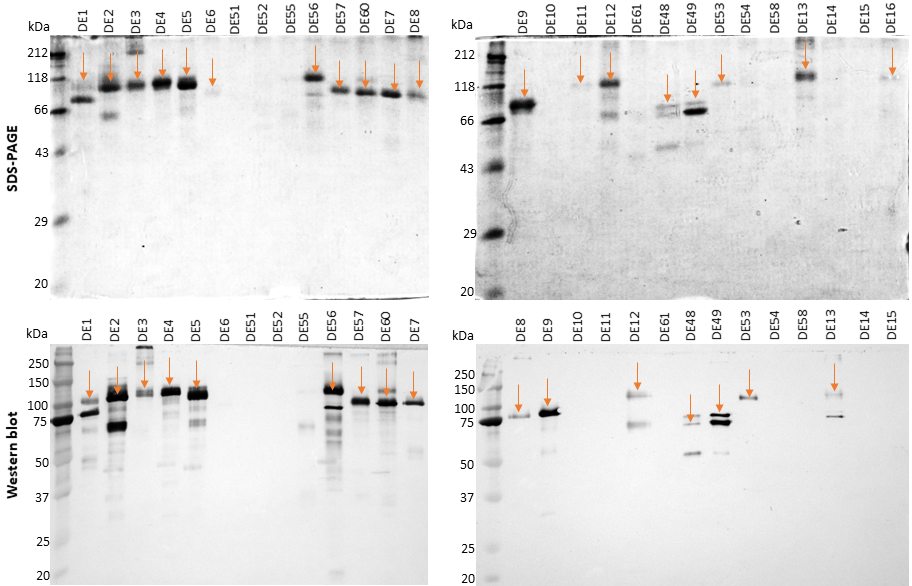

**A**

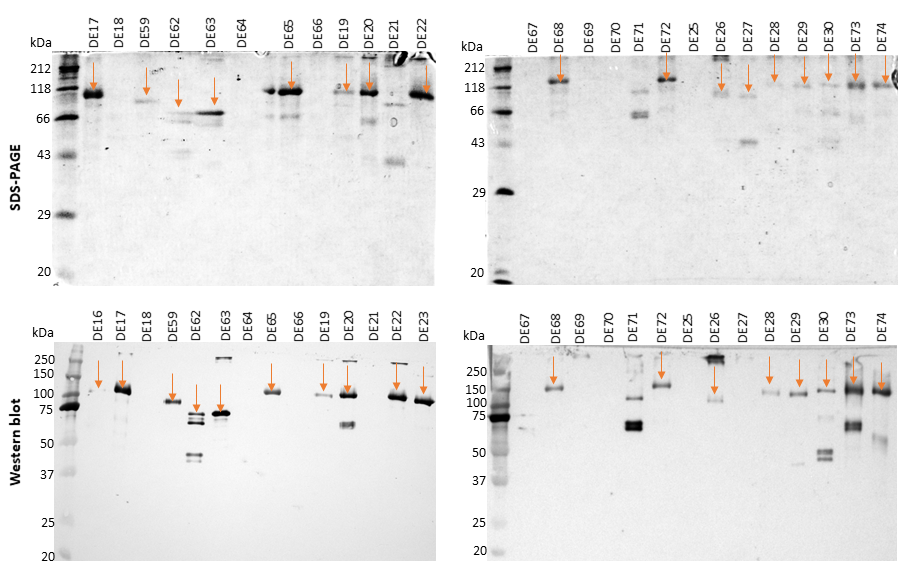

**B**

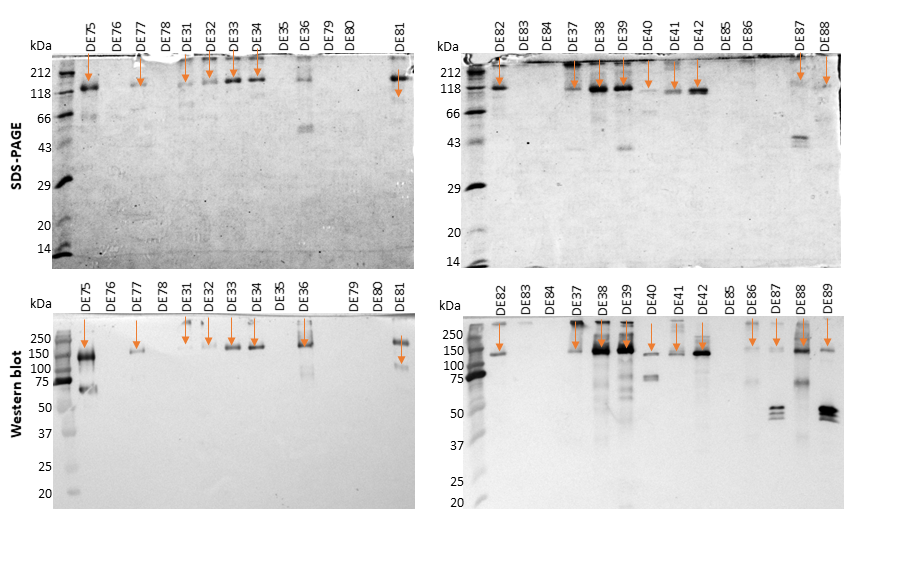

**C**

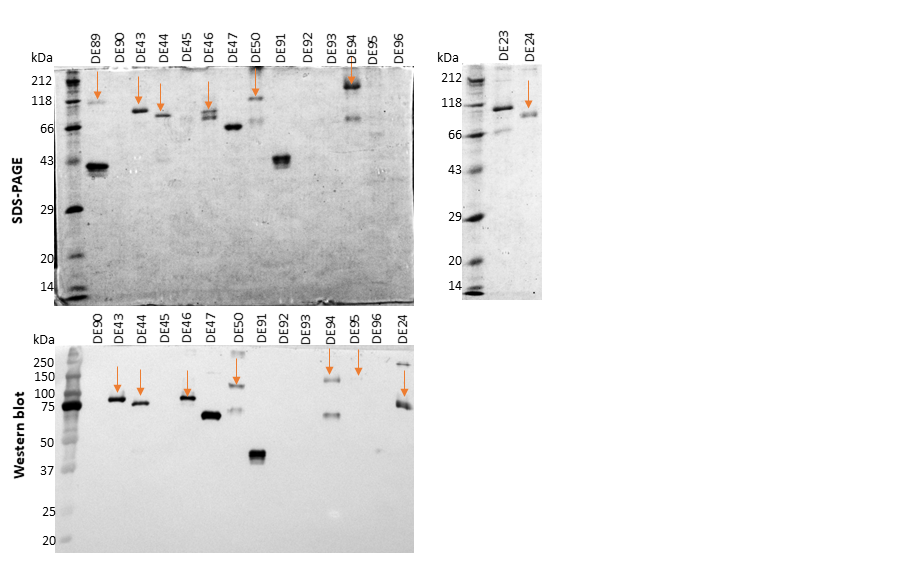

**D**
