## Supplementary File S9 for "A funnel-type pipeline for accelerating discovery of tailored designer xylanosomes for efficient arabinoxylan valorization"

**Supplementary File S9: Results of the ELISA assays for the 96 AX-active DEs against the monovalent scaffoldins**

This ELISA assay reveals the capacity of the 96 bicatalytic DEs to bind to monovalent scaffoldins containing the four cohesins that bind to the 4 different dockerins used in the DEs variants - Coh-*Ct*I, Coh-*Ct*II, Coh-*Rf* and Coh-*Cc*. The y-axis represents the absorbance at 450 nm.

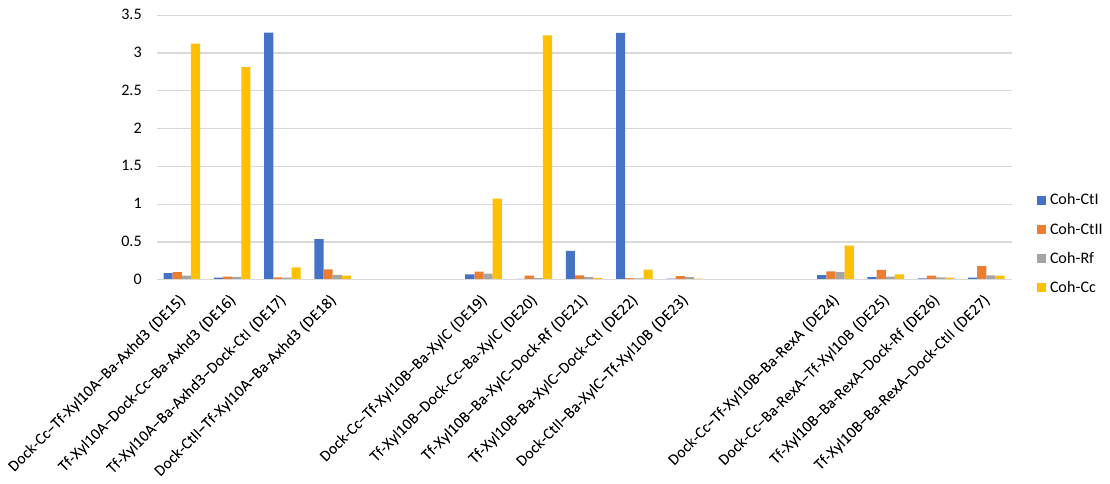

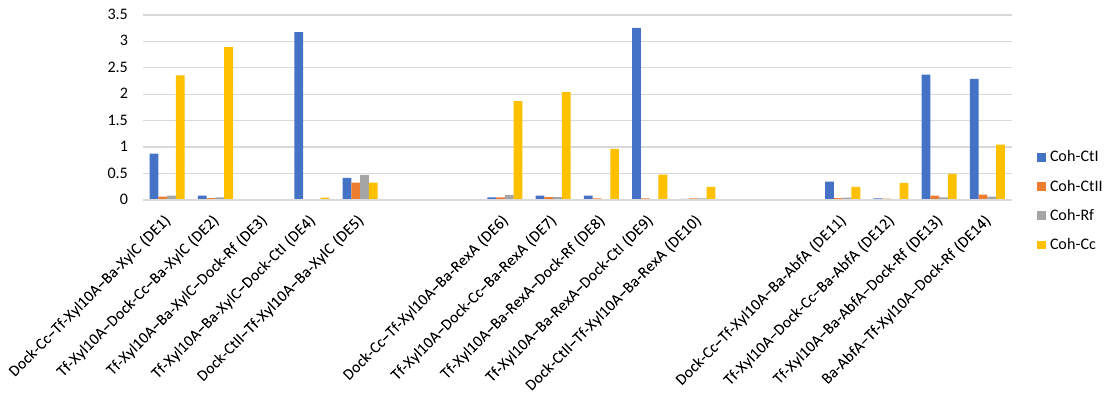

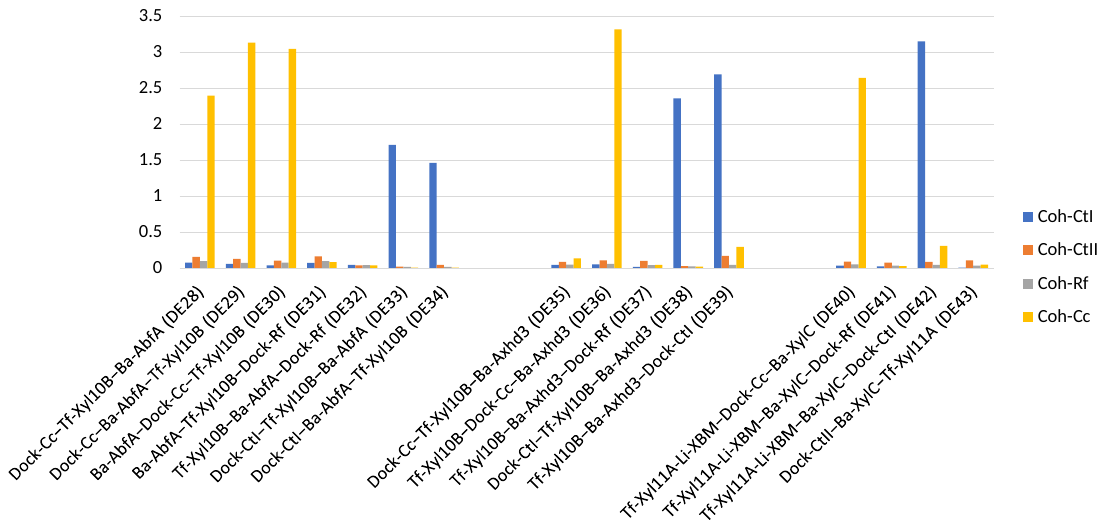

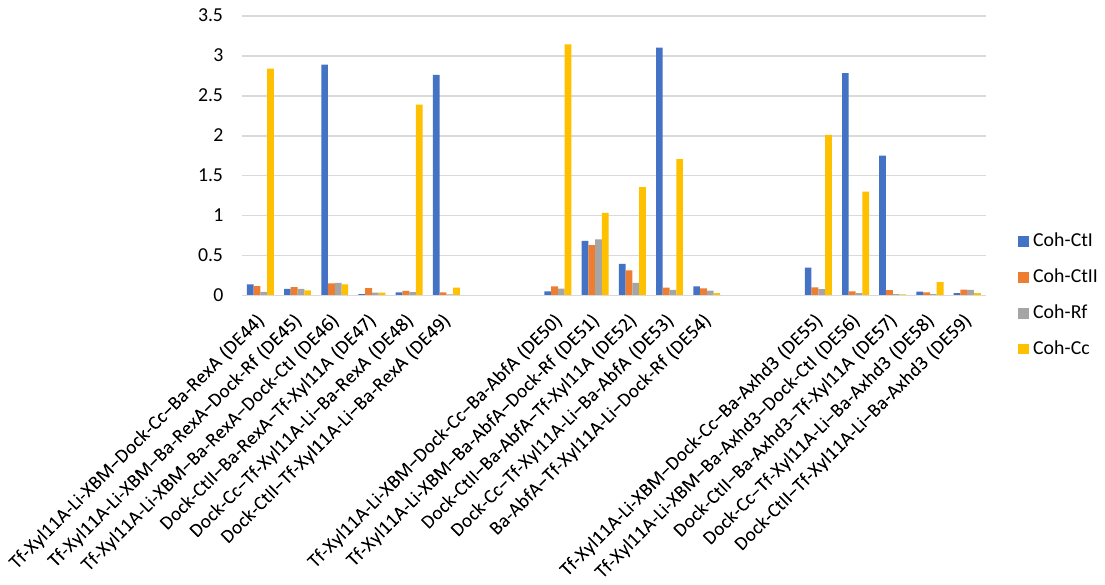

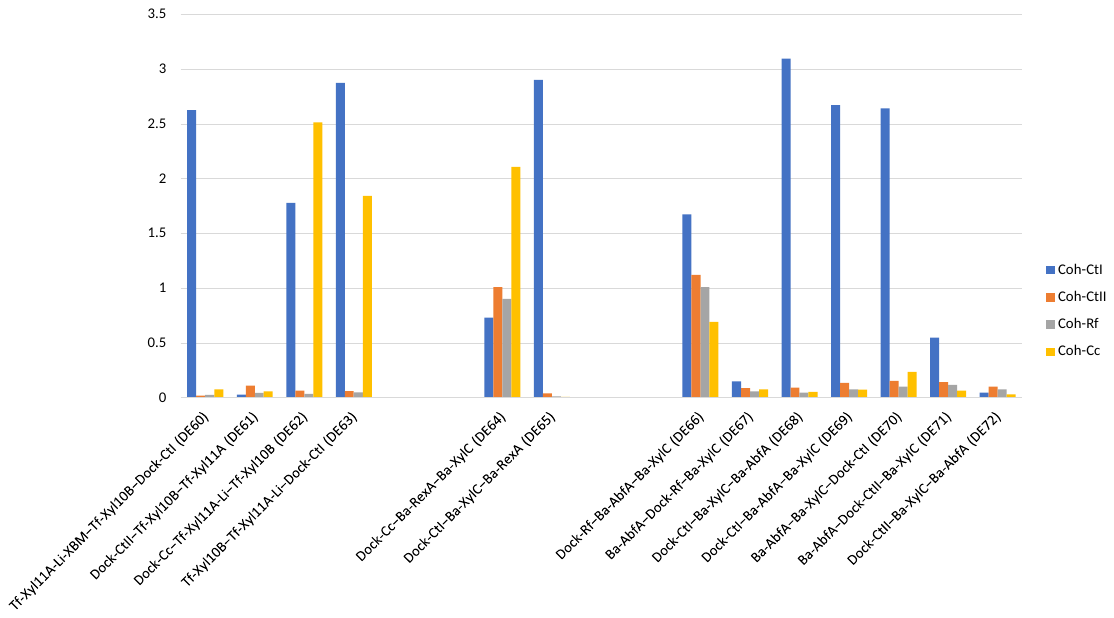

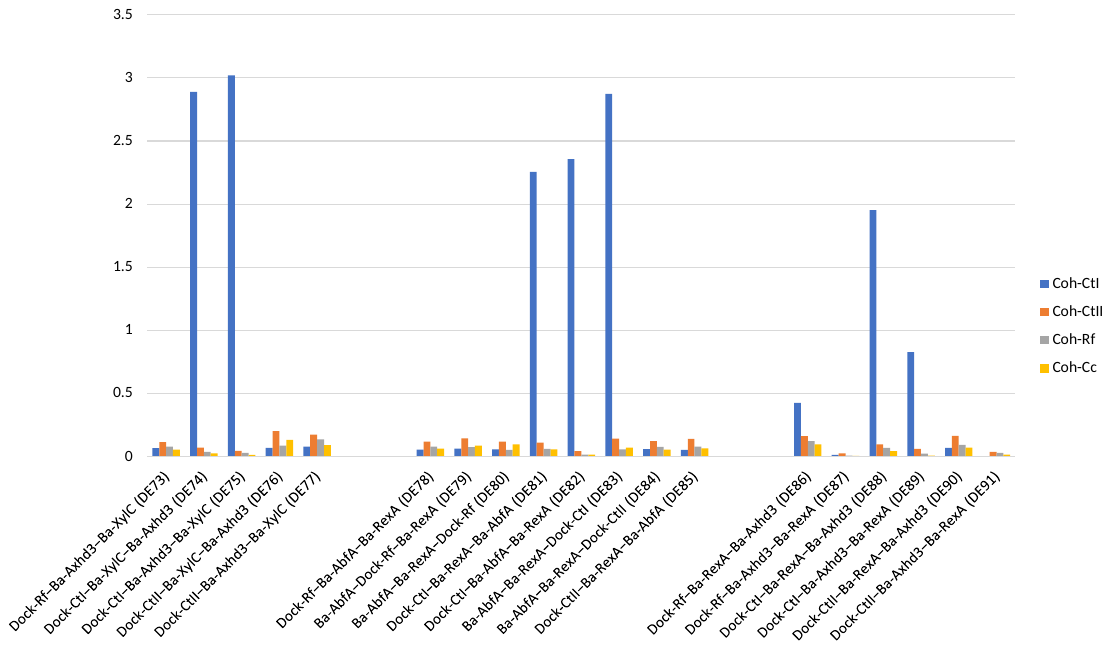

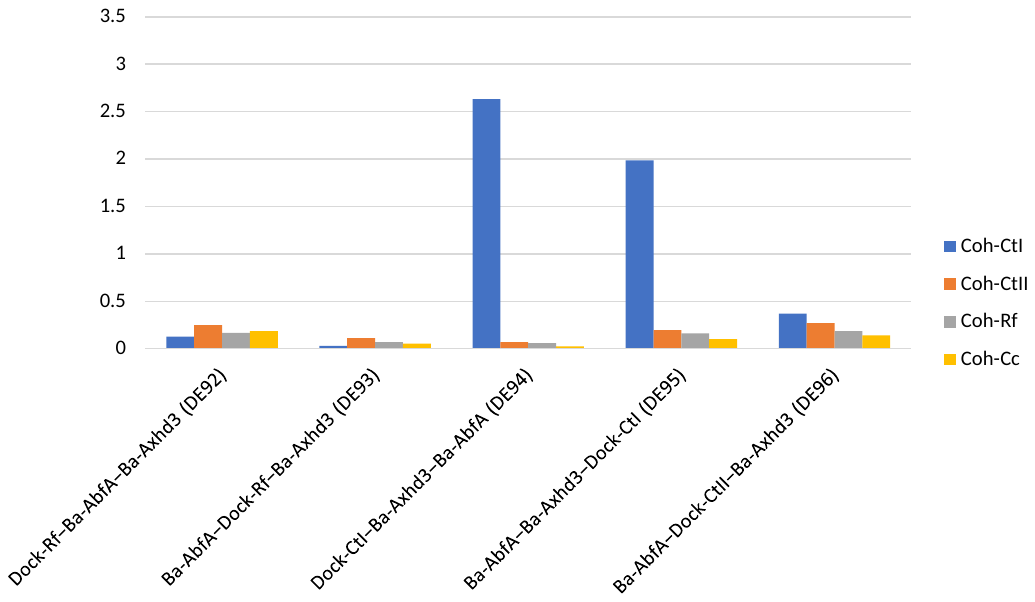
