## Supplementary File S1 for "A funnel-type pipeline for accelerating discovery of tailored designer xylanosomes for efficient arabinoxylan valorization"

**Supplementary File S1: Properties of the AX-active enzymes converted to tiles using VersaTile cloning**

The enzyme modular composition was predicted with InterProScan function. The predicted Glycoside Hydrolase (GH) family (and subfamily in case of GH43 family) or CBM family from the CAZy database is given. GH43-C2 is predicted to be a concanavalin A-like domain, which is typically responsible to bind simple sugars. ND, not determined. Rex, reducing-end xylose-releasing exo-oligoxylanase. AXH, Arabinoxylan arabinofuranohydrolases. *p*NP-X, *p*-Nitrophenyl-β-D-xylopyranoside. *p*NP-X_2_, *p*-Nitrophenyl-β-D-xylobiopyranoside. *p*NP-araf, *p*-Nitrophenyl-α-L-arabinofuranoside. *p*NP-ara, *p*-Nitrophenyl-α-L-arabinopyranoside. *p*NP-G_2_, *p*-Nitrophenyl-β-D-cellobioside

| **Enzyme** | **GenBank ID and domain delineation** | **Enzyme modular composition** | **Optimal temperature** | **Optimal pH** | **Activity reported on (A)XOS/AX** | **Comments** | **Ever used in designer cellulosome?** | **Reference** |
| --- | --- | --- | --- | --- | --- | --- | --- | --- |
| Xyl10A from *Thermobifida fusca*  (*Tf*-Xyl10A) | [AAZ56956.1](https://www.ncbi.nlm.nih.gov/protein/AAZ56956.1)  Residues 39-386 | GH10  CBM2 | 80 °C | 9 | endo-1,4-β-xylanase | - CBM only binds to cellulose - similar substrate specificity as *Tf*-Xyl10B | ^1–3^ | ^4–6^ |
| Xyl10B from *Thermobifida fusca*  (*Tf*-Xyl10B) | [AAZ56824.1](https://www.ncbi.nlm.nih.gov/protein/AAZ56824.1)  Residues 28-399 | GH10 | Room temperature-50 °C | 6 |  | - relatively high binding to insoluble xylan - weak activity on *p*NP-X, *p*NP-X_2_, *p*NP-araf, *p*NP-ara, *p*NP-G_2_ - different substrate specificity than *Tf*-Xyl11A - hydrolyze X_3_ into X_2_ and X | ^1–3,7,8^ | ^6,9^ |
| Xyl11A from *Thermobifida fusca*  (*Tf*-Xyl11A) | [AAA21480.1](https://www.ncbi.nlm.nih.gov/protein/AAA21480.1)  Residues 43-235 | GH11  CBM2 | 50 °C – 65 °C | 5 – 9 |  | - CBM binds to crystalline cellulose and insoluble xylan - weak activity on *p*NP-X_2_ - end-product: X_2_ - degrades both less and heavily branched xylans - hydrolyze X_3_ into X_2_ and X after long reaction times | ^1–3,7,8^ | ^6,9,10^ |
| XylC from *Bifidobacterium adolescentis*  (*Ba*-XylC) | [BAF39209.1](https://www.ncbi.nlm.nih.gov/protein/BAF39209.1)  Residues  1-543 | GH43_11_  GH43-C2 | 50 °C | 6 – 7 | β-xylosidase | - removes xylose units from the non-reducing end of xylan substrates - active on *p*NP-araf and *p*NP-X - weak activity on AX with low degree of substitution - preferred substrate: X_2_ - end product: X | - | ^11,12^ |
| RexA from *Bifidobacterium adolescentis*  (*Ba*-RexA) | [AAO67498.1](https://www.ncbi.nlm.nih.gov/protein/AAO67498.1)  Residues  1-379 | GH8 | 40 °C | 6 | Rex | - removes xylose units from the reducing end of xylan substrates - weak activity on birchwood xylan and oat spelt xylan - preferred substrate: X_3_ - end products: X and X_2_ | - | ^12,13^ |
| AbfA from *Bifidobacterium adolescentis*  (*Ba*-AbfA) | [BAF39204.1](https://www.ncbi.nlm.nih.gov/protein/BAF39204.1)  Residues  1-564 | GH43_12_ | 50 °C | 6 | AXH-m2,3 | - removes *O*-2 and *O*-3 arabinose units from singly substituted xyloses - active on *p*NP-araf and *p*NP-X - active on AX - strongly influenced by heavy arabinose substitution - synergy with *Ba*-Axhd3 | - | ^12,14^ |
| Axhd3 from *Bifidobacterium adolescentis*  (*Ba*-Axhd3) | [AAO67499.1](https://www.ncbi.nlm.nih.gov/protein/AAO67499.1)  Residues  1-529 | GH43_10_  GH43-C2 | 30 °C | 6 | AXH-d3 | - removes *O*-3 arabinose units from doubly substituted xyloses - active on *p*NP-araf - active on AX - not influenced by arabinose substitutions | - | ^12,14^ |
