## Supplementary File S4 for "A funnel-type pipeline for accelerating discovery of tailored designer xylanosomes for efficient arabinoxylan valorization"

**Supplementary File S4: Available tile repository per position**

*Tf*, *Ba*, *Ct*, *Rf* and *Cc* stand for *Thermobifida fusca*, *Bifidobacterium adolescentis*, *Clostridium thermocellum*, *Ruminococcus flavefaciens* and *Clostridium cellulolyticum*, respectively*.* Accession numbers are indicated. NA – Not applicable. *Tf*-Xyl11A-Li is not interesting to have at C-terminus position (PT_3_-PT_e_) as the linker can negatively impact the expression/stability of the overall protein. *Tf*-Xyl11A-Li-XBM was not designed for PT_2_-PT_3_ position as we hypothesize that placing the XBM at an internal position could impact negatively the overall efficiency of the DC. Similarly, Stern at al. have seen that a higher activity was obtained when the CBM was placed at the *N*-terminus of the scaffoldin and not internally. Some tiles were not completed due to recurrent technical failures. A recurrent mistake was seen for *Ba*-XylC, *Ba*-RexA and *Ba*-Axhd3 tiles at the PT_s_-PT_2_ position (deletion in PT_2_). The construction of *Tf*-Xyl11A and *Tf*-Xyl11A-Li tiles at PT_s_-PT_2_ position was not successful during the mutagenesis procedure to remove internal BsaI recognition sites. The *Tf*-Xyl11A-Li-XBM tile at position PT_3_-PT_e_ failed upon PCR amplification.

| **Enzyme/Dockerin** | | | **Position** | | |
| --- | --- | --- | --- | --- | --- |
| GenBank ID | Domain delineation  (AA residues) | Schematic representation | PT_s_-PT_2_ | PT_2_-PT_3_ | PT_3_-PT_e_ |
| *Tf*-Xyl10A  [AAZ56956.1](https://www.ncbi.nlm.nih.gov/protein/AAZ56956.1) | 39-361/386 | 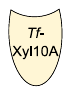 | X | X | X |
| *Tf*-Xyl10B  [AAZ56824.1](https://www.ncbi.nlm.nih.gov/protein/AAZ56824.1) | 28-399 | 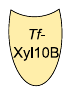 | X | X | X |
| *Tf*-Xyl11A  [AAA21480.1](https://www.ncbi.nlm.nih.gov/protein/AAA21480.1) | 43-235 | 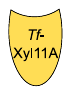 |  | X | X |
| *Tf*-Xyl11A-Li  [AAA21480.1](https://www.ncbi.nlm.nih.gov/protein/AAA21480.1) | 43-245 | 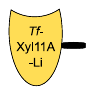 |  | X | NA |
| *Tf*-Xyl11A-Li-XBM  [AAA21480.1](https://www.ncbi.nlm.nih.gov/protein/AAA21480.1) | 43-338 | 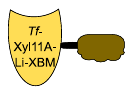 | X | NA |  |
| *Ba*-XylC  [BAF39209.1](https://www.ncbi.nlm.nih.gov/protein/BAF39209.1) | 1-543 | 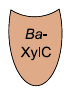 |  | X | X |
| *Ba*-RexA  [AAO67498.1](https://www.ncbi.nlm.nih.gov/protein/AAO67498.1) | 1-379 | 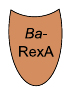 |  | X | X |
| *Ba*-AbfA  [BAF39204.1](https://www.ncbi.nlm.nih.gov/protein/BAF39204.1) | 1-564 | 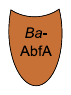 | X | X | X |
| *Ba*-Axhd3  [AAO67499.1](https://www.ncbi.nlm.nih.gov/protein/AAO67499.1) | 1-529 | 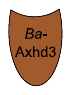 |  | X | X |
| Dock-*Ct*I  [ABN53296.1](https://www.ncbi.nlm.nih.gov/protein/ABN53296.1) | 2-75 | 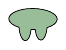 | X | X | X |
| Dock-*Ct*II  [Q06851](https://www.uniprot.org/uniprot/Q06851) | 1702-1853 | 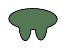 | X | X | X |
| Dock-*Rf*  [CAC83072.1](https://www.ncbi.nlm.nih.gov/protein/CAC83072.1) | 728-808 | 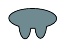 | X | X | X |
| Dock-*Cc*  [AAA23221.1](https://www.ncbi.nlm.nih.gov/protein/AAA23221.1) | 409-475 | 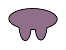 | X | X | X |
