## Supplementary File S5 for "A funnel-type pipeline for accelerating discovery of tailored designer xylanosomes for efficient arabinoxylan valorization"

**Supplementary File S5: Cognate dockerin-cohesin pairs used in this study**

Dockerin and cohesin tiles designed and constructed by Hans Gerstmans and Boris Bekaert, respectively, were available in the tile repository. Dockerins are available for the 3-way assembly (PT_s_-PT_2_, PT_2_-PT_3_ and PT_3_-PT_e_) allowing the construction of DEs with three separate domains like one dockerin and two catalytic domains as in this chapter. Cohesins are available for the 5-way assembly (PT_s_-PT_2_, PT_2_-PT_3_, PT_3_-PT_4_, PT_4_-PT_5_ and PT_5_-PT_e_) allowing the construction of scaffoldins with five separate domains like a pentavalent scaffoldin with five cohesins or a tetravalent scaffoldin with four cohesins and one CBM. Design of dockerin/cohesin amplification primers was done based on complementary DNA sequences used in Haimovitz et al. Natural linkers at the *C*-terminus of the cohesins were included in all positions except for position PT_5_-PT_e_ to avoid scaffoldins expression/stability hitches. Accession numbers are given underneath the name of each sequence. The original protein and host of the dockerin/cohesin sequences is described. All dockerins are *C*-terminal dockerins. A figure with a specific colour for each dockerin/cohesin pair is given and is used throughout this dissertation. Originally the dockerin-cohesin interactions were divided into type I, II and III based on sequence homology and parent protein identity. Type I and II were first described in *Clostridium thermocellum* and type III in *Ruminococcus flavefaciens.* Due to the discovery of divergent dockerin-cohesin sequences and intriguing cellulosome structures, it is likely this classification will be updated. ^a^ To the extent of our knowledge, the dissociation constant (K_D_) between dockerin from scaffoldin ScaA and cohesin n°4 of the ScaB anchoring protein from *Clostridium thermocellum* had not been determined. The K_D_ presented here as a reference corresponds to the affinity between dockerin from scaffoldin ScaA and cohesin from ScaF scaffoldin depending if X module is incorporated or not. ^b^ To the extent of our knowledge, the K_D_ for this pair used in this work has not been determined. The K_D_ presented here as a reference corresponds to a *C*-terminal dockerin from a family 9 GH and a family 12 carbohydrate esterase and ScaA from *Ruminococcus flavefaciens*.

| **Dockerins** | | | | **Cohesins** | | | | | **Type of interaction** | **Constant of dissociation (K_D_)** | **Source** |
| --- | --- | --- | --- | --- | --- | --- | --- | --- | --- | --- | --- |
| **Name** | **Domain dealineation** | **Figure** | **Description** | **Name** | **Domain dealineation** | **Figure** | **Description** | **Observations** |  |  |  |
| Dock-*Ct*I  [ABN53296.1](https://www.ncbi.nlm.nih.gov/protein/ABN53296.1) | 2-75 | 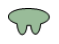 | Dockerin from **GH48** cellobiohydrolase **Cel48S** (*Clostridium thermocellum*) | Coh-*Ct*I  [CCV01465.1](https://www.ncbi.nlm.nih.gov/protein/CCV01465.1) | 182-363 | 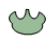 | Cohesin n°2 of the **ScaA** scaffoldin (*Clostridium thermocellum*) | **ScaA** contains a CBM between the 2^nd^ and 3^rd^ of 9 repeated cohesin domains, a type II dockerin domain  at its C-terminal and a X-module | Type-I | > 10^-11^ M | [1,2] |
| Dock-*Ct*II  [Q06851](https://www.uniprot.org/uniprot/Q06851) | 1702-1853 | 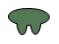 | Dockerin from scaffoldin **CipA** (*Clostridium thermocellum*) | Coh-*Ct*II  [WP_059169945.1](https://www.ncbi.nlm.nih.gov/protein/WP_059169945.1) | 401-779 | 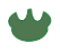 | Cohesin n°4 of the **ScaB** anchoring protein (*Clostridium thermocellum*) | **ScaB** is an anchoring protein that contains a SLH domain that attaches the protein to the cell surface and 4 type-II cohesins^269^ | Type-II | ~10^-9^-10^-10^ M^a,^ | [3,4] |
| Dock-*Rf*  [CAC83072.1](https://www.ncbi.nlm.nih.gov/protein/CAC83072.1) | 728-808 | 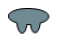 | Dockerin from **GH44** cellulase **EndB** (*Ruminococcus flavefaciens* 17)^274^ | Coh-*Rf*  [CAC34384.3](https://www.ncbi.nlm.nih.gov/protein/CAC34384.3) | 447-788 | 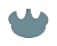 | Cohesin n°3 of the **ScaA** scaffoldin (*Ruminococcus flavefaciens* 17) | **ScaA** contains 3 cohesins and 1 dockerin that interacts with cohesin of scaffoldin ScaB | Type-III | ~10^-7^-10^-8^ M^b^ | [5,6] |
| Dock-*Cc*  [AAA23221.1](https://www.ncbi.nlm.nih.gov/protein/AAA23221.1) | 409-475 | 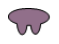 | Dockerin from **GH5** cellulase **Cel5A** (*Clostridium cellulolyticum*) | Coh-*Cc*  [AAC28899.2](https://www.ncbi.nlm.nih.gov/protein/AAC28899.2) | 288-434 | 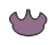 | Cohesin n°1 of the **CipC** scaffoldin (*Clostridium cellulolyticum*) | **CipC** contains a CBM and 8 cohesin domains with X2 domains in between CBM and 1^st^ cohesin and 7^th^ and 8^th^ cohesins | Type-I | 2.5×10^-10^ M | [7] |
