## Supplementary File S6 for "A funnel-type pipeline for accelerating discovery of tailored designer xylanosomes for efficient arabinoxylan valorization"

**Supplementary File S6: *Ct*-CBM3a peptide sequence used for rabbit immunization to generate the antiserum used in ELISA**

>ANTPVSGNLKVEFYNSNPSDTTNSINPQFKVTNTGSSAIDLSKLTLRYYYTVDGQKDQTFWCDHAAIIGSNGSYNGITSNVKGTFVKMSSSTNNADTYLEISFTGGTLEPGAHVQIQGRFAKNDWSNYTQSNDYSFKSASQFVEWDQVTAYLNGVLVWGKEPGGSVV*
