## Supplementary File S7 for "A funnel-type pipeline for accelerating discovery of tailored designer xylanosomes for efficient arabinoxylan valorization"

**Supplementary File S7: Number of possible AX-active DEs created by the VersaTile platform**

A) Number of dockerin tiles (
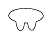
) used for this study and AX-active enzymes (
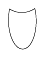
) added to the VersaTile repository. B) Number of possible bicatalytic dockerin containing enzymes (DEs) active on AX constructed by VersaTile based on the number of tiles used.

| **A: Number of tiles used from the repository** | | | | | | **B: Number of possible bicatalytic DEs** | | | |
| --- | --- | --- | --- | --- | --- | --- | --- | --- | --- |
| **PT_s_-PT_2_** | | **PT_2_-PT_3_** | | **PT_3_-PT_e_** | | 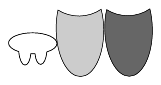 | 224 | 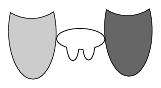 |  |
|  |  |  |  |  |  | 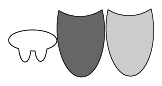 |  | 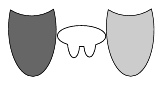 | 112 |
| 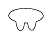 | 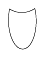 | 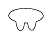 | 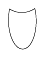 | 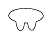 | 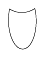 | 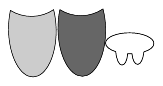 | 128 | **Total n° of possible bicatalytic DE** | **464** |
| 4 | 4 | 4 | 8 | 4 | 7 | 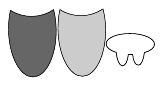 |  |  |  |
