## Supplementary File S10 for "A funnel-type pipeline for accelerating discovery of tailored designer xylanosomes for efficient arabinoxylan valorization"

**Supplementary File S10: Dot blot and Western blot results to determine the volumes of DE and scaffoldin lysates to be used for DX assembly**

*Section 1/3 – SDS-PAGE (A) and Western Blot (B) results of DEs and scaffoldin lysates*

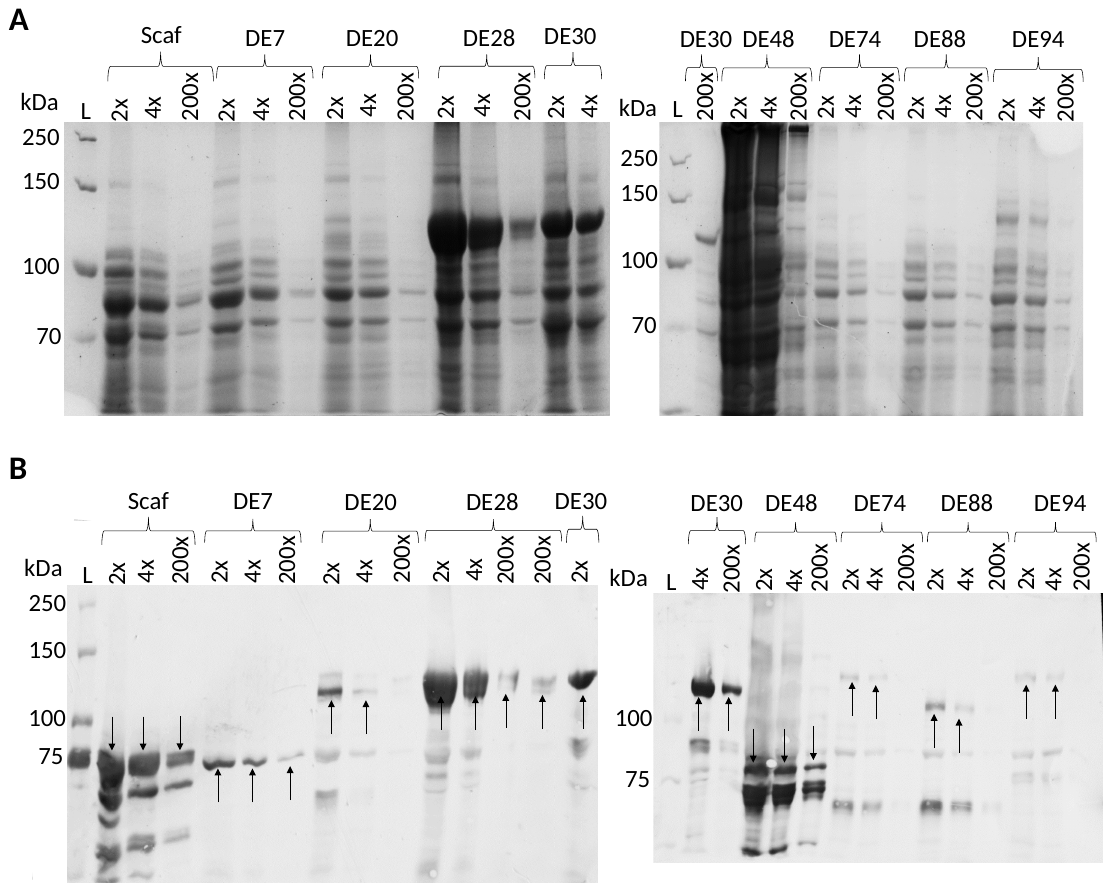

Figure S10.1 – SDS-PAGE (A) and WB (B) of lysates of DEs and scaffoldin to be used to construct DXs. All proteins were expressed using *E. coli* BL21 (DE3) CodonPlus RIL, except DE48 which was expressed by *E. coli* Rosetta™ (DE3) pLysS as DE48 was not expressed by *E. coli* BL21 (DE3) CodonPlus RIL when Terrific broth (TB) was used. Scaffoldin (Scaf, 84 kDa), DE7 (90 kDa), DE20 (113 kDa), DE28 (113 kDa), DE30 (113 kDa), DE48 (75 kDa), DE74 (131 kDa), DE88 (112 kDa) and DE94 (131 kDa) were expressed in parallel in 5 mL TB to boost expression yields, lyzed with BugBuster Master mix, diluted 2, 4 and 200x and loaded on 8% acrylamide gels for SDS-PAGE and WB analysis. The black arrows in WB indicate bands at the expected MW for intact proteins. The PageRuler unstained broad range protein ladder (Thermo Fisher Scientific) and the Precision Plus Protein™ Dual Color (Bio-Rad) standards were used for SDS-PAGE and WB, respectively.

*Section 2/3 – ImageJ peak areas of the dot blot assay performed with DE and scaffoldin lysates for DX assembly.*

Duplicates of two microliters of undiluted, 2x, 5x, 10x and 20x diluted scaffoldin and DE samples were analyzed by dot blot. In green, the values used to estimate relative amounts of scaffoldin and DEs. A limitation of the approach is a considerable variation in peak area values for the replicates due to the limited quantitative power of dot plots. We tolerate these values as rough estimators to accommodate the need for a high-throughput approach for DX assembly.

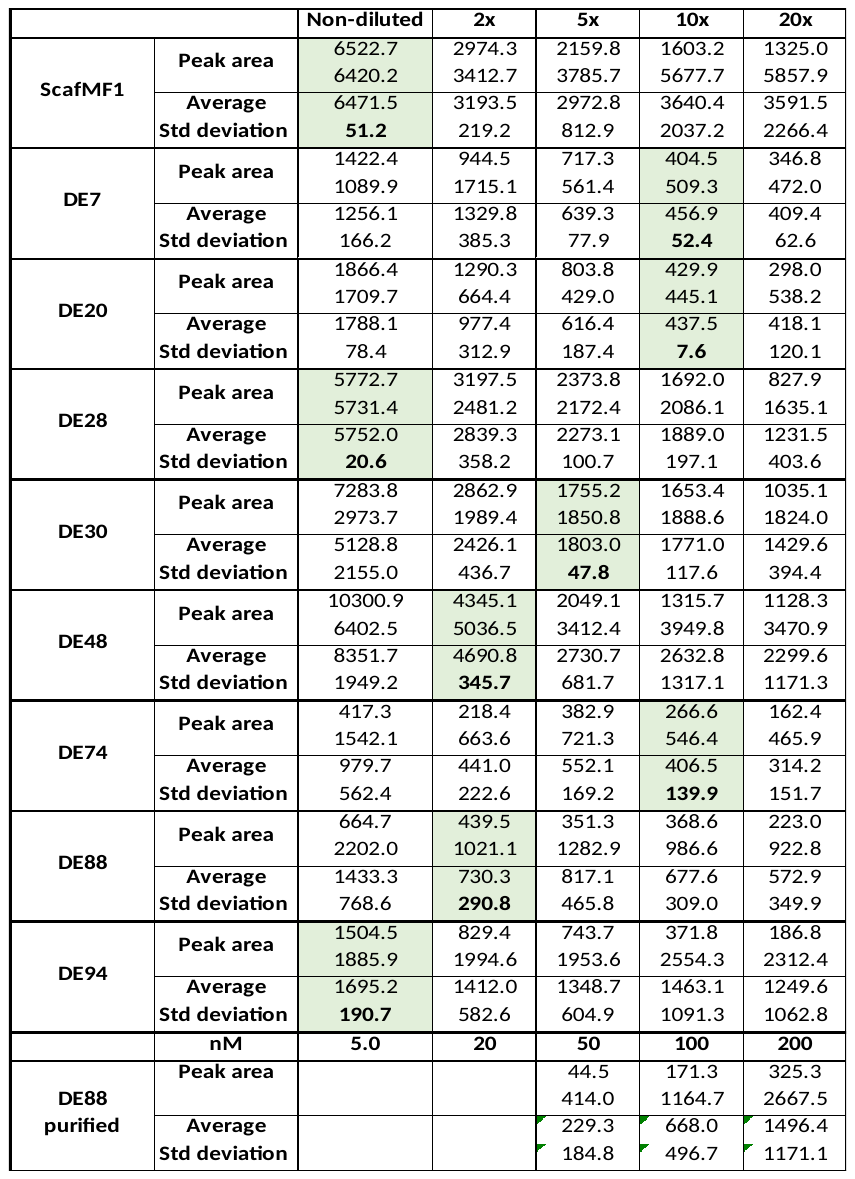

*Section 3/3 – Volumes of DE and scaffoldin lysates used to assemble the DXs.*

The scaffoldin used contains from *N*- to *C*- terminus a glutathione S-transferase (GST) tag, a *C. thermocellum* ScaA CBM from CAZy database family 3 (*Ct*-CBM3a), the cohesin n°2 of the ScaA scaffoldin (*C. thermocellum*), the cohesin n°1 of the CipC scaffoldin (*C. cellulolyticum*) and a His_6_ tag (Coh-*Ct*I−Coh-*Cc*). The DEs contain 1 dockerin (Dock-*Ct*I or Dock-*Cc*), 2 catalytic domains and 1 His_6_ tag at the *C*-terminus. DX1-8 are bivalent DXs containing 1 DE. These are controls to assess the affinity of DEs to the bivalent scaffoldin used. DX9-23 are bivalent DXs containing 2 bicatalytic DEs. C1-9 are control samples containing only DEs or scaffoldin. Per DX, the volume of DE and scaffoldin lysate is indicated. While the scaffoldin lysate volume was kept constant, an excess amount of DE lysate was added based on semi-quantitative assessment of the western blot and dot plot analysis. An adjusted buffer volume was added to obtain a final volume of 500 µL.

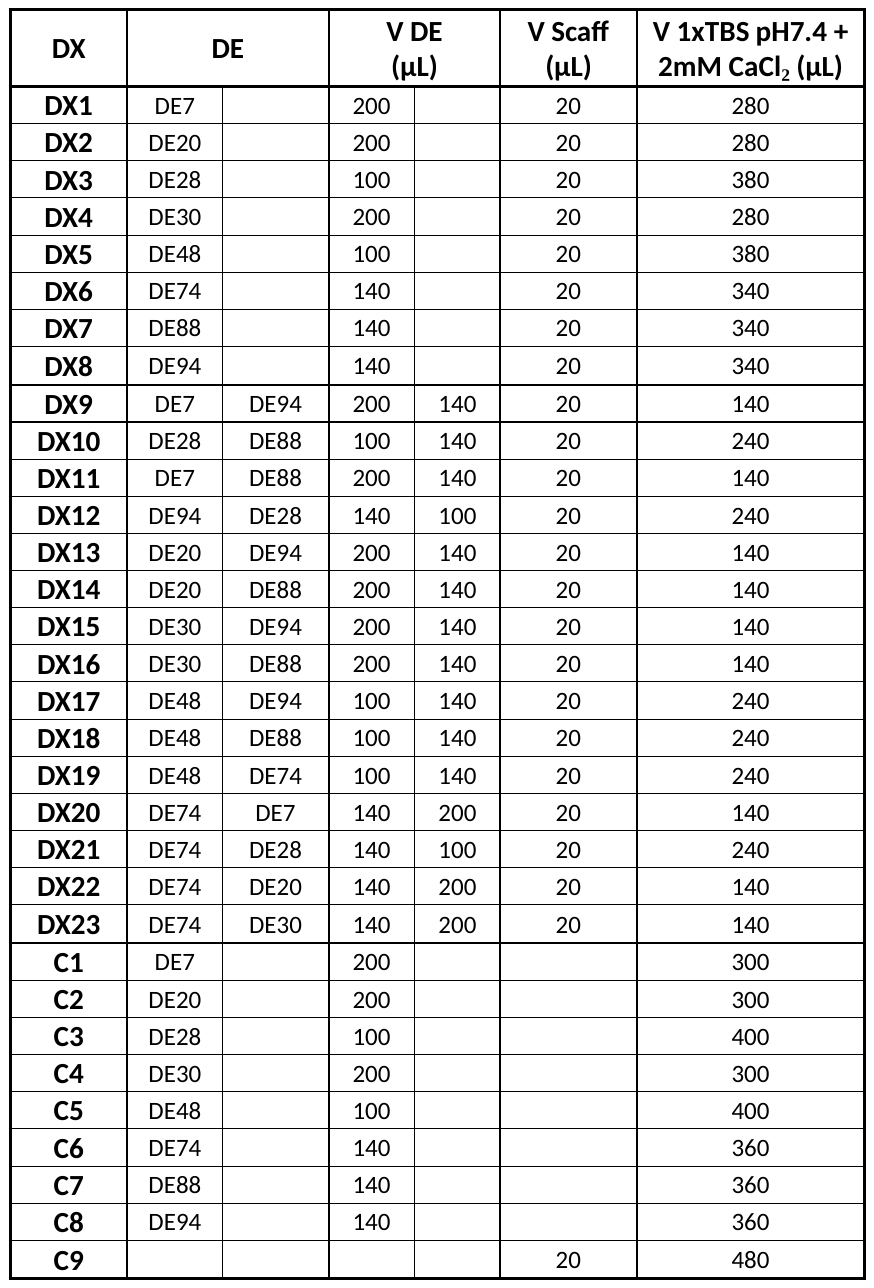
