## Supplementary File S11 for "A funnel-type pipeline for accelerating discovery of tailored designer xylanosomes for efficient arabinoxylan valorization"

c)

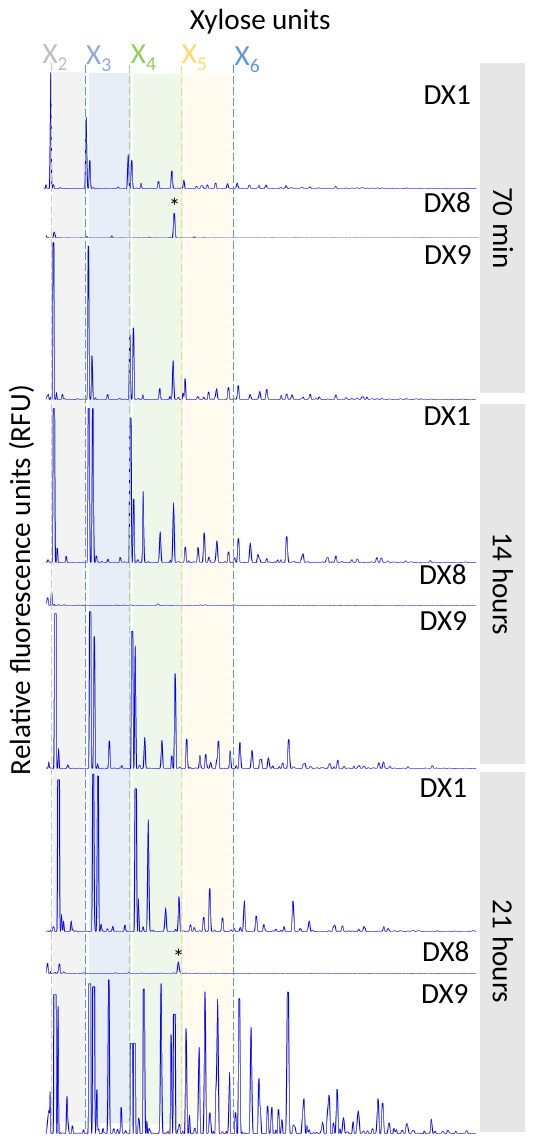

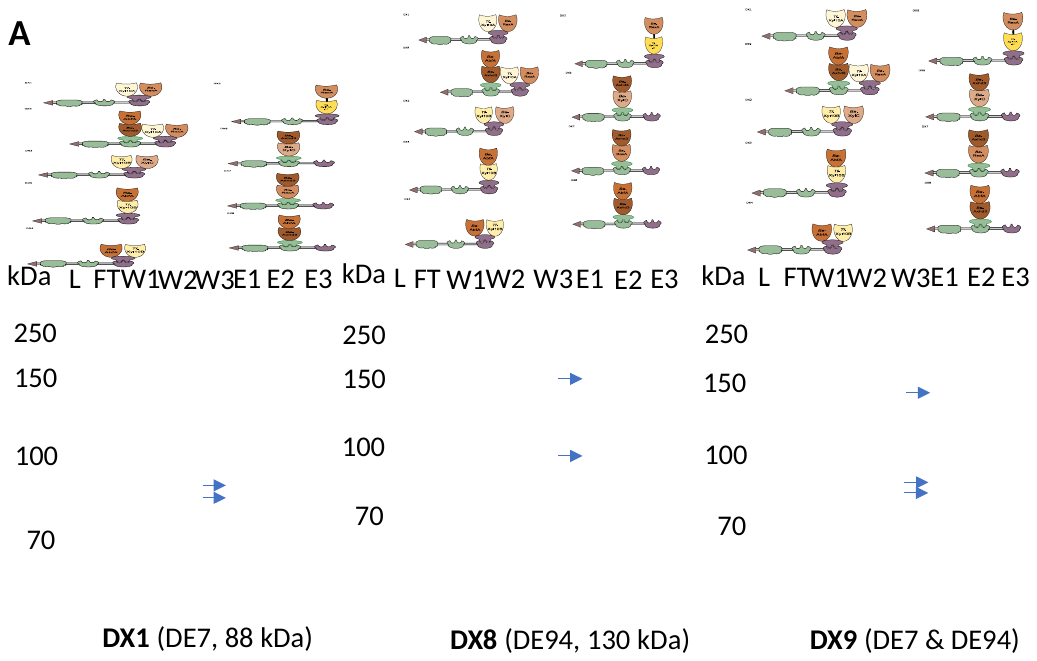

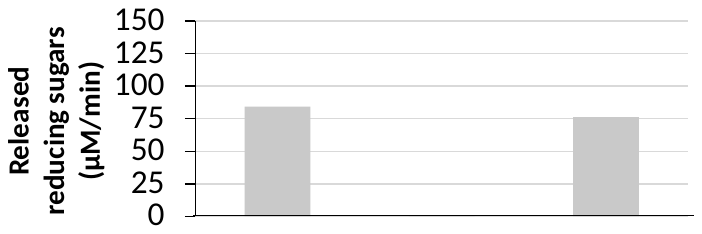

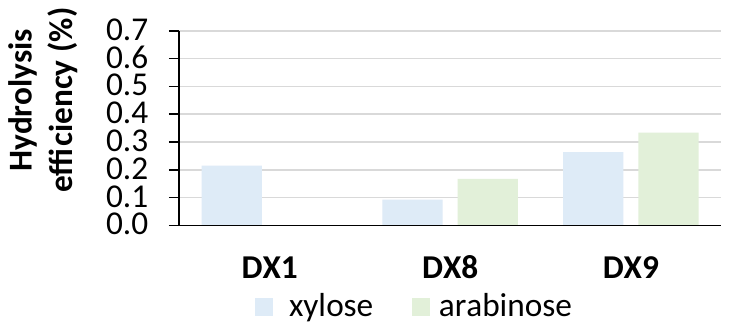

a)

b)

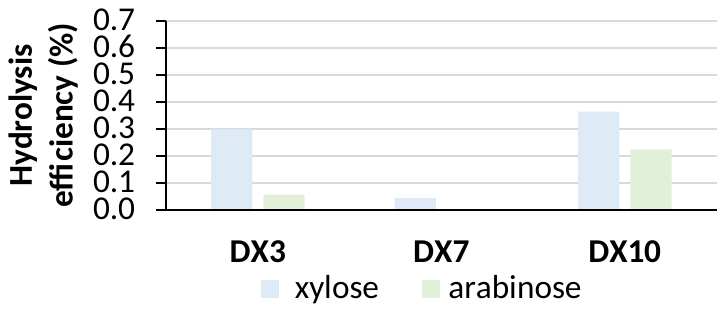

c)

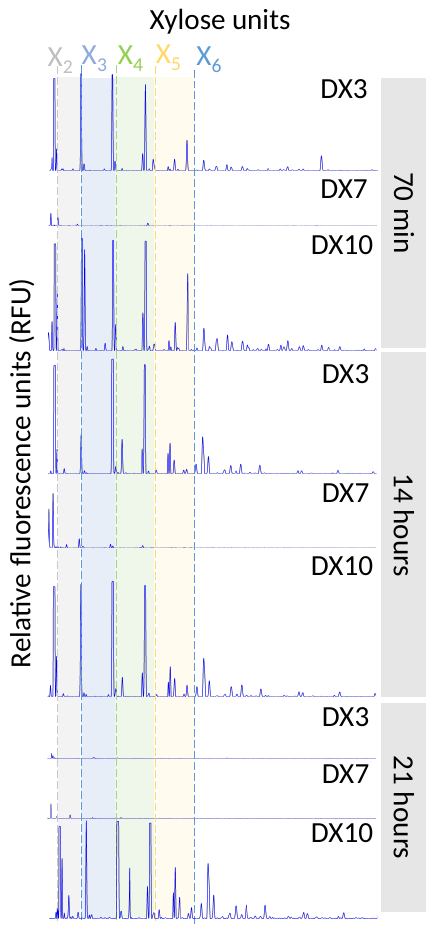

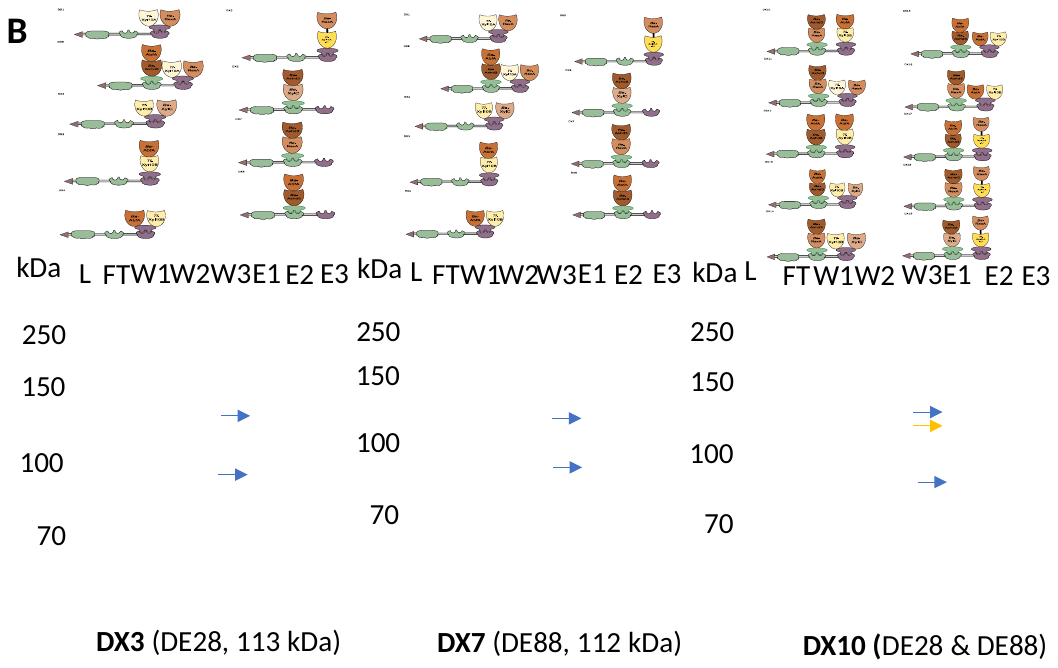

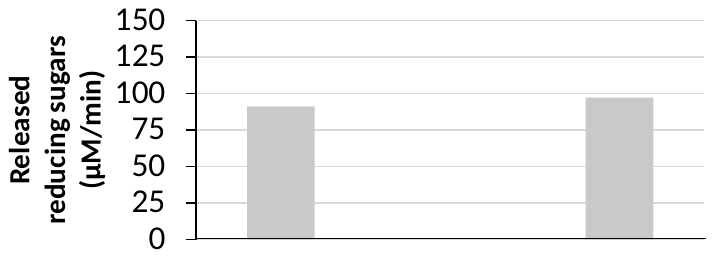

a)

b)

c)

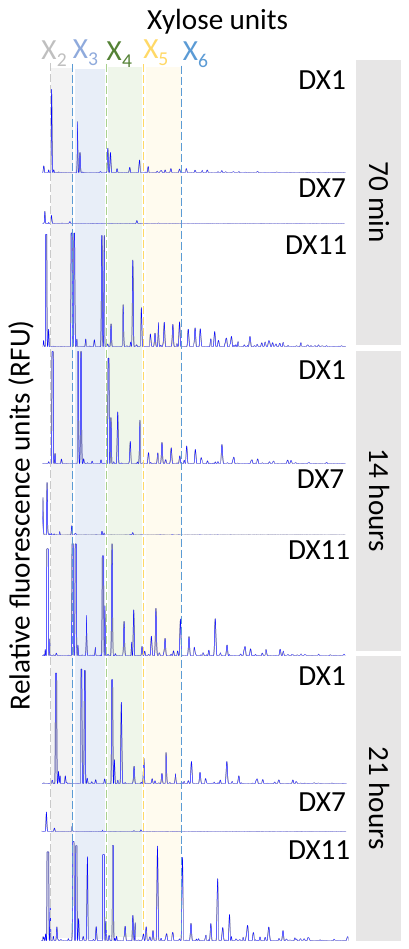

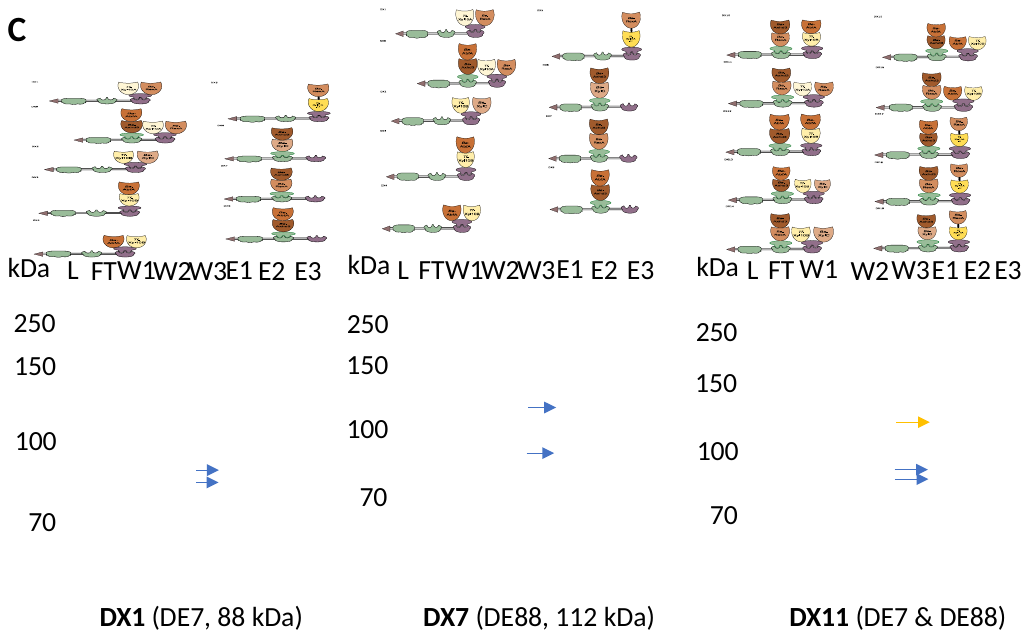

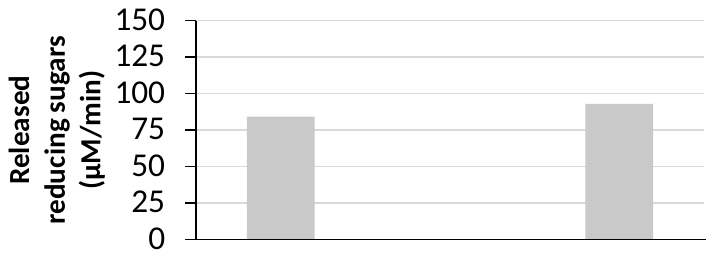

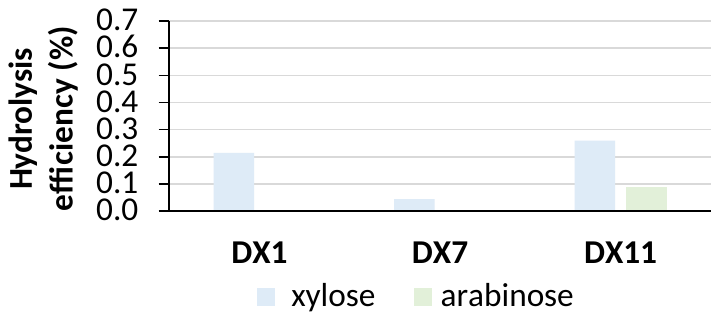

b)

a)

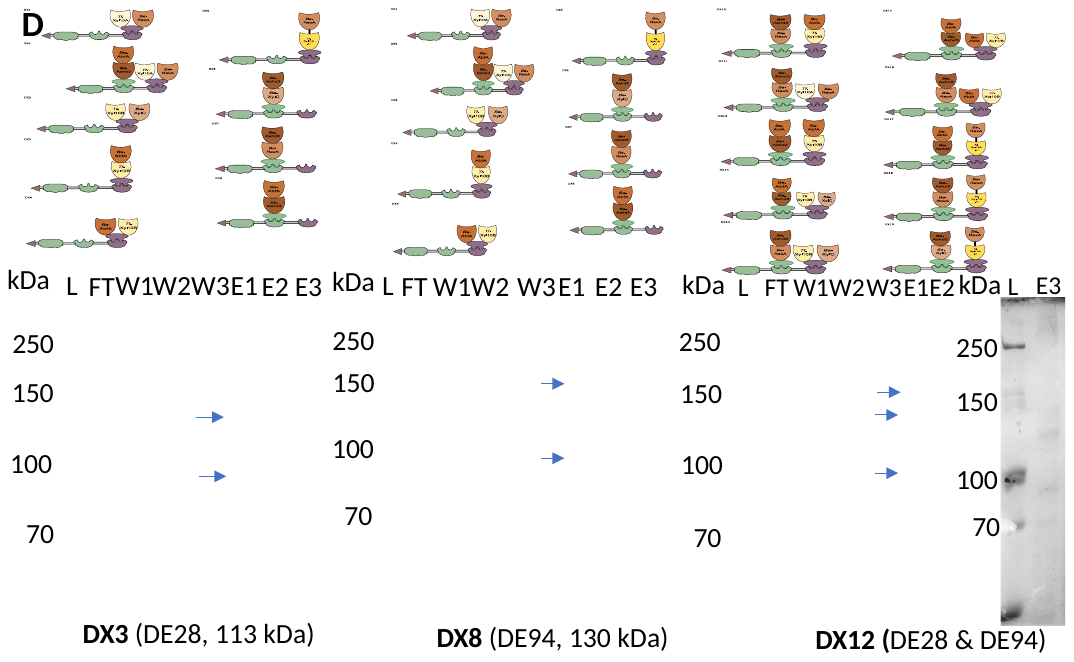

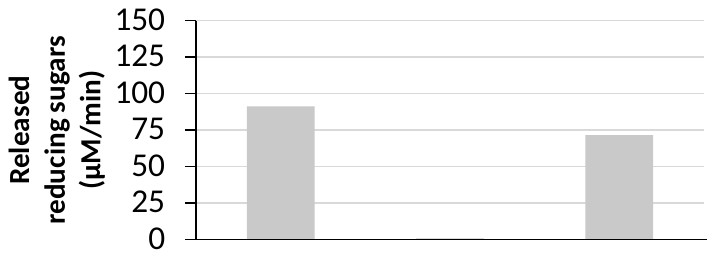

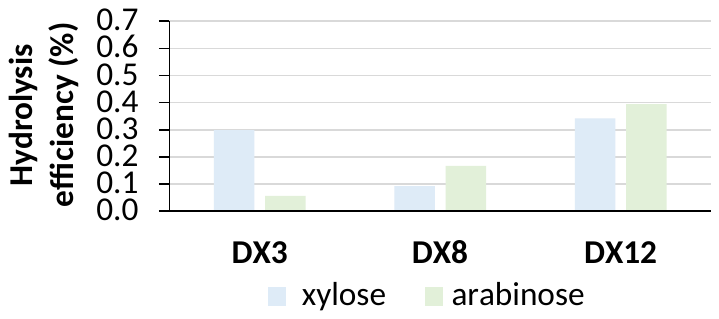

a)

b)

c)

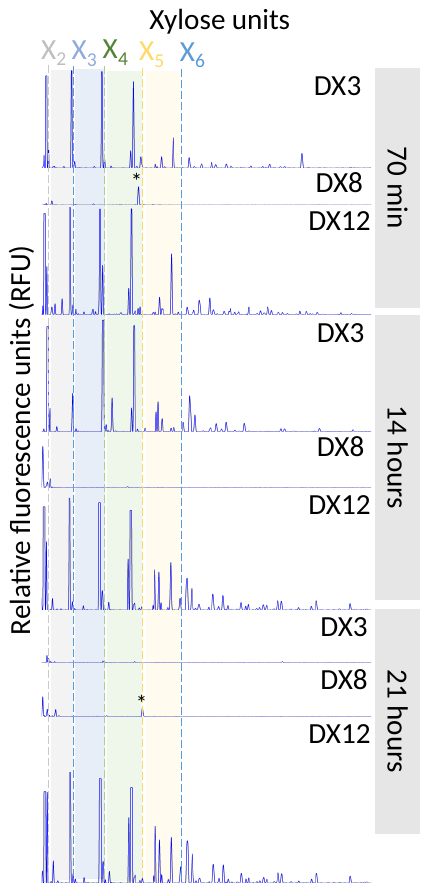

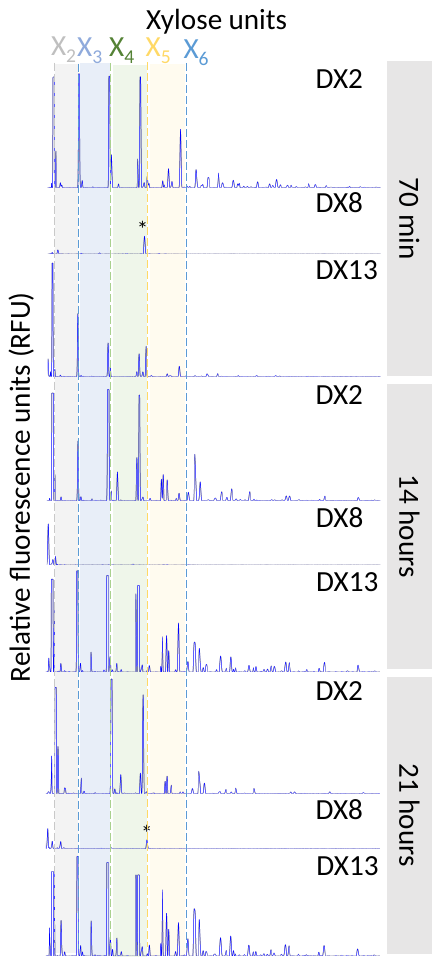

c)

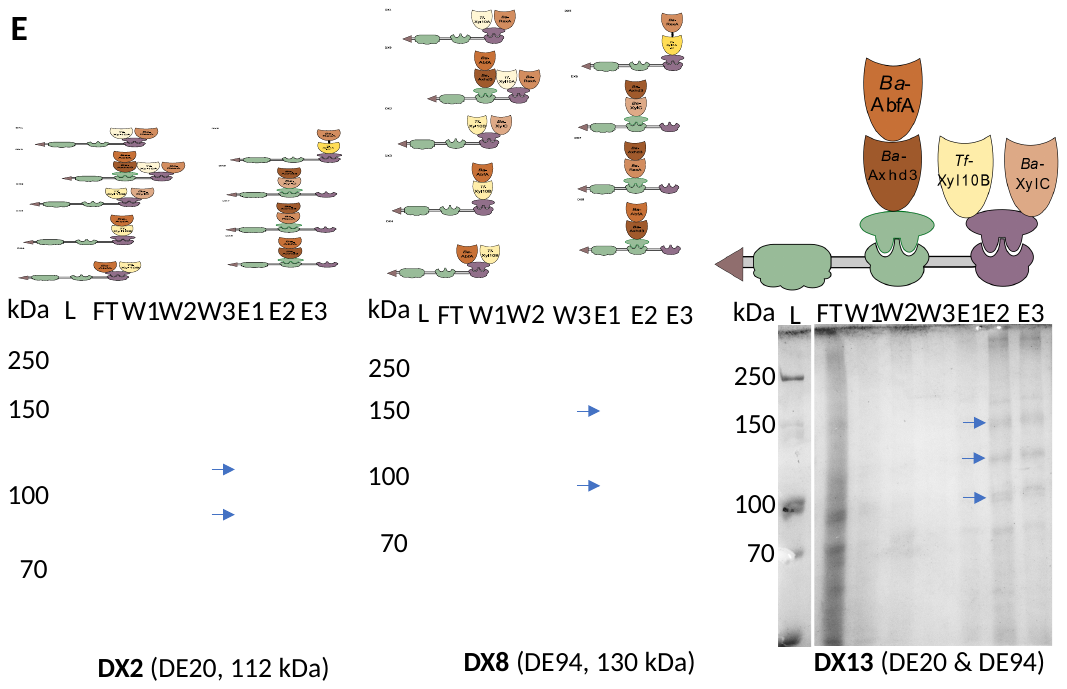

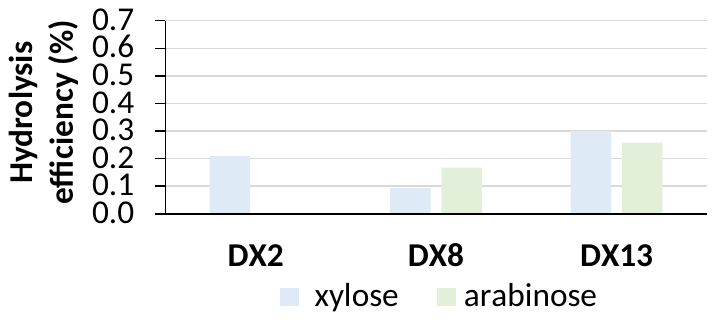

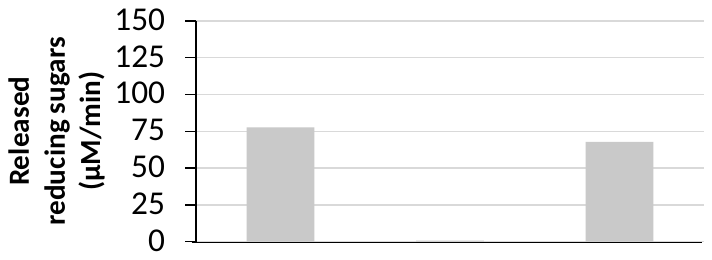

a)

b)

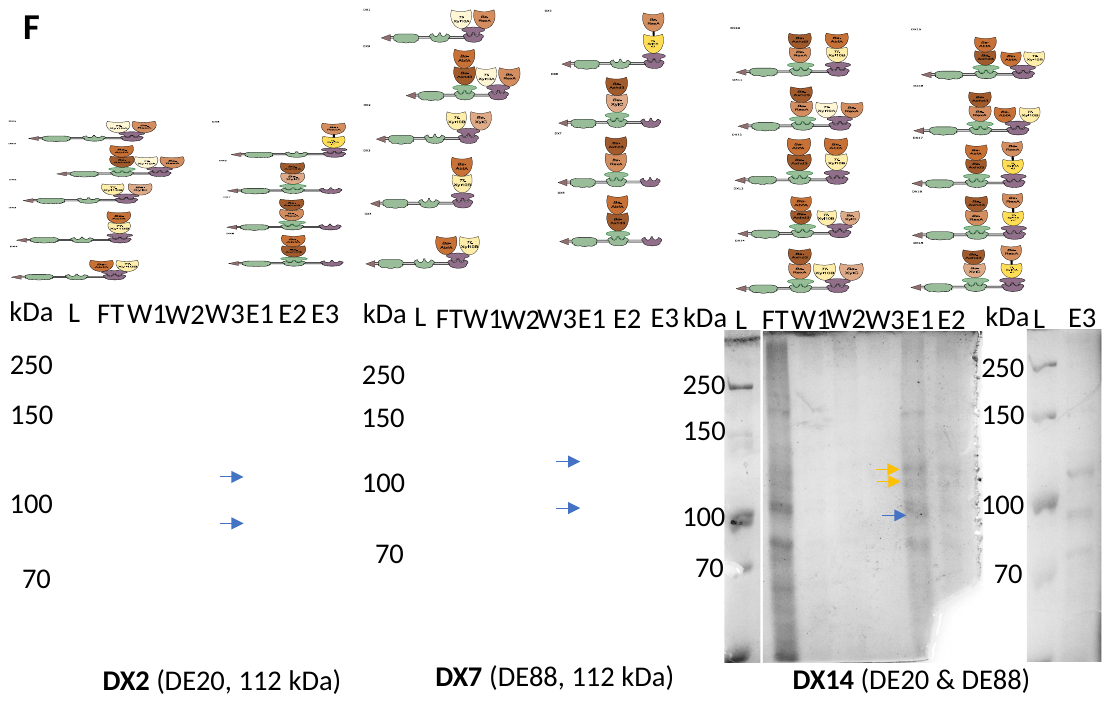

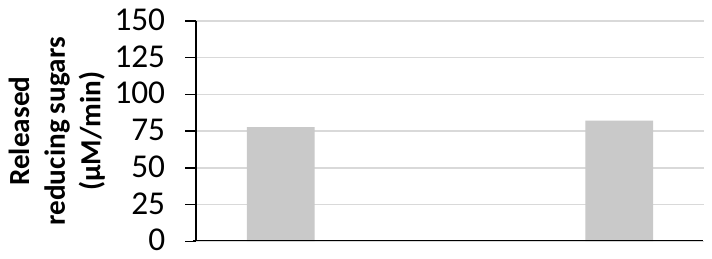

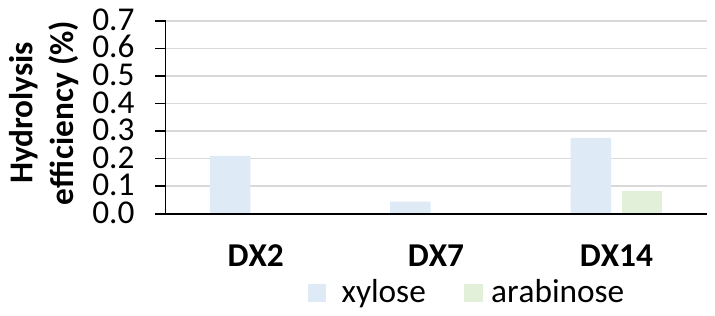

a)

b)

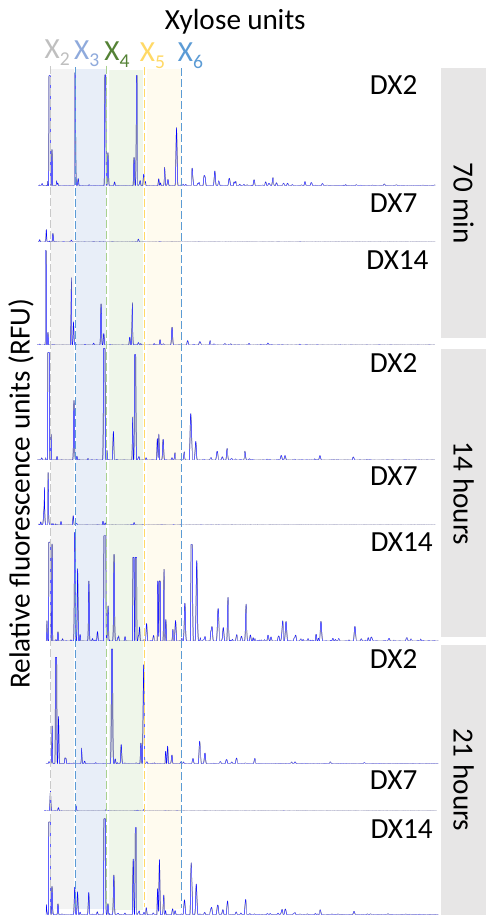

c)

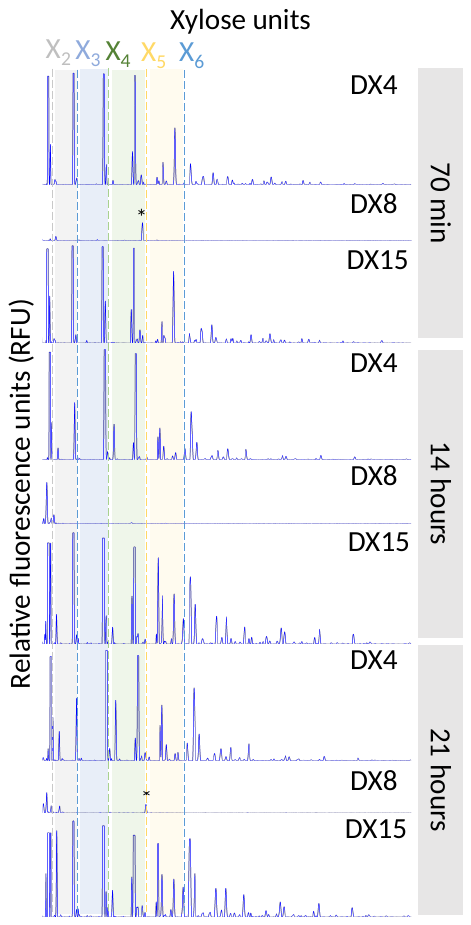

c)

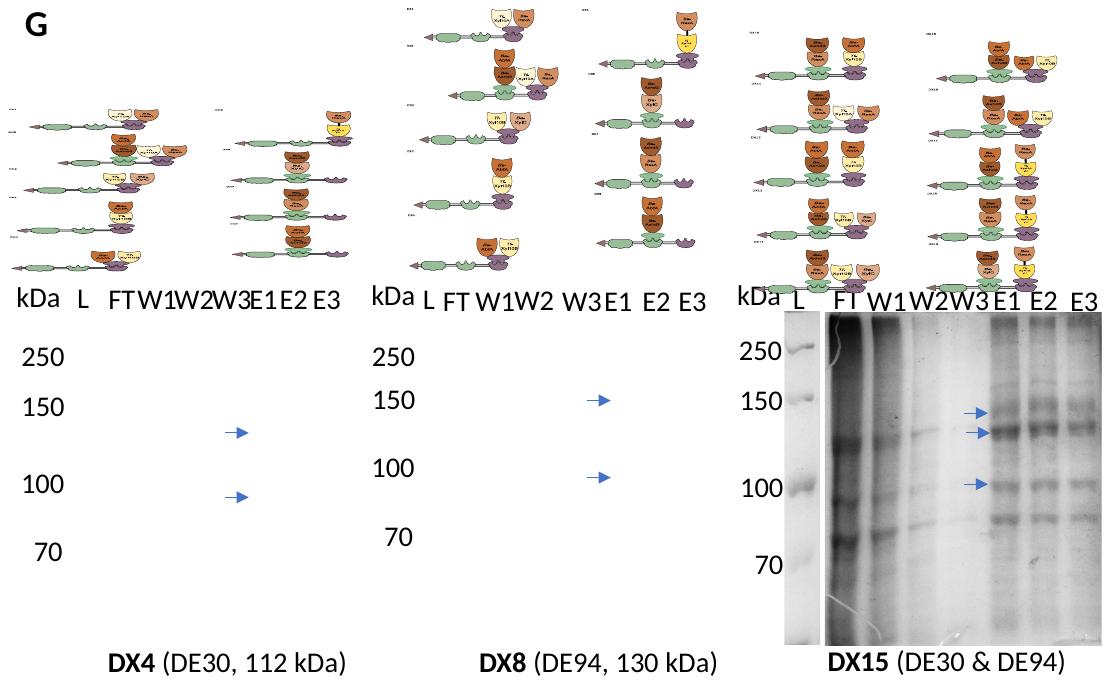

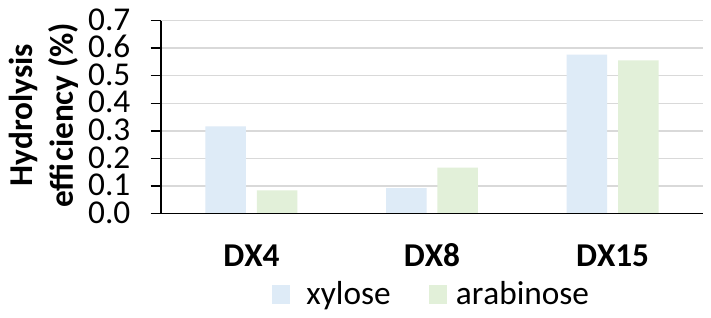

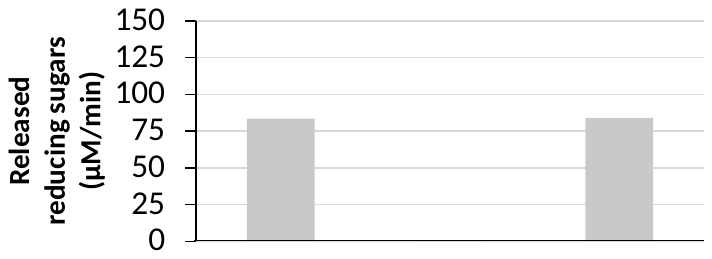

a)

b)

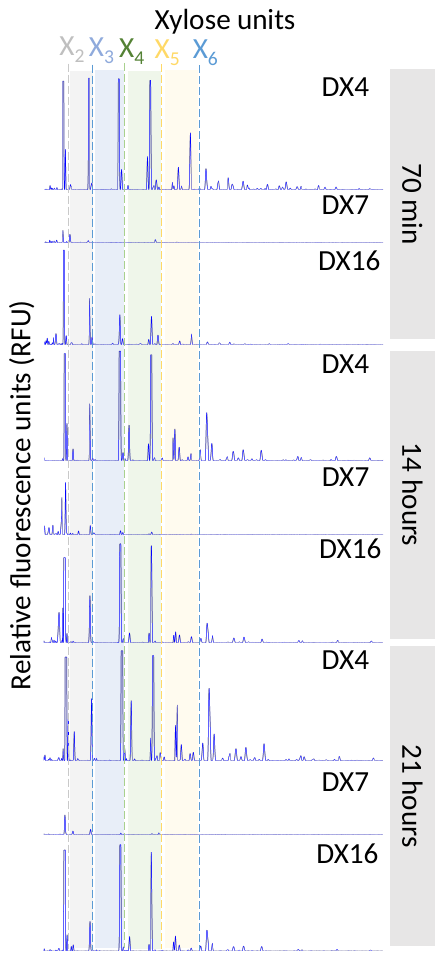

c)

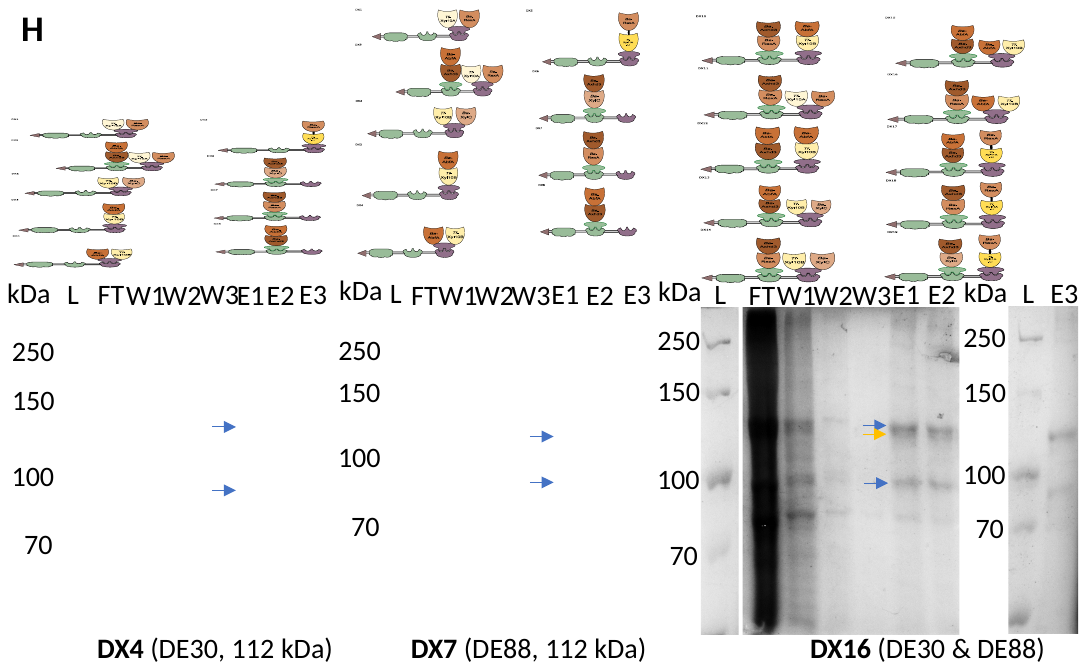

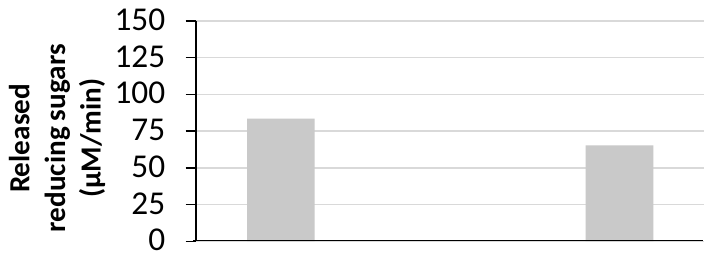

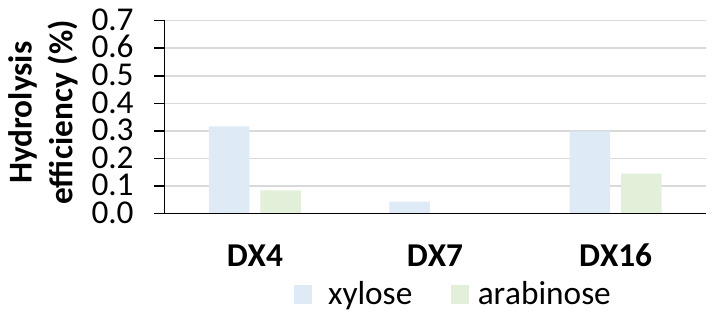

a)

b)

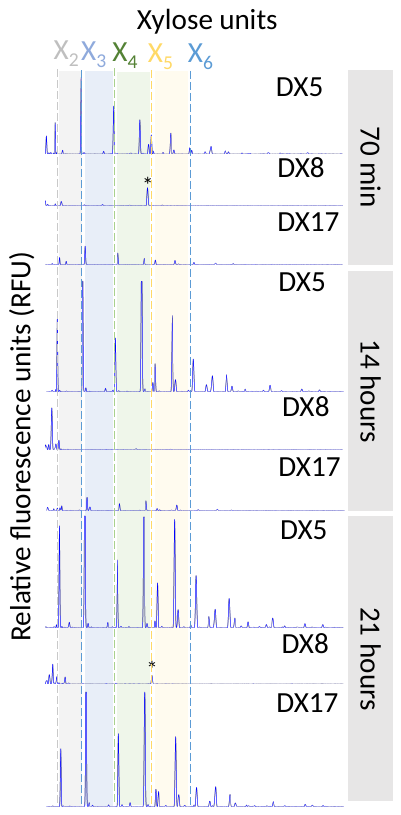

c)

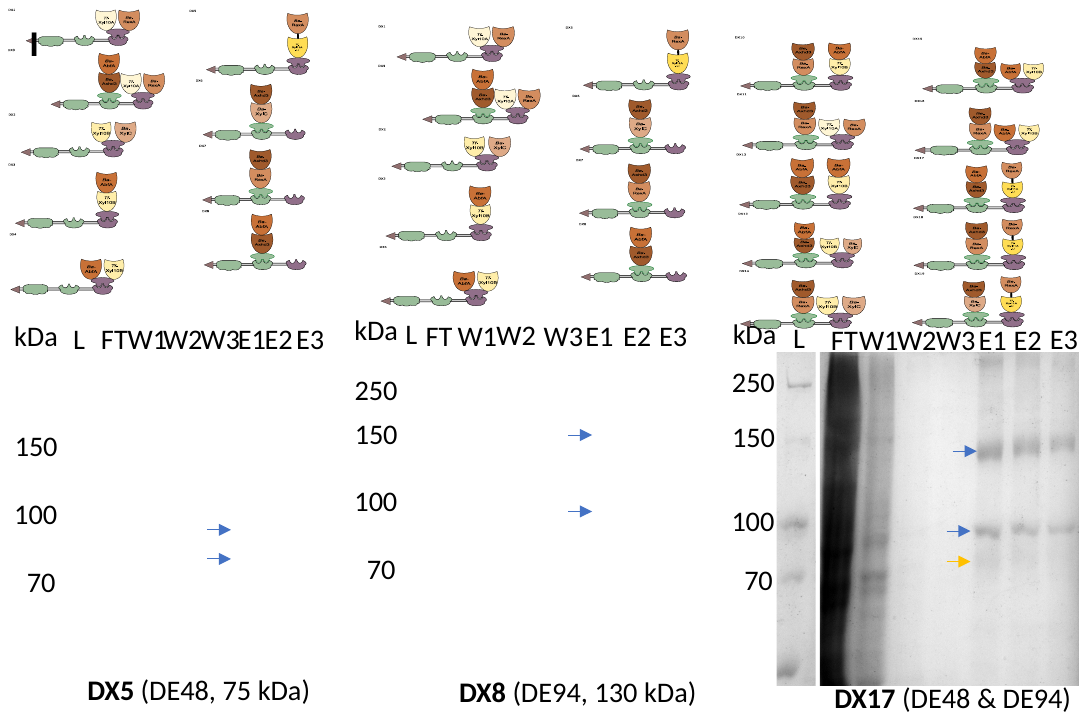

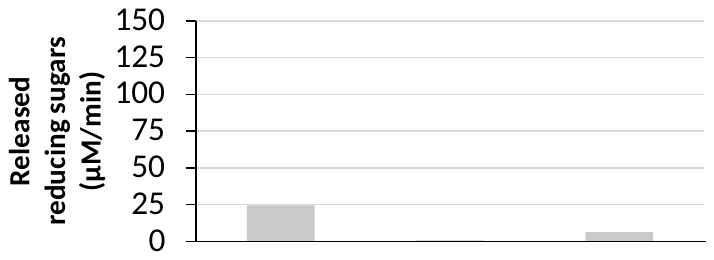

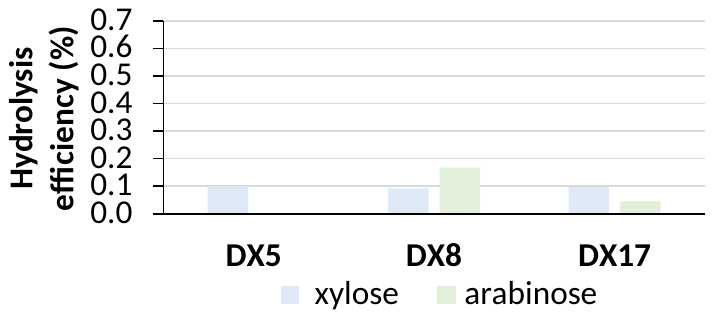

a)

b)

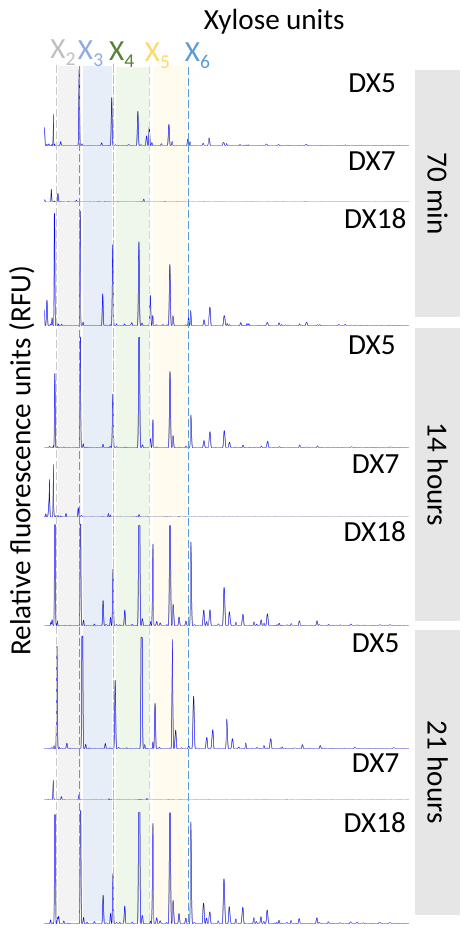

c)

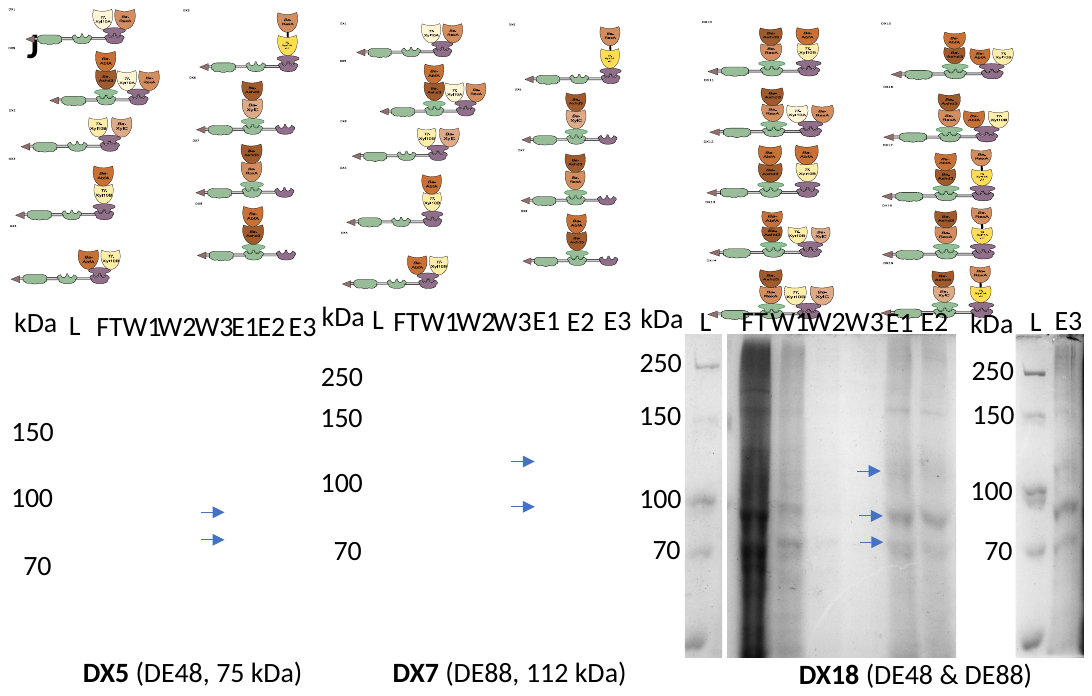

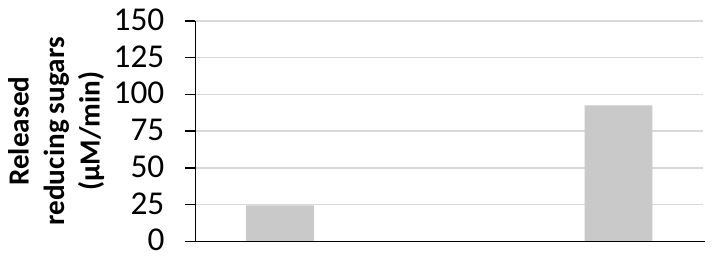

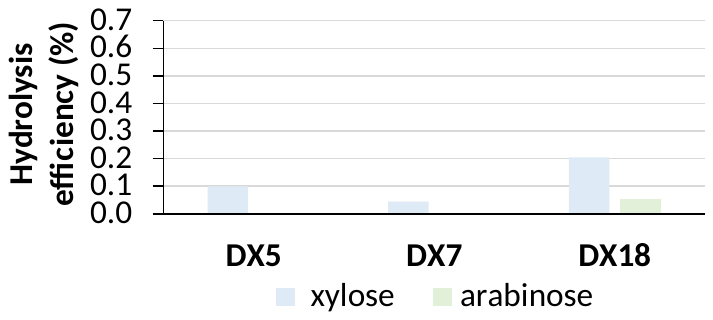

a)

b)

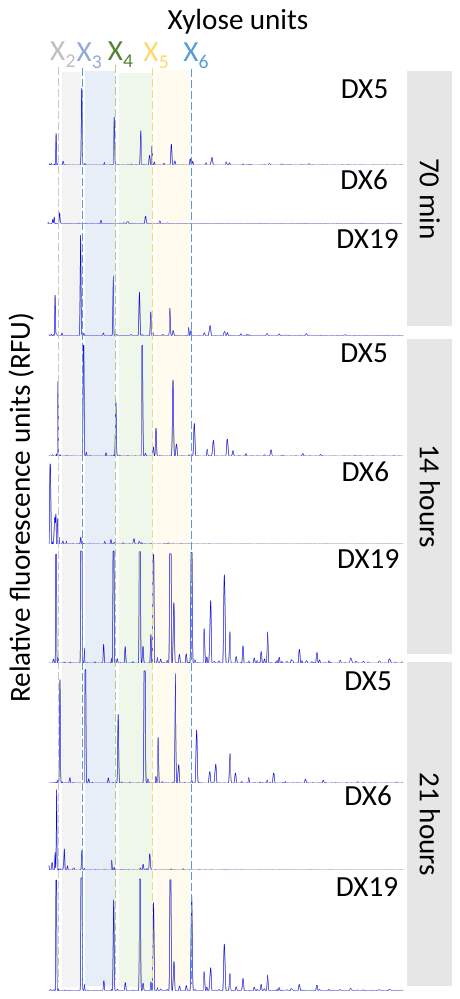

c)

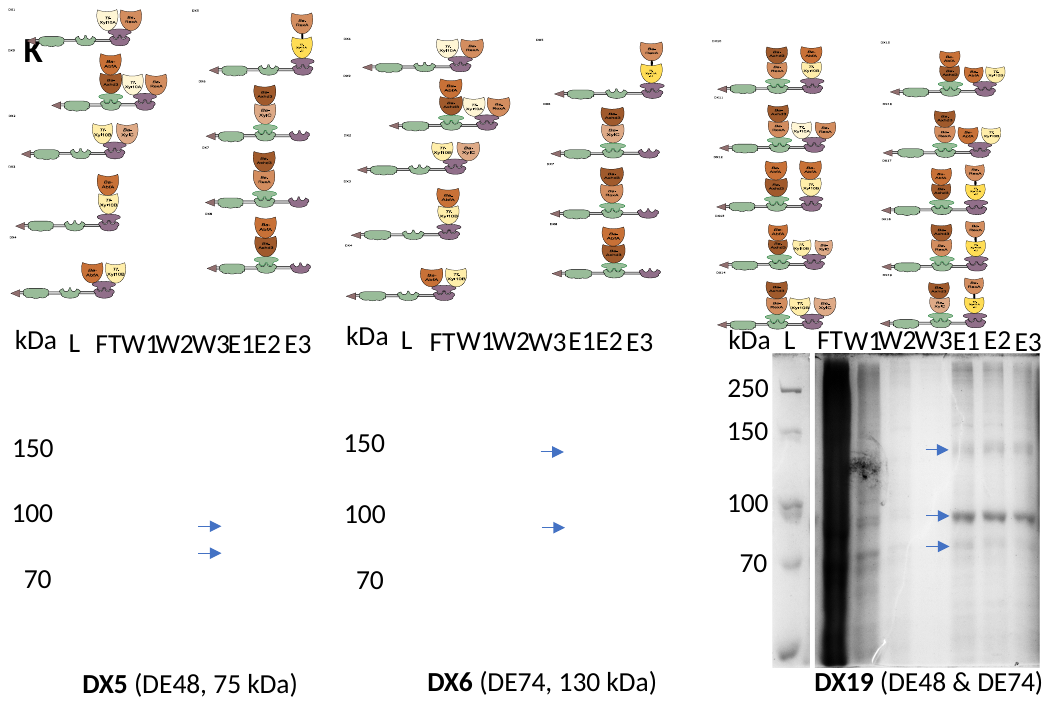

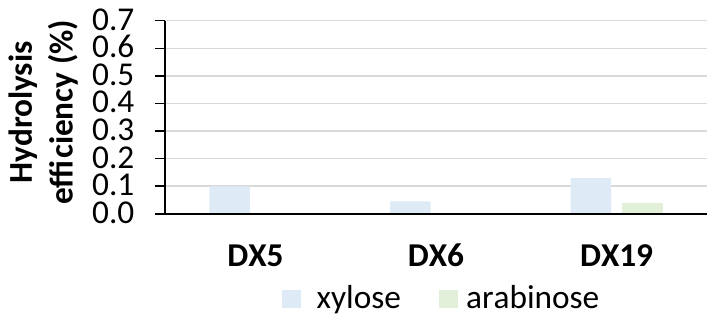

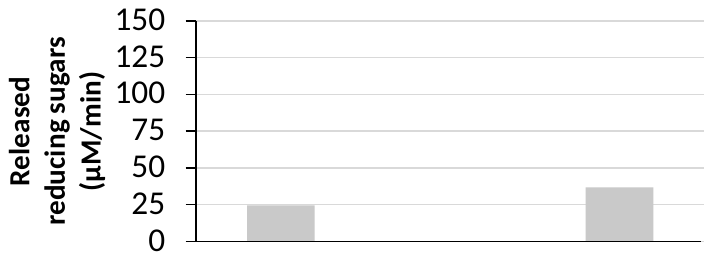

a)

b)

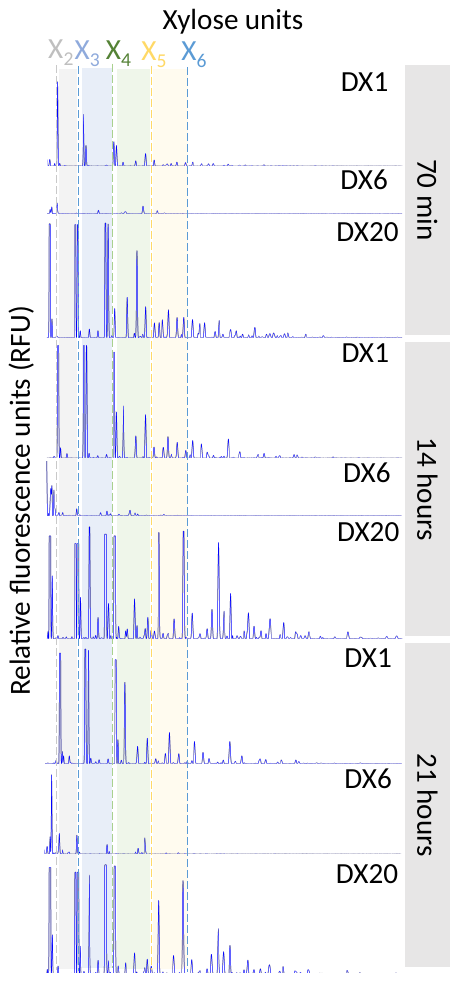

c)

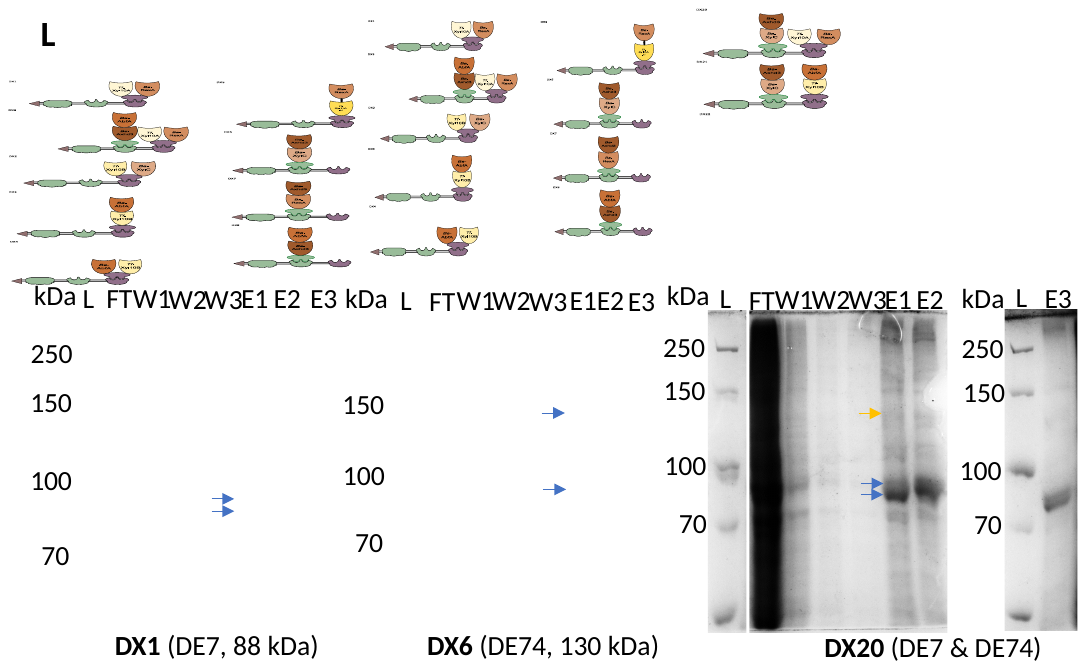

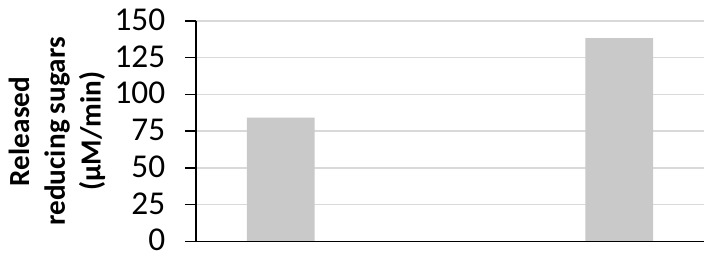

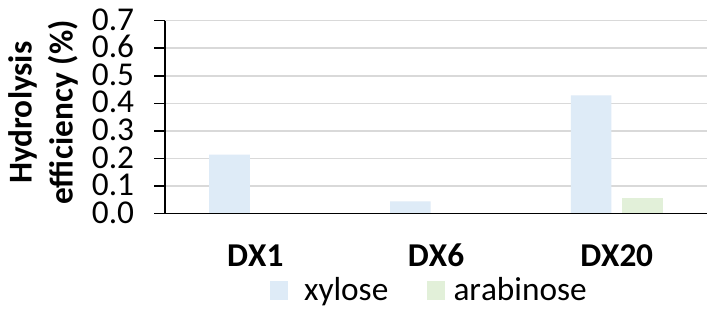

a)

b)

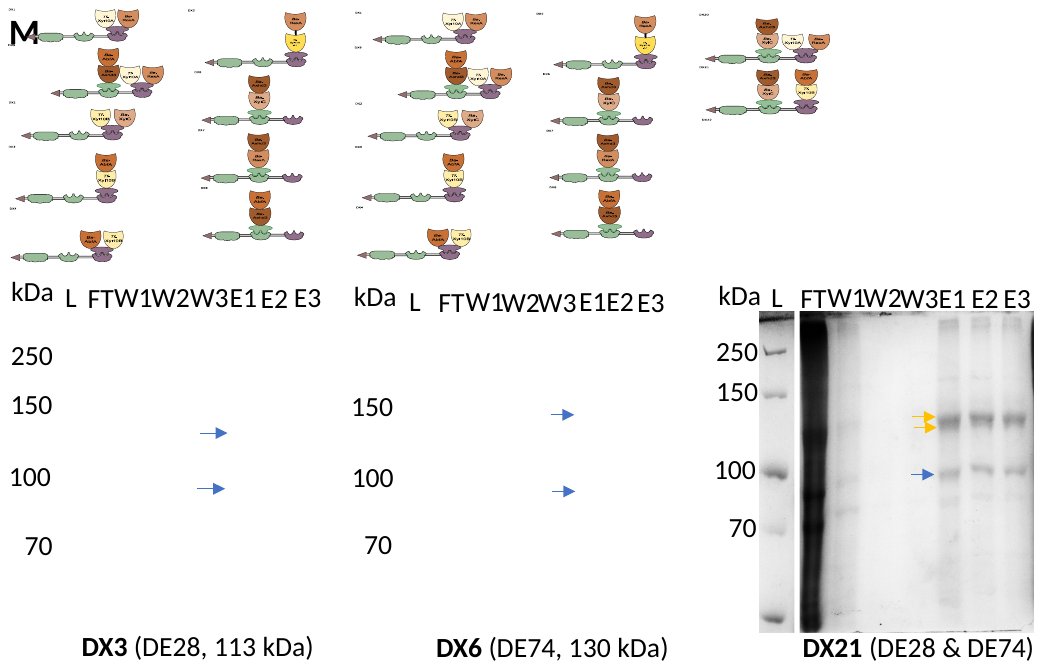

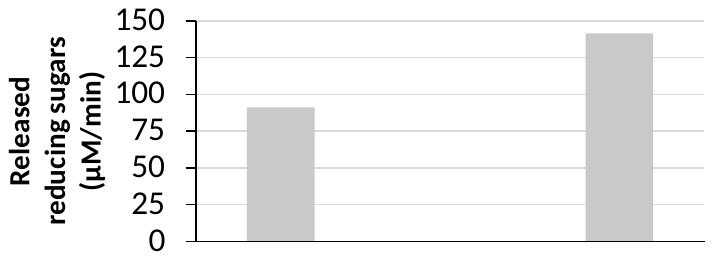

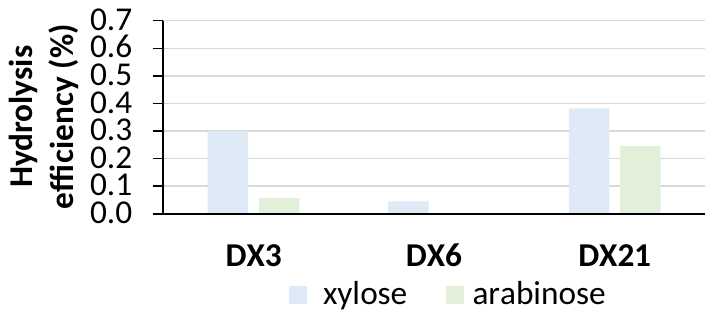

a)

b)

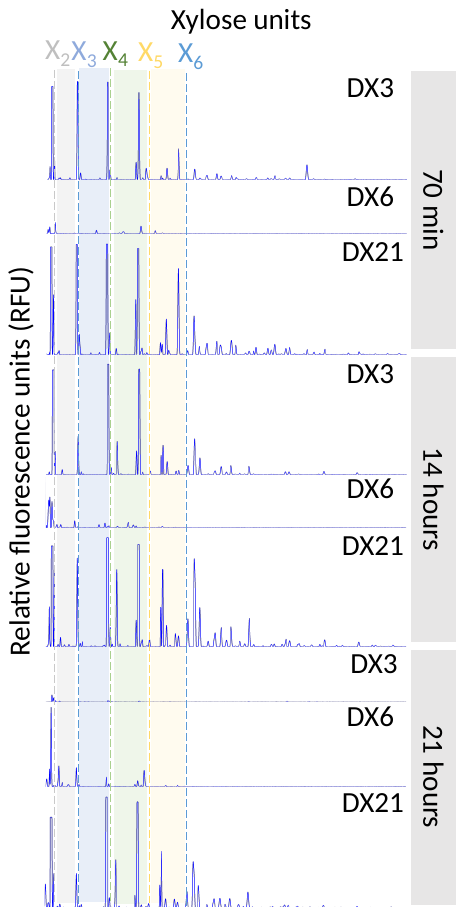

c)

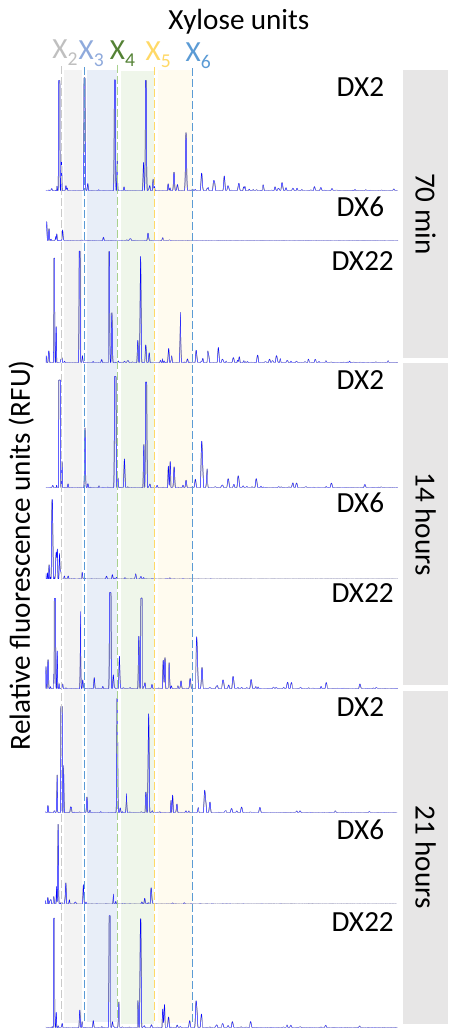

c)

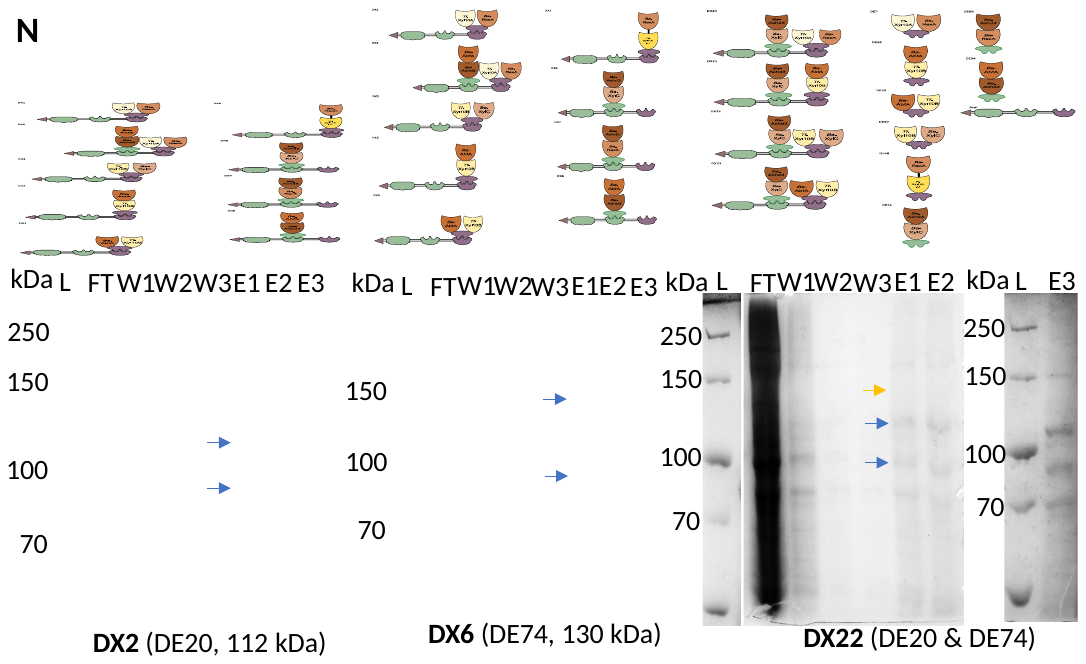

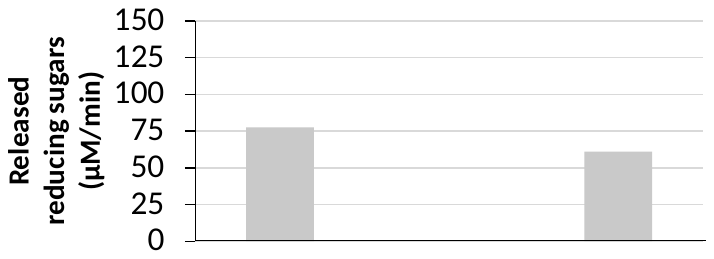

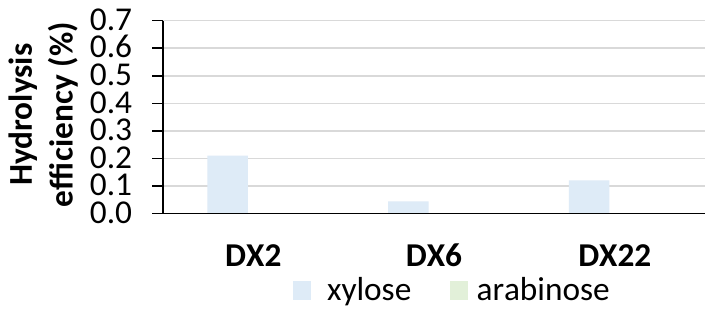

a)

b)

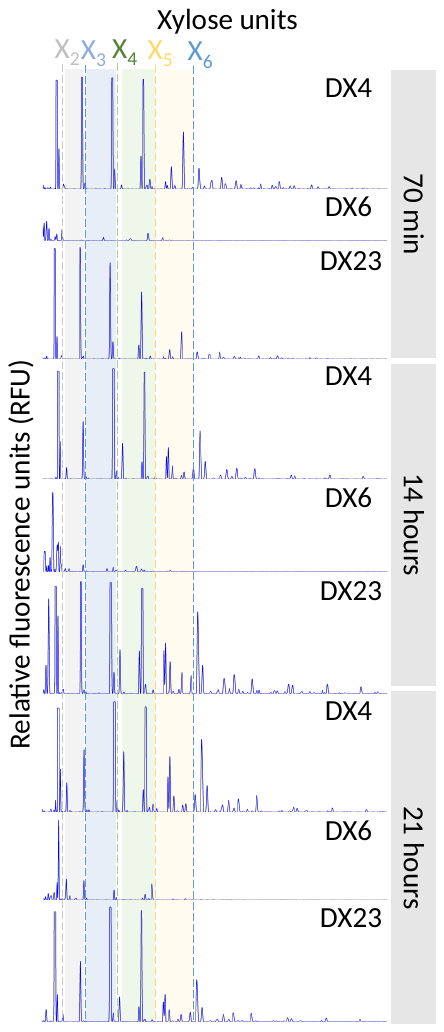

c)

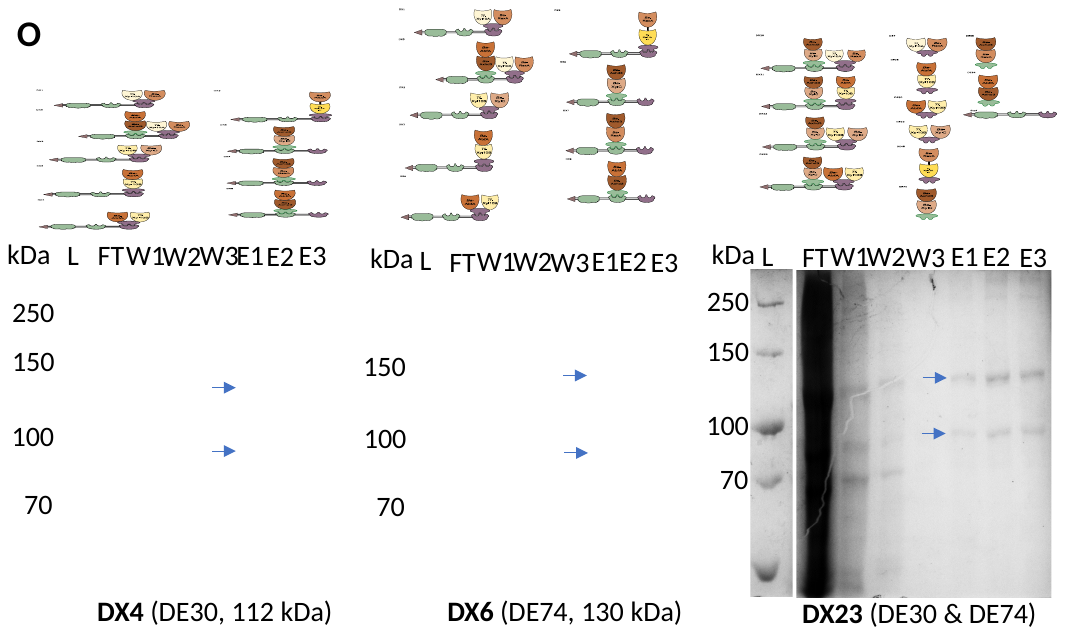

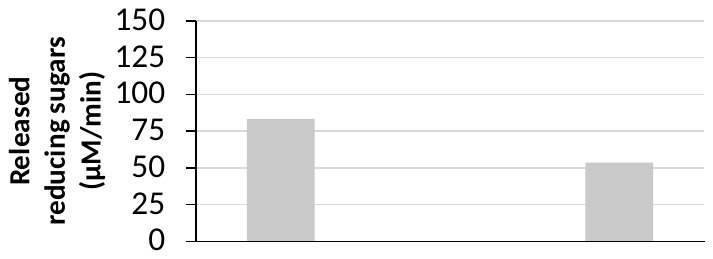

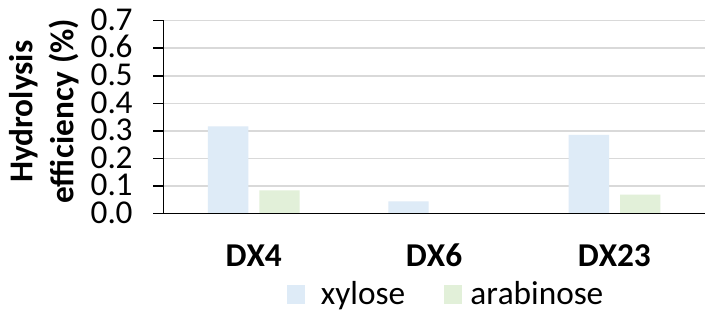

a)

b)

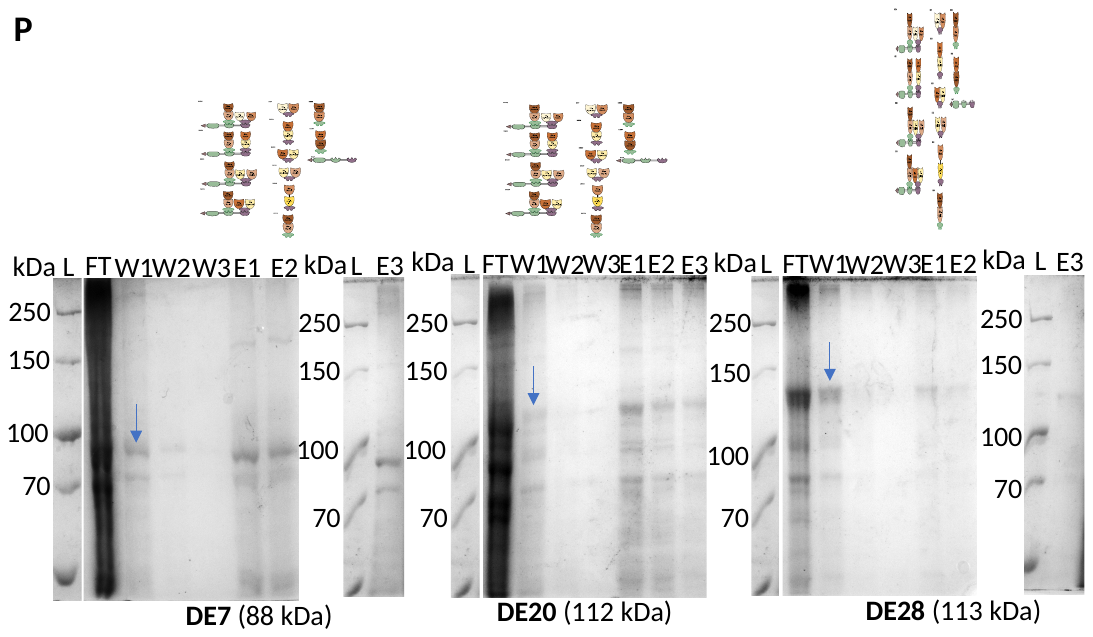

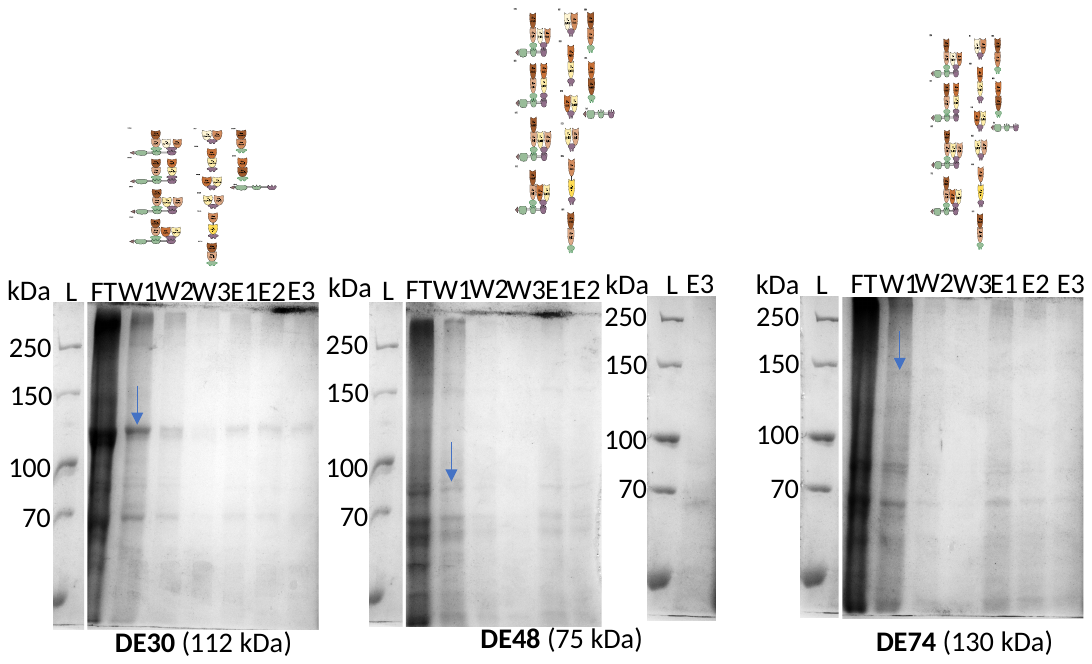

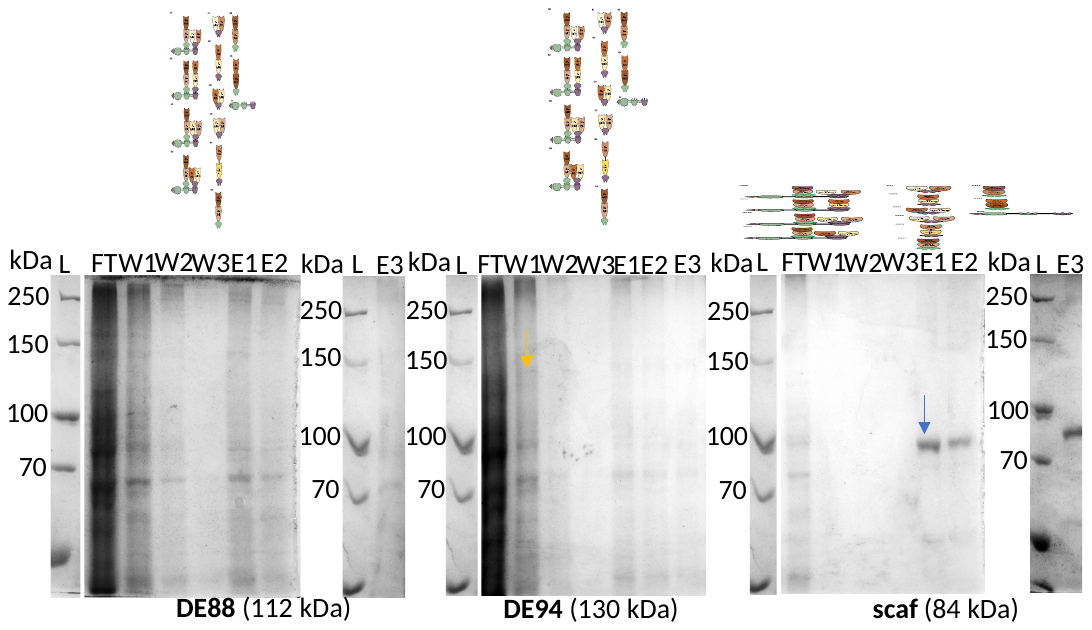

Rate of released reducing sugars and xylose and arabinose hydrolysis efficiency upon incubation of the newly assembled DXs with wheat flour arabinoxylan was checked. Panels A-O: fifteen DXs (DX9-DX23) containing the bivalent scaffoldin (Coh-*Ct*I−Coh-*Cc*) and two bicatalytic docking enzymes (DEs) (DE7 (*Tf*-Xyl10A−Dock-*Cc*−*Ba*-RexA), DE20 (*Tf*-Xyl10B−Dock-*Cc*−*Ba*-XylC), DE28 (Dock-*Cc*−*Tf*-Xyl10B−*Ba*-AbfA), DE30 (*Ba*-AbfA−Dock-*Cc*−*Tf*-Xyl10B), DE48 (Dock-*Cc*−*Tf*-Xyl11A-Li−*Ba*-RexA), DE74 (Dock-*Ct*I−*Ba*-XylC−*Ba*-Axhd3), DE88 (Dock-*Ct*I−*Ba*-RexA−*Ba*-Axhd3), (Dock-*Ct*I−*Ba*-Axhd3−*Ba*-AbfA)) were assembled in parallel. DXs containing the scaffoldin (Coh-*Ct*I−Coh-*Cc*) and one of the bicatalytic DEs (DX1-DX8) were also constructed to test the dockerin-cohesin affinity of each specific DE to the scaffoldin used. A) SDS-PAGE analysis of the fractions obtained from the GST-pull down assay was performed. The flow-through (FT), three wash (W1-3) and three elution (E1-3) fractions of GST pull-down assay were analyzed. Above each SDS-PAGE gel a schematic representation of the DE/DX tested is shown. Below the gels the code for each DE/DX tested is mentioned; and in the case of DXs, the DEs incorporated in the DX along with its molecular weight. The blue arrows point to the predicted DE and scaffoldin bands. The yellow arrows point to where one would expect the DE band to be seen but it is not clearly visible or absent. B) For each DX assembled the rate at which reducing sugars (checked with the DNS assay) are released is shown under the corresponding SDS-PAGE gel. The xylose and arabinose released by each DX was checked with the D-xylose and L-arabinose/D-galactose assay kits from Megazyme after 21 hours of reaction. The hydrolysis efficiency is calculated by dividing the amount of xylose/arabinose determined upon 21 hours of reaction by the maximum amount of xylose/arabinose that could be removed from the substrate used. C) Hydrolysates of each DX upon 70 min, 14 and 21 hours of reaction were analyzed by DSA-FACE. The dashed lines are the electrophoretic mobilities of the standards X_2_, X_3_, X_4_, X_5_ and X_6_. The regions highlighted in grey, blue, green and yellow correspond to electrophoretic mobilities of AXOS with DP3, DP4, DP5 and DP6, respectively. Panel P: the GST-pull down assay was also performed with DEs and scaffoldin alone to test the applicability of the GST-pull down assay in parallel.
